# A novel tubulin binding molecule drives differentiation of acute myeloid leukaemia cells

**DOI:** 10.1101/2021.06.03.446952

**Authors:** Thomas R. Jackson, Aini Vuorinen, Laia Josa-Culleré, Katrina S. Madden, Daniel Conole, Thomas J. Cogswell, Isabel V. L. Wilkinson, Laura M. Kettyle, Douzi Zhang, Alison O’Mahony, Deanne Gracias, Lorna McCall, Robert Westwood, Georg C. Terstappen, Stephen G. Davies, Edward W. Tate, Graham M. Wynne, Paresh Vyas, Angela J. Russell, Thomas A. Milne

## Abstract

Acute Myeloid Leukaemia (AML) continues to have a poor prognosis, especially in the elderly. One reason for this is that many treatment regimens are not well tolerated by elderly patients. Much current focus is on the development of therapies that can target specific vulnerabilities of AML while having fewer toxic side effects. However, despite much recent progress in developing better drugs, many patients with AML still die within a year of diagnosis, partly due to the fact that it is difficult to identify therapeutic targets that are effective across multiple AML subtypes. One common factor across AML subtypes is the presence of a block in differentiation. Thus screening for compounds that can overcome this block in genetically diverse AML models should allow for the identification of agents that are not dependent on a specific mutation for their efficacy. Here, we used a phenotypic screen to identify novel compounds that stimulate differentiation in several AML cell lines. Lead compounds were shown to decrease tumour burden and to increase survival *in vivo*. Using multiple complementary target deconvolution approaches, these compounds were revealed to be anti-mitotic tubulin disruptors that cause differentiation by inducing a G2-M mitotic arrest. Together, these results reveal a novel function for tubulin disruptors in causing differentiation of AML cells.

## Introduction

Acute Myeloid Leukaemia (AML) is a haematological malignancy with around 3,000 new cases per year in the UK ^1^ and about 21,000 new cases per year in the US ^2^. AML has a very poor survival rate, especially in elderly patients (5-year survival less than 11% ^3^) which represent the majority of cases ^1^. The current standard of care involves inducing remission using intensive chemotherapy, such as cytarabine in combination with anthracycline-derived antibiotics such as daunorubicin, followed by consolidation chemotherapy or bone marrow transplantation ^3–5^. Such therapies are not well tolerated by elderly patients and often have extensive side effects ^3^.

The promise of precision medicine is to develop targeted therapies that can specifically impact cancer cells while leaving normal cells unharmed, with the hope that such therapies will be effective and also have fewer toxic side effects. Characterization of a patient’s underlying mutational profile is becoming increasingly important for identifying patient subgroups that will be sensitive to specific targeted therapies. For example, patients that carry mutations in genes such as *fms like tyrosine kinase 3* (*FLT3*) or *isocitrate dehydrogenase 1 or 2* (*IDH1/2*) can be treated with small molecules designed to specifically target these mutations ^6, 7^. However, due to the high levels of heterogeneity among AML patients^8^, even with patients carrying the same driver mutation, often only a subset of patients respond well to targeted therapy, such as in the case of *IDH2* mutations and the drug enasidenib ^9–12^. In general, despite their promise, there has been a high degree of failure in clinical trials for AML which utilise among novel targeted therapies ^13^.

Another issue with targeted therapies is that many of them benefit small patient populations only, leaving the wide range of AML patients without effective treatments ^13^. The promise of immunotherapies ^14^ and recent exciting clinical trials with the B-cell lymphoma 2 (BCL-2) inhibitor venetoclax ^3, 15–17^ represent two approaches for targeting AML not limited to specific mutations. Even with these promising new areas of treatment, there remain patients who do not respond well to treatment, and much work is being done on possible combination therapies although this is often developed empirically without clear underlying mechanistic principles guiding the process ^3^. In all, there continues to be an urgent unmet need for new, well-tolerated therapies that can provide complete durable remission, especially for patient subsets that do not have a clear, well defined molecular target underlying the malignancy ^3, 13, 18^.

A defining hallmark of AML is a block in the normal myeloid differentiation process, blocking the production of downstream blood lineages and disrupting normal haematopoiesis. An exciting new paradigm in AML treatment is the possibility of inducing normal differentiation of AML cells by removing the differentiation block. Such therapies could be both more effective and less toxic than conventional chemotherapies, and may also provide effective partners for novel combination therapies. As an exemplar of such an approach, a breakthrough in differentiation therapy was achieved in the treatment of acute promyelocytic leukaemia (APL), which represents subset of about 10% of all AML patients ^19, 20^. APL is defined by a specific translocation involving the retinoic acid receptor to create a fusion oncoprotein ^21, 22^ which initially correlated with a poor prognosis. APL is now treatable with an 85% 5-year survival rate ^23^ due to the introduction of differentiation therapy with all-trans retinoic acid (ATRA); in combination with arsenic trioxide, a compound that causes degradation of the PML-retinoic acid receptor alpha oncogenic driver fusion protein ^24^. However, this therapy targets the specific oncoprotein which represents a vulnerability of APL not found in other AML subsets. Nonetheless, the success of this treatment suggests that the induction of differentiation by other mechanisms could provide novel treatments or new combination therapies for other subtypes of AML. Indeed, when AML patients carrying *IDH1/2* neomorphic mutations (mIDH, 15-25%) respond to treatment with the specific inhibitors ivosidenib (which targets mIDH1 ^25^), or enasidenib (which targets mIDH2 ^26^), they often display evidence of differentiation ^27, 28^.

These examples suggest that inducing differentiation in AML may be more effective than current treatments. However, in each of these examples, the drugs have been developed for highly specific targets which are not present in the majority of leukaemia patients. Recent work using an *in vitro* screening approach to identify novel inducers of differentiation resulted in the identification of a new class of dihydroorotate dehydrogenase (DHODH) inhibitors. In early preclinical work, DHODH inhibitors appear to be effective at inducing differentiation in AML cells in a non-mutation specific manner^29–31^. Although it remains unclear how DHODH inhibition directly induces differentiation in AML cells ^31^, this work provides a promising proof of principle that such screening approaches are an effective way of finding novel compounds for differentiation therapy. However, until the mechanism of action of such compounds is better understood, it is unclear exactly which patient subsets will respond to such a therapy, which could be part of the reason why recent clinical trials for a DHODH inhibitor were terminated due to lack of benefit^29^. Thus, there remains a further need for the identification of novel compounds and alternative mechanisms that can induce differentiation in AML cells in a mutation agnostic manner.

Here, we developed an *in vitro* flow cytometry-based phenotypic screen to identify new classes of small molecules which are capable of promoting differentiation in AML blasts, and validated their differentiation profiles using RNA-seq. As AML is a highly heterogeneous disease^32^, the phenotypic screen was performed using several AML cell lines to identify molecules whose efficacy was not limited to a particular genetic subtype. From the confirmed hits thus identified, a number of compound series were selected for further optimisation. The resulting compounds showed *in vivo* efficacy in reducing tumour burden in a subcutaneous model and displayed increased survival following oral dosing in an orthotopic xenograft model. Using a combination of RNA-seq, BioMAP analysis (an *in vitro* platform which uses primary human cells to test drug efficacy and toxicity) and chemoproteomics, tubulin beta chain was identified as a direct binding target of these compounds. Using other known, and structurally distinct tubulin binders, we showed that tubulin disruption causes mitotic arrest, and mitotic arrest results in initiation of differentiation, thus highlighting a novel mechanism of action and usage for the compounds we have identified.

## Results

### A phenotypic screen for differentiation in multiple AML cell lines identifies novel compounds

A screen was initiated using a library containing 1000 structurally-diverse, commerciallt available small molecules, in order to identify compounds that were capable of differentiating four AML cell lines (HL-60, OCI-AML3, THP-1, KG-1). Properties of these cell lines are summarised in Table 1, and together they represent approximately 30% of known AML mutations ^32^. CD11b is a known cell surface marker of differentiated myeloid cells ^33, 34^ and flow cytometry (FACs) analysis was used to quantify upregulation of CD11b expression after compound treatment. As positive controls, phorbol 12-myristate 13-acetate (PMA ^35^) was used to induce the differentiation of HL-60 and KG-1 (Fig. 1A, Supplementary Fig. 1), while tranylcypromine (TCP ^36^) was used as a positive control for THP-1 cells (Supplementary Fig. 1) and GS87 was used as a positive control for OCI-AML3 cells. Cells were treated with 10 μM of compound for 4 days. Compounds that upregulated CD11b more than 10% in at least 3 cell lines were considered potential hits and selected for further investigation. Using this criterion, we identified 44 positive hits (Fig. 1B, Supplementary Fig. 1). The structure and CD11b upregulation data of an example hit, OXS000275 **1**, is shown in Fig. 1C-D.

**Table 1:**
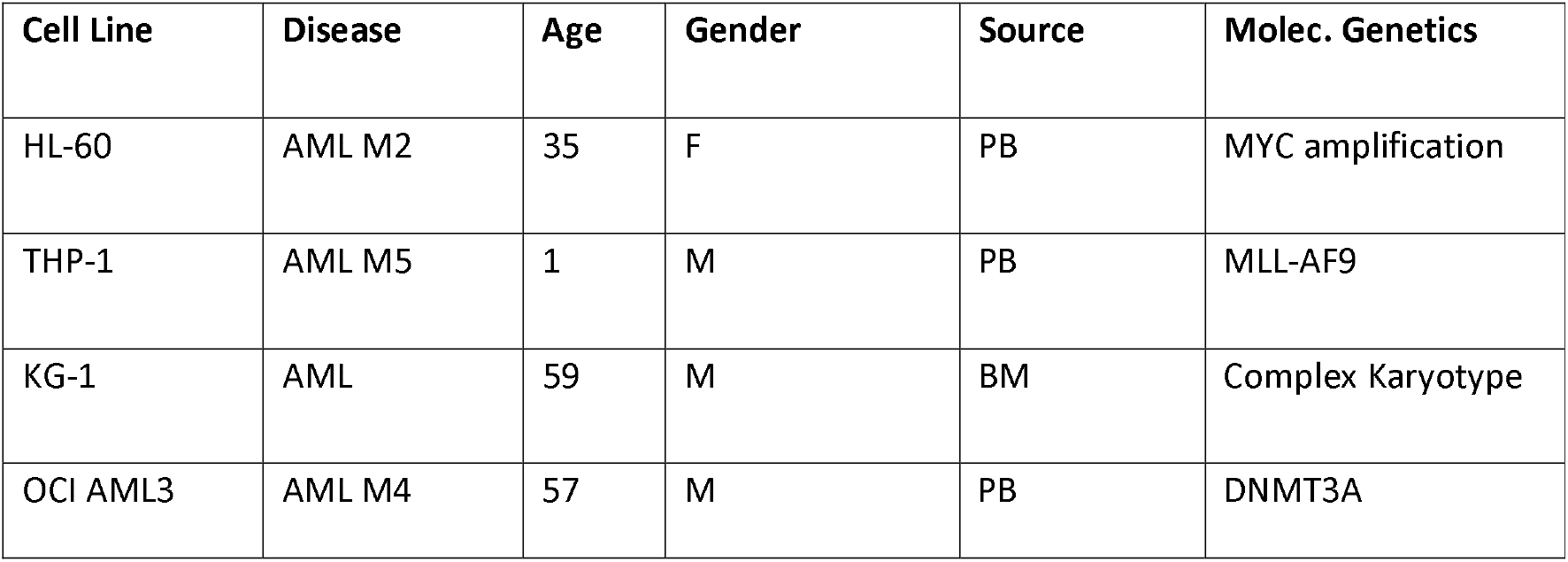
Cell line properties

| Cell Line | Disease | Age | Gender | Source | Molec. Genetics |
| --- | --- | --- | --- | --- | --- |
| HL-60 | AML M2 | 35 | F | PB | MYC amplification |
| THP-1 | AML M5 | 1 | M | PB | MLL-AF9 |
| KG-1 | AML | 59 | M | BM | Complex Karyotype |
| OCI AML3 | AML M4 | 57 | M | PB | DNMT3A |

**Figure 1:**
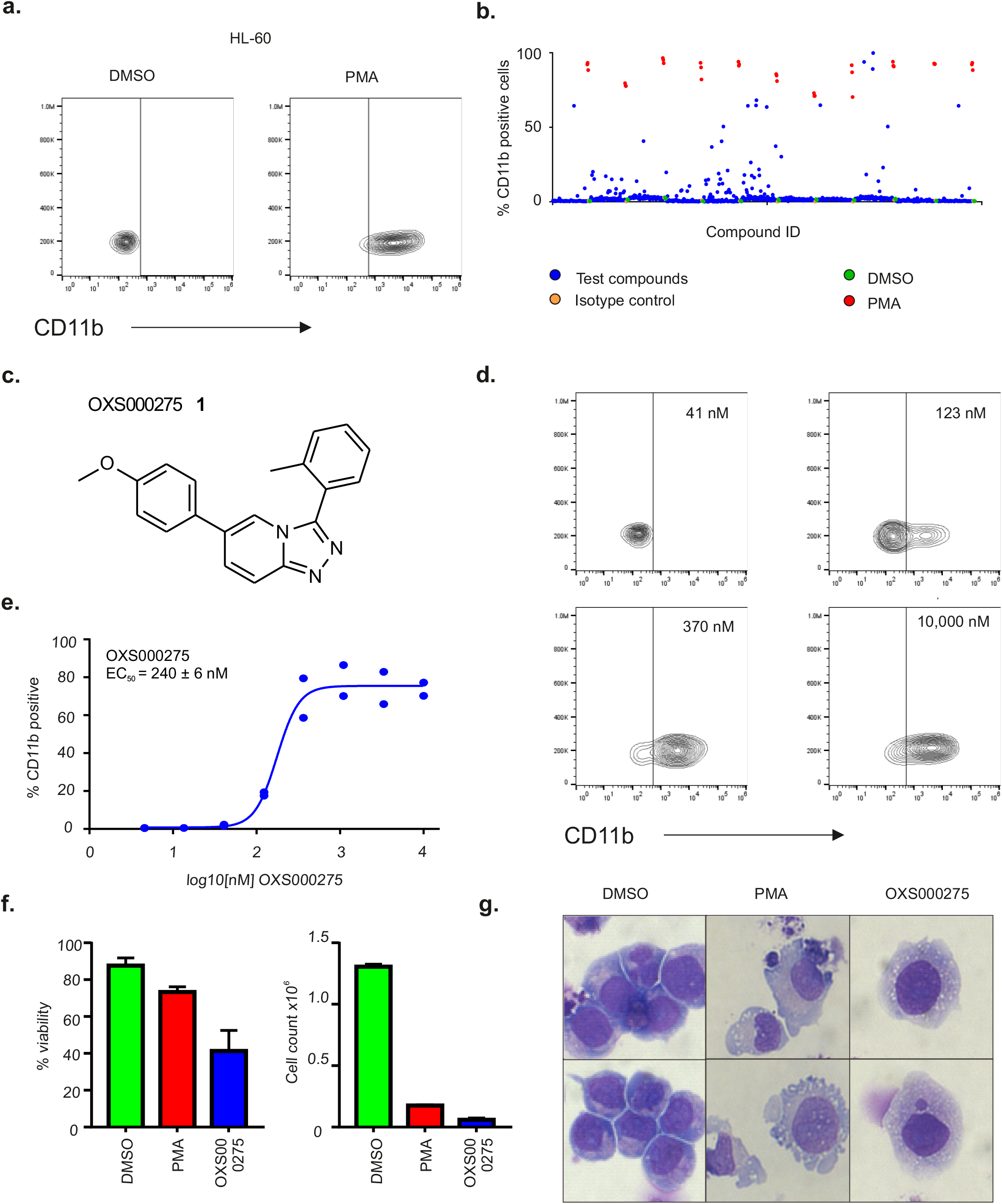
Phenotypic Screening and Validation of Primary Hits in HL-60 AML cell line. A) Upon treatment with positive control, PMA, HL-60 cells differentiate and up-regulate cell surface marker CD11b as detected by flow cytometry. B) Scatter plot distribution showing the results of HL-60 screening of 1000 compound library. C) Example of a biologically active compound identified (OXS000275 **1**). D) OXS000275 up-regulated CD11b at lower concentration as shown by flow cytometry. E) OXS000275 upregulated CD11b in a dose dependent manner with an EC_50_ of 238 ± 52 nM. F) At 4 days post treatment with OXS000275 a reduced viability and cell number was observed. G) Cytospin preparations of OX000275-treated or PMA-treated HL-60 cells stained with Wright-Giemsa showed signs of myeloid maturation.

### Novel identified compounds induce both neutrophil and macrophage differentiation in an AML cell line

Concentration-dependent responses were confirmed for hit compounds such as OXS000275, which was found to have a calculated EC_50_ of 240 ± 6 nM (Fig. 1D-E). Further effects of hit compounds on cell proliferation and viability were measured by staining for dead cells with DAPI and using acridine orange as a counterstain for all cells (Fig. 1F). Differentiation was also validated using morphology as characterised by Wright-Giemsa staining, with lighter cytoplasm, a higher cytoplasm to nuclei ratio and an increase in granulation used as signs of differentiation towards a macrophage phenotype (Fig. 1G). OXS000275 significantly inhibited cell proliferation and reduced cell viability (Fig. 1F) as well as promoting a morphology consistent with differentiation in all four of the cell lines (Fig. 1G, Supplementary Fig. 2).

To further confirm that the hits were inducing differentiation, selected compounds were further investigated using RNA-seq analysis, with PMA treatment used as a positive control for differentiation in HL-60 cells. Both PMA and OXS000275 caused genome wide changes in gene expression after 72 hours (Fig. 2A), but hierarchical clustering of genes showed that OXS000275 and PMA clustered separately from each other and from the DMSO control, demonstrating OXS000275 and PMA caused distinct gene expression profiles (Fig. 2B). However, there is a significant overlap between gene expression changes caused by OXS000275 and PMA, suggesting they both modulate common biological processes (Fig. 2B and C). Using principal component analysis (PCA), compound-treated HL-60 cells were compared to primary human cells of the myeloid lineage ^37^ (Fig. 2D). As expected, HL-60s treated with DMSO were found to cluster closer to stem and progenitor cell populations whereas cells treated with PMA and OXS000275 clustered closer to terminally differentiated monocyte populations (Fig. 2D). RNA-seq signatures were also analysed using EnrichR with ARCHS4 signatures, and both PMA and OXS000275 upregulated genes significantly overlapped with those of macrophages, but not stem or progenitor cells (Fig. 2E). Despite PMA inducing a larger number of differentially expressed genes, the macrophage signature in PMA treated cells was found to be less significant and produced a lower enrichment score than in OXS000275 treated cells (Fig. 2E). This result potentially reflects the promiscuity of PMA and its subsequent impact on a wide range of different biological processes. Besides producing a more specific macrophage signature compared to PMA, OXS000275 treatment also produced a significant neutrophil gene expression profile while PMA had little effect on the expression of neutrophil specific genes (Fig. 2E). Finally, gene set enrichment analysis (GSEA) analysis confirmed both PMA and OXS000275 signatures were enriched for macrophage genes, while OXS000275 was found to also be enriched for genes associated with neutrophils whereas PMA was not (Fig. 2F). Taken together these data suggested OXS000275 is able to induce global gene expression changes associated with differentiation at least to the same extent as PMA treatment. However, gene expression changes induced by OXS000275 treatment appeared to be more specific to differentiation than those induced by PMA. Finally, unlike PMA, OXS000275 treatment was able to induce upregulation of both macrophage and neutrophil associated RNA signatures. A summary of EnrichR analysis of other confirmed hits from the phenotypic screen can be found in Supplementary Table 1.

**Figure 2:**
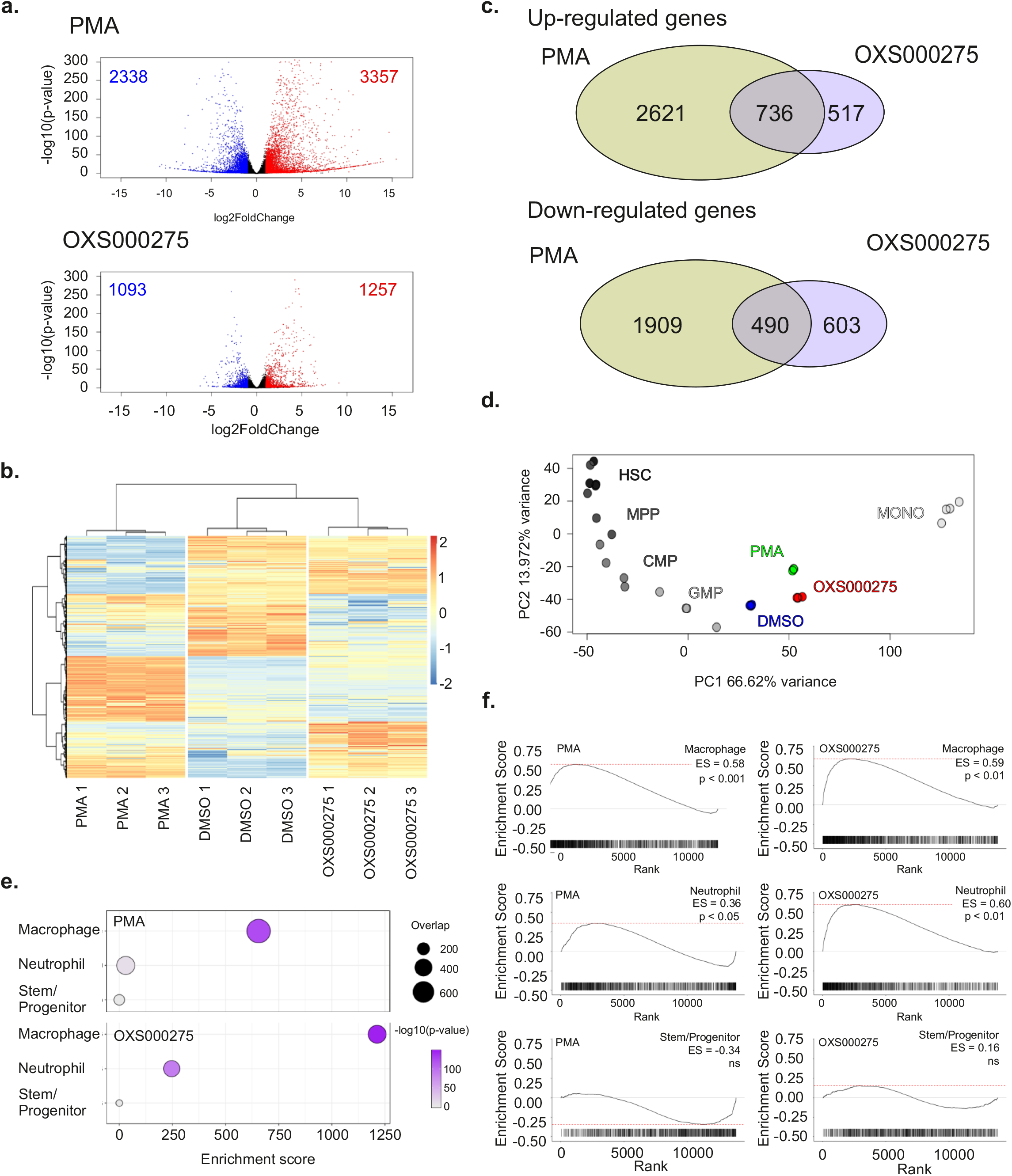
Confirmation of HL-60 Differentiation by Global Gene Expression Analysis. Cells were treated for 3 days with 10 nM of PMA, 1 μM of OXS000275 or 0.1% DMSO. A) Volcano plot of differentially expressed genes between DMSO control and PMA (top panel) and OXS000275. B) Venn diagram of differentially expressed genes post PMA and OXS000275 treatment. C) Heatmap of differentially expressed genes. D) Bulk RNA-sequencing of primary Hematopoietic cells from each population: PCA of each primary Hematopoietic cell population using the top 300 most varied expressed genes: HL-60 cells treated with vehicle (DMSO), PMA or OXS000275 projected onto plot. E) Treatment with OXS000275 and PMA lead to up regulation of genes consistent with myeloid differentiation when assessed by EnrichR and ARCHS4 Tissues signatures. P values were calculated using Fisher exact test, and Enrichment score as enrichment score = log(*p*) · *z*, *where* z is the z-score computed by assessing the deviation from the expected rank. F) Treatment with OXS000275 and PMA lead to gene-expression changes consistent with myeloid differentiation by gene set enrichment analysis.

### Development of Lead Compounds

Starting from confirmed hits, further optimisation afforded the lead compounds OXS007417 **2** and OXS007464 **3**, both of which had higher potency and improved ADME properties relative to the starting compound (Fig. 3A and B, detailed chemistry in ^38^). We confirmed that both OXS007417 **2** and OXS007464 **3** also caused differentiation of AML cell lines by showing that they upregulated CD11b cell surface expression in HL-60 cells with comparable EC_50_ values of 57 ± 3 nM and 36 ± 1 nM respectively (Fig. 3B). Differentiation was further confirmed by morphology as previously described (Fig. 3C).

**Figure 3:**
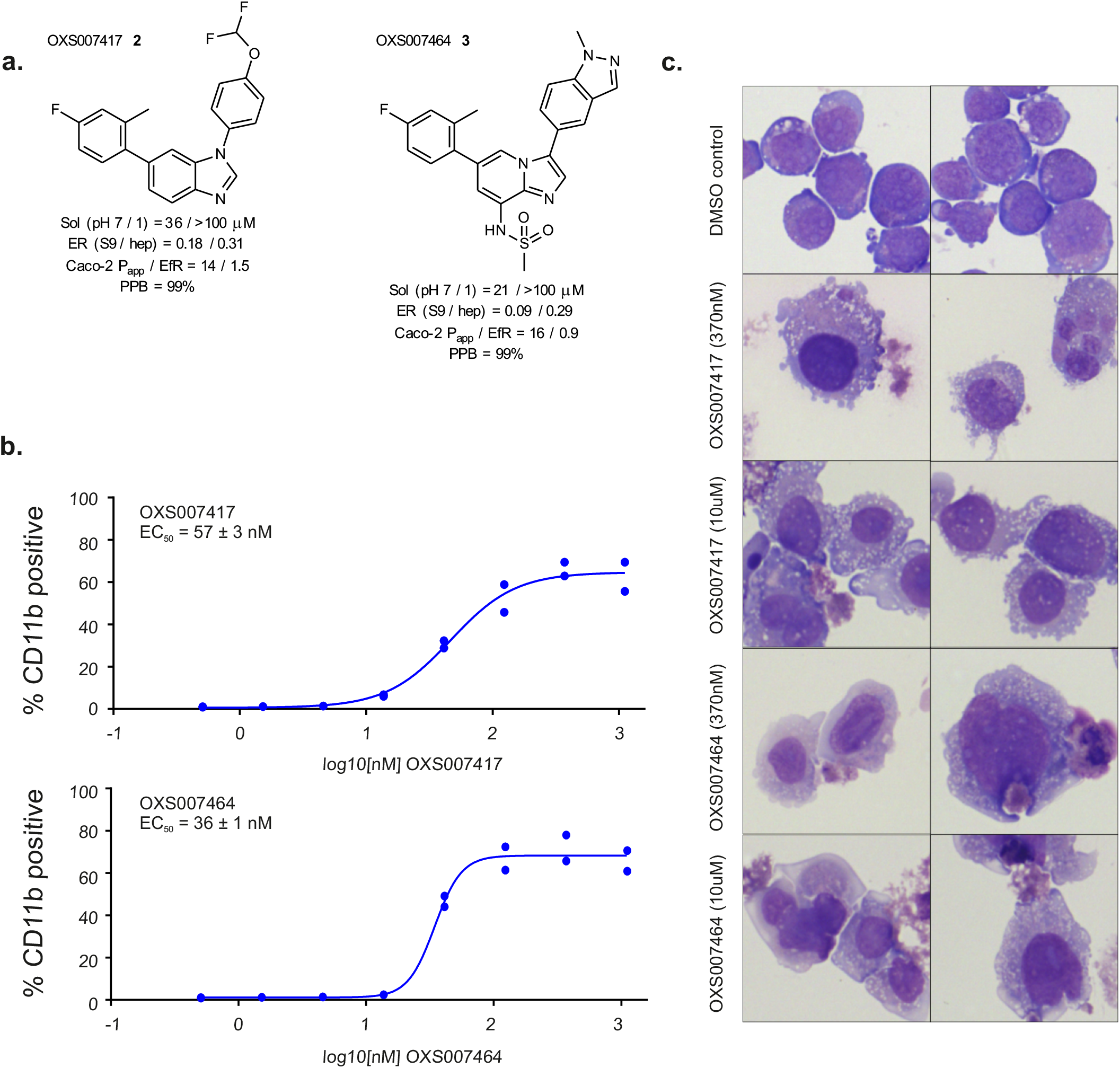
Development of lead compounds. A) Lead compounds OXS007417 **2** and OXS007464 **3** were developed from original hits. B) Lead compounds up-regulated CD11b in HL-60 cells by flow cytometry. C) Cytospin preparations of lead compound-treated HL-60 cells stained with Wright-Giemsa showed signs of myeloid maturation.

### Lead Compounds Demonstrate Anti-leukaemia Activity *In Vivo* in a subcutaneous Xenograft Model of AML

To study the ability of OXS007417 **2** and OXS007464 **3** to inhibit tumour growth *in vivo*, a subcutaneous xenograft model was used in the first instance and compared to standard chemotherapeutic agents. HL-60 cells were implanted into the flank of female NOD SCID mice, and tumours were allowed to reach a volume of 150 mm^3^ before commencement of treatment (Fig. 4A). OXS007417 and OXS007464 were administered per os (PO) twice daily for 4 weeks at 10 and 3 mg/kg respectively, while cytarabine (araC) was used as a reference of the standard of care (SoC) for AML^3^ and administered via intraperitoneal (IP) injection, 20 mg/kg once daily ^39^. Finally, ATRA (PO, 5 mg/kg 5 on/2 off) was used as reference differentiating agent. Treatments are summarised in Fig. 4A and Table 2.

**Figure 4:**
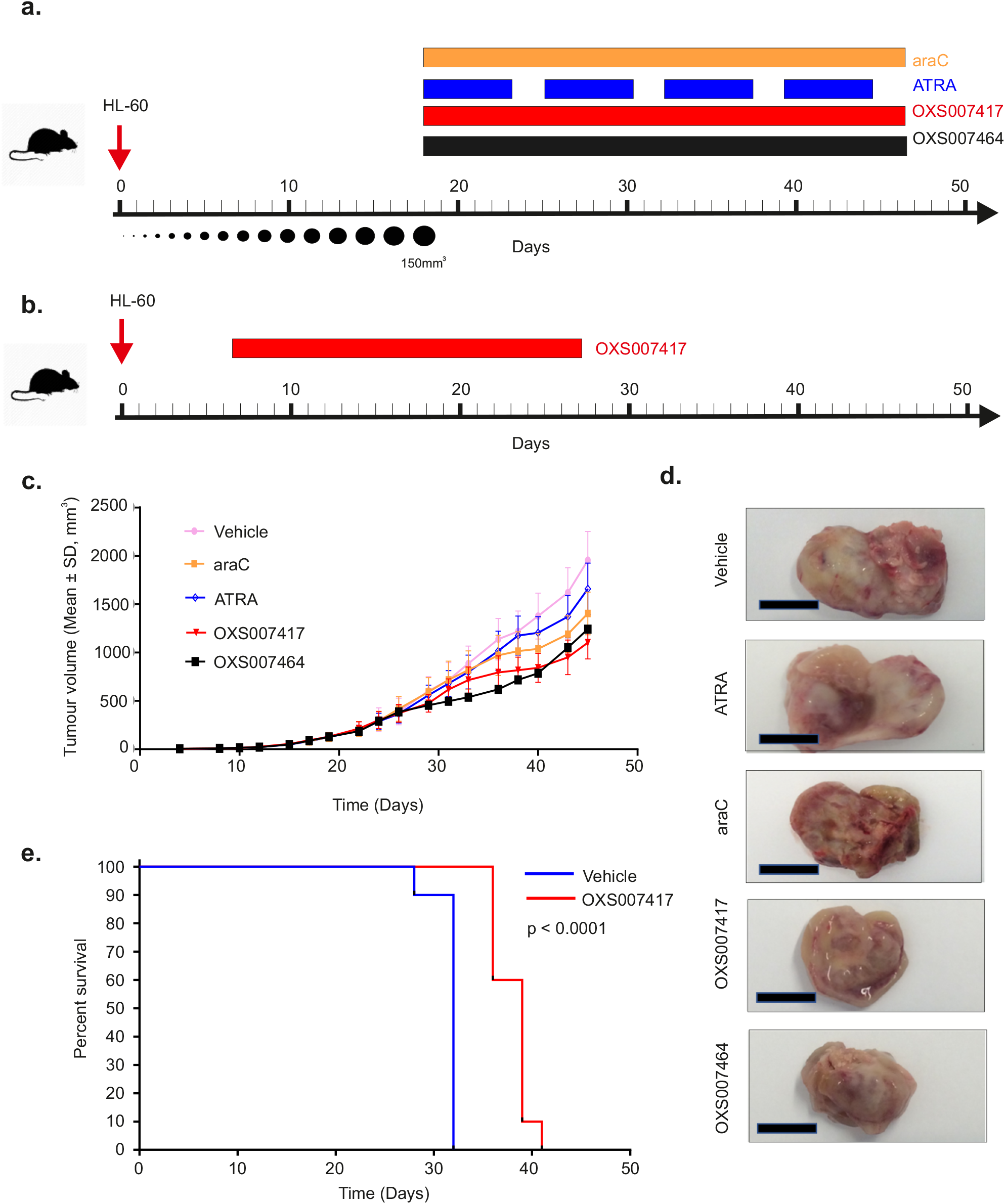
Lead Compounds Demonstrate Anti-leukemia Activity *In Vivo* in Subcutaneous Xenograft Model and Increased Survival in Orthotropic Model. A) Experimental outline of HL-60 subcutaneous xenograft model. B) Experimental outline of orthotropic model. C) HL-60 cells were implanted subcutaneously onto the flank of female NOD SCID mice, and the mice were treated with vehicle or indicated compounds; treatment with OXS007417 and OXS007464 reduced tumour growth. D) Example of excised tumours at termination of study; scale bar = 10 mm. E) OXS007417 prolonged the survival in an orthotropic HL-60 model.

**Table 2:** Treatment regimens for subcutaneous model

| Group | n | Treatment | Dose Level | Dosing Route | Dosing Regimen | Formulation | Dose volume |
| --- | --- | --- | --- | --- | --- | --- | --- |
| 1 | 10 | Vehicle | -- | PO | BID | 5% DMSO and 95% PBS + 0.1% Tween 20 | 10 ml / kg |
| 2 | 10 | AraC | 20 mg/kg | IP | QD | sterile water | 10 ml / kg |
| 3 | 10 | ATRA | 5 mg/kg | PO | 5 on/2 off | ethanol 10%; CMC-NA 90% | 10 ml / kg |
| 4 | 10 | OXS007417 | 10 mg/kg | PO | BID | 5% DMSO and 95% PBS + 0.1% Tween 20 | 10 ml / kg |
| 5 | 10 | OXS007464 | 3 mg/kg | PO | BID | 5% DMSO and 95% PBS + 0.1% Tween 20 | 10 ml / kg |

Treatment with OXS007417 was well-tolerated and did not lead to significant body weight loss in the animals (Supplementary Fig. 3). After 28 days of dosing, OXS007417 significantly delayed the growth of HL-60 tumours, with a tumour control ratio (T/C) of 55%, when compared to vehicle group (p<0.0001), (Fig. 4C and D).

The standard of care araC showed a less pronounced effect, with T/C of 70% (p<0.0001). The smallest effect was observed in the group treated with ATRA, which did not show a significant (p = 0.07) reduction of tumour volume.

At conclusion of the study, plasma and tumour samples were taken for bioanalysis. In animals treated with OXS007417, compound exposure was detected in both samples, at 6.84 x 10^−7^ M (252 ng/mL plasma) and 2.41 x 10^−9^ mol/g (888 ng/g tumour) respectively. In animals treated with OXS007464, compound exposure was detected in both samples, at 3.38 x 10^−7^ M (152 ng/mL plasma) and 4.89 x 10^−10^ mol/g (220 ng/g tumour) respectively. OXS007464 was not as well-tolerated as OXS007417 and the dosage had to be reduced with some mice showing weight loss (Supplementary Fig. 3) and was therefore not used in further *in vivo* studies. The reason for this differing tolerance is unclear.

### OXS007417 causes increased survival using an *in vivo* murine orthotopic Xenograft Model of AML

Having shown *in vitro* to *in vivo* correlation using the subcutaneous HL-60 based model, the anti-leukaemia activity of OXS007417 was also evaluated in an orthotopic AML model. HL-60 cells were injected into the tail vein of female NCG mice, and 7 days were allowed for engraftment. Animals were then dosed with OXS007417 bid (formulated in 5% DMSO : 0.1% Tween 20 in PBS; PO; 10 mg/kg, with a dosing volume of 10 ml/kg) for 3 weeks (Fig. 4B). Significantly, OXS007417 produced prolonged survival (p<0.0001) compared with the vehicle control group (Fig. 4E).

### Analysis from the BioMAP Phenotypic Platform suggests tubulin disruption as a MoA for OXS007417 and OXS007464

With compounds in hand displaying *in vitro* differentiation properties and *in vivo* efficacy in two different tumour models, we next set out to identify the biological targets of the compounds and to identify possible mechanisms of action (MoA) for their activity. The Diversity PLUS panel (BioMAP^®^, Eurofins Discovery) is an *in vitro* platform which uses different primary human cell types to generate activity profiles designed to aid MoA studies and the identification of off-target effects as well as potential toxicity issues. The method utilises 12 primary cell-based systems modeling a broad scope of human tissue and disease biology and 148 protein biomarkers to create a “biomarker signature profile” that can be compared to a database of mechanistic signatures for >4600 compounds with known MoAs. The BioMAP study found OXS007417 and OXS007464 to be anti-proliferative to human primary B, T cells, coronary artery smooth muscle cells, endothelial cells, and fibroblasts (Fig. 5A), but not cytotoxic at the four concentrations tested (12 – 800 nM). Comparing the biological activities of OXS007417 and OXS007464 to those of known bioactive agents in the BioMAP reference database, OXS007417 and OXS007464 were found to have profiles most similar to microtubule disruptors (Table 3 and 4).

**Figure 5:**
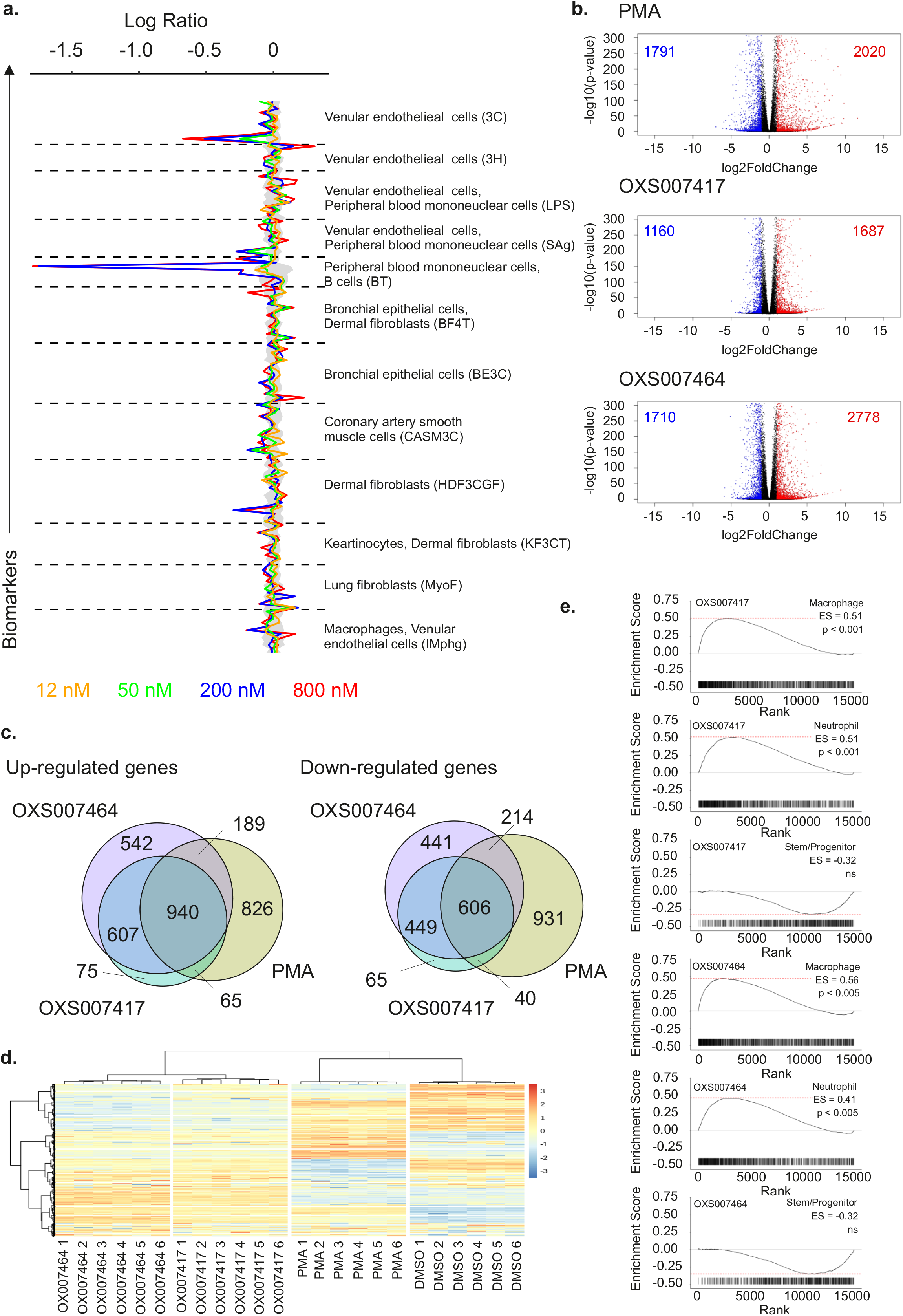
Elucidation of Lead Compounds Mechanisms of Action. A) BioMAP analysis of OXS007417 and OXS007464. Primary cell systems created form pooled donors were treated at four indicated concentrations. Cell system specific readouts were taken at time points optimised for each cell system. Readings from treated samples were divided by the average of control readings to generate a ration that was the log_10_ transformed. Significance prediction envelopes (grey) were calculated from control data at 95% confidence intervals. B) Volcano plot of differentially expressed between DMSO control and PMA (top panel), OXS007417 and OXS007464. C) Venn diagram of differentially expressed genes post PMA, OXS007417 and OXS007464 treatment. D) Heatmap of differentially expressed genes in six biological replicates treated with either OXS007417, OXS007464, PMA (as a positive control for differentiation) or DMSO (solvent only control). E) Treatment with OXS007417 and OXS007464 lead to gene-expression changes consistent with myeloid differentiation by gene set enrichment analysis.

**Table 3:** Top 3 similarity matches from an unsupervised search of the BioMAP Reference Database of > 4,000 agents for each concentration of OXS007464. The similarity between agents is determined using a combinatorial approach that accounts for the characteristics of BioMAP profiles by filtering (Tanimoto metric) and ranking (BioMAP Z-Standard) the Pearson’s correlation coefficient between two profiles. Profiles are identified as having mechanistically relevant similarity if the Pearson’s correlation coefficient is ≥ 0.7.

| Dose | Database match | BioMap Z-standard | Pearson's Score | # of common biomarkers | Mechanism Class |
| --- | --- | --- | --- | --- | --- |
| 800 nM | Colchicine 1.1 μM | 17.479 | 0.896 | 148 | Microtubule Disruptor |
| | Fosbretabulin Disodium 1.1 μM | 17.247 | 8.892 | 148 | Microtubule Disruptor |
| | Fosbretabulin Disodium 3.3 μM | 17.157 | 0.891 | 148 | Microtubule Disruptor |
| 200 nM | Fosbretabulin Disodium 3.3 μM | 19.495 | 0.924 | 148 | Microtubule Disruptor |
| | Fosbretabulin Disodium 10 μM | 19.257 | 0.922 | 147 | Microtubule Disruptor |
| | Fosbretabulin Disodium 30 μM | 19.188 | 0.921 | 148 | Microtubule Disruptor |
| 50 nM | GSK46136A, 370nM | 16.977 | 0.877 | 148 | PLK1 inhibitor |
|  | Vincristine Sulfate, 14 nM | 16.926 | 0.877 | 148 | Microtubule Disruptor |
|  | Pironection, 14 nM | 16.486 | 0.881 | 146 | Microtubule Disruptor |
| 12 nM | Erastin, 370nM | 5.678 | 0.496 | 112 | VDAC2 Blocker |
| | SR-2640, 30 μM | 4.851 | 0.392 | 140 | Leukotriene |
| | Bemegride, 32 μM | 4.364 | 0.428 | 94 | GABA-A Receptor Antagonist |

**Table 4:** Top 3 similarity matches from an unsupervised search of the BioMAP Reference Database of > 4,000 agents for each concentration of OXS007417. The similarity between agents is determined using a combinatorial approach that accounts for the characteristics of BioMAP profiles by filtering (Tanimoto metric) and ranking (BioMAP Z-Standard) the Pearson’s correlation coefficient between two profiles. Profiles are identified as having mechanistically relevant similarity if the Pearson’s correlation coefficient is ≥ 0.7.

| Dose | Database match | BioMap Z-standard | Pearson's Score | # of common biomarkers | Mechanism Class |
| --- | --- | --- | --- | --- | --- |
| 800 nM | Pironetin, 14 nM | 18.362 | 0.911 | 146 | Microtubule Disruptor |
|  | Pironetin, 41 nM | 17.424 | 0.897 | 146 | Microtubule Disruptor |
|  | Epothilone B, 37 nM | 17.402 | 0.895 | 148 | Microtubule Disruptor |
| 200 nM | Pironetin, 14 nM | 21.666 | 0.948 | 146 | Microtubule Disruptor |
|  | Epothilone B, 37 nM | 20.508 | 0.936 | 148 | Microtubule Disruptor |
|  | Epothilone B, 12 nM | 19.731 | 0.927 | 148 | Microtubule Disruptor |
| 50 nM | Paroxetine Hydrochloride, 14 μM | 7.380 | 0.639 | 98 | SERT Antagonist |
| | Nifedipine, 14 μM | 6.934 | 0.612 | 98 | L-type Ca<sup>++</sup> Channel Antagonist |
| | TMP-153, 370 μM | 6.926 | 0.519 | 148 | ACAT Inhibitor |
| 12 nM | Droperidol, 14 nM | 4.935 | 0.388 | 148 | Dopamine R Antagonist |
|  | Fludrortisone, 14 nM | 4.739 | 0.376 | 147 | GR Agonist |
|  | Sunitinib Malate, 14nM | 4.577 | 0.363 | 148 | VEGFR2 Inhibitor |

### Analysis by early RNA-seq suggests tubulin disruption as a MoA for OXS007417 and OXS007464

To further investigate the MoA of our novel compounds, HL-60 cells were treated for 6 hours with OXS007417 **2** (1 μM), OXS007464 **3** (1 μM), PMA (10 nM) or DMSO control before global gene expression changes were assessed by RNA-seq (Fig. 5 B-E). Even at this early time point, OXS007417, OXS007464 and PMA induced genome wide changes in gene expression (Fig. 5 B-D).

As with OXS000275, hierarchical clustering of genes showed that both OXS007417 and OXS007464 had distinct gene expression profiles from PMA and DMSO vehicle control (Fig. 5D). Significant overlap of differentially expressed genes with PMA again suggested the activation of common biological processes (Fig. 5C). Although forming their own distinct clusters, OXS007417 and OXS007464 clustered together with very similar gene expression profiles compared to each other, relative to PMA and DMSO treatment (Fig. 5D). The similarity between OXS007417 and OXS007464 gene signatures is further highlighted by the almost complete overlap of differentially expressed genes caused by these two compounds (Fig. 5C). The similarity between OXS007417 and OXS007464 at this early time point suggests a common mechanism of action is shared by these two compounds.

The L1000CDS2 database ^40^ was used to search for substances that can mimic the gene expression changes observed when treating HL-60 cells with PMA, OXS007417 and OXS007464. The top ranked compounds are shown in Table 5. The top ranking matches for our PMA-generated signatures were monopolised by signatures produced by PMA itself or ingenol 3,20-dibenzoate (also a PKC activating compound). Top ranking matches of OXS007417 and OXS007464 were, however, dominated by microtubule disruptors, confirming the BioMAP analysis. In addition, gene set enrichment analysis (GSEA) of OXS007417 and OXS007464 gene signatures confirmed the up-regulation of macrophage and neutrophil differentiation profiles at this time point (Fig. 5E).

**Table 5:** Top 14 similarity matches in L1000CDS^2^ data base of 6 h RNA-seq signatures.

|  | PMA |  |  | OXS007417 |  |  | OXS007464 |  |  |
| --- | --- | --- | --- | --- | --- | --- | --- | --- | --- |
| RANK | Perturbation | 1-cos($\alpha$) | Cell line | Perturbation | 1-cos($\alpha$) | Cell Line | Perturbation | 1-cos($\alpha$) | Cell Line |
| 1 | Ingenol 3, 20-dibenzoate | 0.6076 | PL12 | CYT997 | 0.5679 | PL21 | CYT997 | 0.5625 | PL2 |
| 2 | PMA | 0.632 | PL12 | PX12 | 0.5868 | PL21 | PX12 | 0.5773 | PL2 |
| 3 | Ingenol 3, 20-dibenzoate | 0.6407 | SKM1 | LY-2183240 | 0.5929 | THP1 | LY-2183240 | 0.5866 | THP1 |
| 4 | PMA | 0.6433 | NOMO1 | CYT997 | 0.5981 | THP1 | CYT997 | 0.5982 | THP1 |
| 5 | PX12 | 0.6581 | PL21 | ABT-751 | 0.6034 | PL21 | ABT-751 | 0.5993 | PL21 |
| 6 | Ingenol 3, 20-dibenzoate | 0.6613 | HA1E | LY-2183240 | 0.613 | PL21 | LY-2183240 | 0.603 | PL21 |
| 7 | PMA | 0.6658 | HA1E | PX12 | 0.6152 | THP1 | PX12 | 0.6161 | THP1 |
| 8 | PMA | 0.675 | SW60 | SB225002 | 0.6159 | THP1 | ABT-751 | 0.6199 | THP1 |
| 9 | BRD-k9114395 | 0.6828 | MCF7 | ABT-751 | 0.6192 | THP1 | SB-225002 | 0.6249 | THP1 |
| 10 | Ingenol 3, 20-dibenzoate | 0.6847 | SW60 | CYT997 | 0.6274 | SKM1 | CYT997 | 0.6298 | SKM1 |
| 11 | CYT997 | 0.6848 | PL21 | ABT-751 | 0.6333 | NOMO1 | PMA | 0.6346 | SW620 |
| 12 | PMA | 0.6865 | A375 | PMA | 0.6391 | SW620 | Ingenol 3, 20-dibenzoate | 0.6366 | SW620 |
| 13 | Ingenol 3, 20-dibenzoate | 0.6881 | MDST8 | SB225002 | 0.6395 | PL21 | ABT-751 | 0.6379 | NOMO1 |
| 14 | PX12 | 0.6885 | THP1 | Ingenol 3, 20-dibenzoate | 0.6436 | SW620 | BRD-K92317137 | 0.6394 | THP1 |

### Chemoproteomics with photoaffinity-labelled probes identifies tubulin as the drug target

Chemoproteomics using clickable photoaffinity labelled probes has proven to be a powerful and useful method for identification of small molecules’ direct target(s) within live cells ^41–45^. Particularly, UV-induced covalent bond formation between a photoaffinity-labelled probe and its target(s) has proven especially useful for target identification of small molecules which bind reversibly and with relatively low affinity to their targets through noncovalent interactions ^46^. The clickable alkyne tag enables the Cu(I)-catalyzed azide⍰alkyne cycloaddition (CuAAC) reaction with a fluorophore- or biotin-azide allowing visualization and isolation of bound target(s) by pull-down experiments respectively. Thus, on the basis of structure-activity relationship information of the parent compound OXS007464 ^38^, we devised an affinity-based protein profiling strategy to elucidate the protein targets of OXS007464 through the design and synthesis of clickable photoaffinity-labelled probe **4**, which retained the ability to upregulate CD11b with reasonable levels of potency (EC_50_ = 506 ± 44 nM, Fig. 6A).

**Figure 6:**
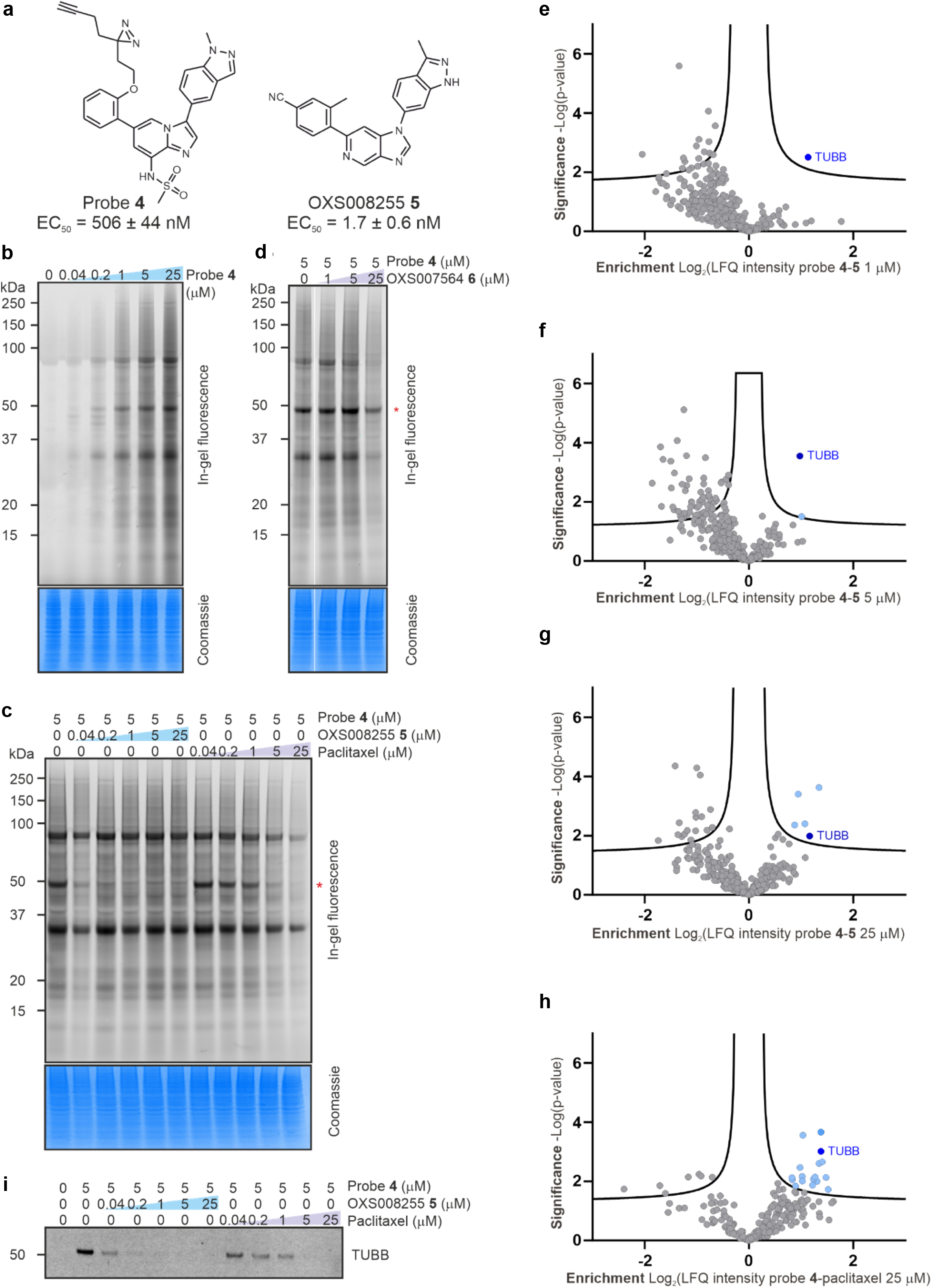
Target identification using chemical probe and chemoproteomics. A) Chemical structures of probe **4** and OXS008255 **5**. EC_50_ values for CD11b upregulation are represented as means ± SEM. In-gel fluorescence showing: B) dose-dependent labelling by probe **4**; C) competition of OXS008255 **5** and paclitaxel with probe **4**; D) competition of inactive analogue OXS007564 **6** with probe **4**. Coomassie stain shows equal protein loading on each gel. Uncropped gels are available in the supplementary information. Volcano plots showing significantly enriched proteins in the pull-down experiment by probe **4** compared to competition with: E) 1 μM of OXS008255 **5**; F) 5 μM of OXS008255 **5**; G) 25 μM of OXS008255 **5**; H) 25 μM of paclitaxel. Full list of proteins for each volcano plot is available in the supplementary information; I) Confirmation of tubulin beta chain enrichment by pull-down and immunoblotting.

In order to profile the ability of probe **4** to bind to proteins in cells, probe-treated HL-60 cells were exposed to irradiation (365 nm) and the resulting lysates were treated with TAMRA-azide under CuAAC conditions. Labelled proteins were separated by SDS-PAGE and visualized by in-gel fluorescence. Probe **4** demonstrated clear concentration-dependent labelling of proteins in HL-60 cells (Fig. 6B). Furthermore, it was confirmed that labelling is UV-dependent (Supplementary Fig. 4).

Next, we wanted to distinguish therapeutically relevant target(s) from non-specific binding by carrying out competition control experiments using an excess of more potent parent compounds to block the binding of probe **4**. HL-60 cells were pre-treated with increasing concentrations of parent compounds OXS007464 **3** and OXS008255 **5** (EC_50_ = 1.7 ± 0.6 nM, Fig. 6C-D), the latter compound was a result of further optimization of OXS007417. Furthermore, the competition experiment was performed with known tubulin binder paclitaxel ^47^. Only one band (^~^ 50 kDa, the known molecular weight of tubulin) was clearly competed away in all cases, (Fig. 6C, Supplementary Fig. 4), with OXS008255 **5** showing more potent competition than paclitaxel (Fig. 6C). Additionally, pre-treatment with OXS007564 **6** (Supplementary Fig. 5), a structurally similar but inactive analogue of OXS007464 **3**, had no influence on the binding of probe **4** up to 5 μM (Fig. 6D), confirming the importance of the 50 kDa band as a relevant target.

To confirm the identity of the 50 kDa band and to identify other potential targets that may be less abundant and not easy to identify in the gel-based assay, we next performed a proteome-wide pull-down experiment. HL-60 cells were treated with probe **4** alongside DMSO vehicle and competition controls consisting of OXS008255 **5** and paclitaxel. After photocrosslinking, the resulting lysates were subjected to CuAAC reaction with biotinylated AzRB capture reagent (Supplementary Fig. 6) ^48^. Bound proteins were isolated by using NeutrAvidin beads followed by on-bead digestion and analysis of resulting peptides by nanoLC-MS/MS (full proteomics data available in Supplemental data S1).

Data analysis revealed that tubulin beta chain (TUBB) was the protein most significantly enriched by probe **4** when compared to DMSO vehicle (Supplementary Fig. 7). Interestingly, competition experiments with OXS008255 **5** (Fig. 6E - 1 μM and 6F - 5 μM) highlighted the high selectivity of our compounds for the tubulin beta chain. Moreover, the competition experiments with 25 μM of OXS008255 **5** (Fig. 6G) and paclitaxel (Fig. 6H) indicated tubulin beta chain and several other proteins as significant, however, tubulin beta chain was the only consistent target across all four conditions. The identification of tubulin beta chain was further confirmed by immunoblotting after the pull-down experiment (Fig. 6I). The results clearly show concentration-dependent competition with OXS008255 **5** and paclitaxel confirming tubulin beta chain as a direct target of our compounds.

### OXS007417 disrupts tubulin polymerisation in a cell-free system and causes metaphase arrest *in vitro*

Next, OXS007417 and OXS007464 were tested for their ability to inhibit polymerisation of tubulin in a cell-free system. Both compounds were found to inhibit tubulin polymerisation with IC_50_ values of 1.1 μM OXS007464, and 1.7 μM for OXS007417 (Fig. 7A). The ability of OXS007417 to disrupt the cell cycle of HL-60 cells was subsequently analysed by DNA and P-H3 staining. Cell cycle analysis showed OXS007417 was able to cause G2-M mitotic arrest with cell cycle profiles comparable to those produced by positive control vinblastine, a known microtubule disruptor (Fig. 7B-C). Finally, mitotic spindle disruption was observed by immunohistochemistry (Fig. 7D). OXS007417 was found to disrupt spindle formation with spindle morphology comparable to that of vinblastine treated cells. Together, these result show that using an unbiased phenotypic screen, we were able to identify a novel series of compounds that induce differentiation of four AML cell lines by binding directly to tubulin and causing a G2-M cell cycle arrest.

**Figure 7:**
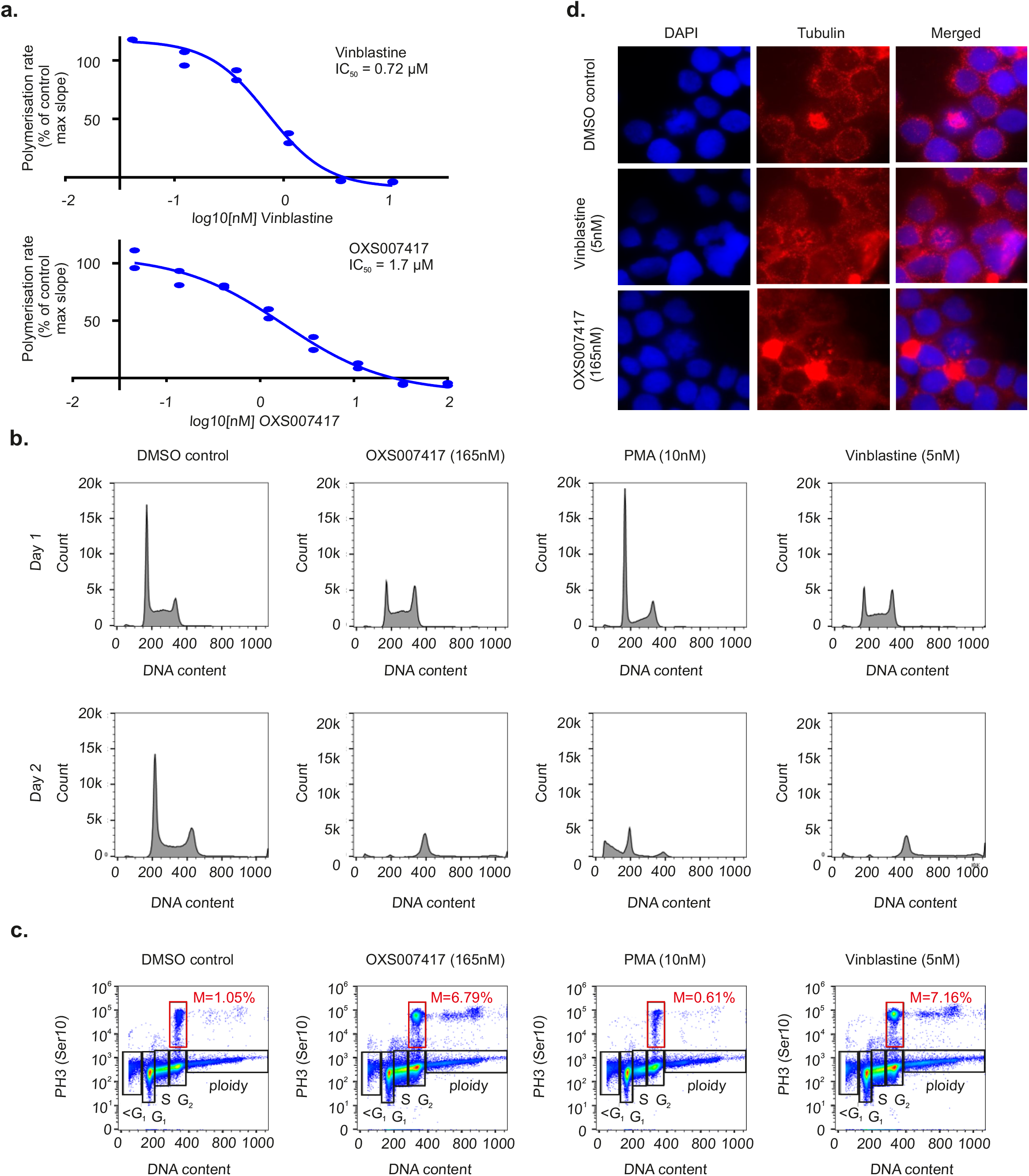
OXS007417 causes mitotic arrest via tubulin disruption in HL-60 cells. A) OXS007417 disrupts tubulin polymerisation in a cell free assay. B) Analysis of DNA content by flow cytometry demonstrates G2M arrest in OXS007417 treated HL-60 cells. C) PH3-staining confirms metaphase arrest upon treatment with OXS007417. D) Immunohistochemistry for tubulin (red) and counter staining for DAPI demonstrates spindle disruption by OXS007417.

## Discussion

Despite the long term need for new therapies in AML, it has only been the last few years that have produced several new therapies approved for use in the clinic ^3, 49^. However, despite these exciting recent advances, many of these new therapies are only able to target a subset of AML patients. In addition, even when new treatments are highly successful in targeting a specific patient subset, there are many individual patients who still do not respond fully to treatment. Thus, there is still a need for drugs that can target patients in a mutation and subtype independent manner, preferably with low toxicity either alone or in combination with other therapies. One promising new avenue of research to this end is the identification of compounds such as DHODH inhibitors that can induce differentiation of leukaemia cells ^31, 50^. Here, using an unbiased phenotypic screening approach, we have identified a novel class of tubulin binding molecules and have revealed a novel mechanism in which tubulin binding can cause a G2-M arrest and differentiation of AML cells.

In order to accomplish this, we used a flow cytometry based phenotypic screen with CD11b as a pan marker of myeloid differentiation to identify novel compounds that could differentiate several AML cell lines. Differentiation was confirmed by an arrest in cell proliferation, appearance of a differentiated cell morphology and by global gene expression analysis. Thereafter, through iterative rounds of optimisation of these hits, a number of lead molecules were developed from initial hits. Using these compounds *in vivo*, efficacy was demonstrated in two different xenograft models.

The advantage of phenotypic screening is that unlike target-based approaches, a well-designed phenotypic screen can directly select for compounds that have the desired cellular activity, in this case differentiation of AML cells. This provides the possibility of selecting for compounds that have polypharmacology as well as identifying compounds that impact desired pathways through novel mechanisms. It is generally not possible to easily accomplish either of these goals with single target-based approaches, as extensive knowledge of the target is a precondition of drug development. Because of this, it has been argued that successful target-based approaches have often been dependent on previous phenotypic screening for identification and development of first-in-class compounds, and that phenotypic screening is an important complementary approach to target based methods^51, 52^.

However, there are several challenges of phenotypic screens including the importance of including counterscreening approaches to filter out compounds which interact with undesirable, undruggable or toxicity inducing targets; as well as identifying the target of a novel compound and then understanding the underlying mechanism of how target engagement impacts the cell. Here, we used several complementary approaches to identify the direct target of our novel compounds and to ultimately decipher their mechanism of action. To accomplish this, representative lead compounds were subjected to BioMAP analysis, early time point RNA-seq analysis and chemoproteomics, with all three methods converging on tubulin disruption as the MoA. Furthermore, the compounds were found to upregulate differentiation signatures after just 6 h treatment.

Tubulin inhibitors are well known drugs in cancer therapy ^53^, but as far as we know they have not previously been identified as compounds that could cause differentiation of cells. More commonly, disruption of tubulin is known to lead to cell death, something we also observe in our data here. Exactly how a G2-M mediated arrest could lead to differentiation is not clear, but interestingly, DHODH inhibitors also seem to have this dual role in promoting a choice between differentiation or cell death ^31^.

In conclusion, we have shown that phenotypic screening can be employed to identify novel compounds that exhibit the desired phenotypic activity (differentiation induction) and that a variety of complementary methods can be used in the context of target deconvolution to decipher their mechanism of action. Further work needs to be done to identify other compounds capable of causing differentiation of AML cells, as this is likely to have a long-term impact on cancer therapy, especially in those areas currently lacking safe and effective treatments.

## Methods

### Cell Culture

AML cell lines were purchased from the American Type Culture Collection (ATCC; http://www.atcc.org) or from DSMZ (https://www.dsmz.de/). The cells were maintained in RPMI supplemented with 10% FBS and 1% *L*-Glutamine.

### Compound Treatment before Flow Cytometry

Compound stock solutions (10 mM) were prepared in DMSO and stored at −20 ºC. Serial dilutions were carried out in cell medium prior to use in each experiment and final concentration of DMSO was maintained at 0.1% except for final compound concentrations above 10 mM. Cells were seeded at in a 96-well plate at a density of 2×104 cells/well, in a 95 mL volume, then 5 mL of compound solutions (x20 of desired concentration) were added. Cells were incubated for 4 days

### Flow Cytometry

Cells were pelleted by centrifugation at 1000 rpm and suspended in 40 mL of blocking buffer (10% FBS in IMDM, no phenol red), then 10 mL of anti-human CD11b/Mac-1 (555388, BD Bioscience) solution (25% in blocking buffer) was added. Cells were stored in ice for 20 min. The cell suspension was centrifuged, washed three times with staining buffer (1% FBS in IMDM, no phenol red), and resuspended in 200 mL of staining buffer with 1 mg/mL DAPI (Sigma-Aldrich, D9542). Flow cytometry was performed on an Attune NxT flow cytometer (Thermo Fisher Scientific UK) with previous compensation. Data was analysed using Attune NxT software and Flow Jo (v9).

### Cell counts and viability assessment

Solution 13 containing acridin orange and DAPI was purchased from ChemoMetec (910-3013). After the appropriate cell treatment, one volume of solution 13 was added into 19 volumes of the pre-mixed cell suspension, and analysed using NucleoCounter^®^ NC-300TM (ChemoMetec).

### Cytospins and Modified Wright’s Staining

Cells were prepared in staining buffer (IMDM, no phenol red + 1% FBS) at a concentration of approximately 1 x 10^5^ cells/ml. Cytospins were made (1,000 rpm, 5 min), and the cells allowed to air-dry. Cells were stained with Modified Wright’s stain using a Hematek^®^. Stained cells were allowed to air-dry and coverslips were affixed with DPX mount prior to microscopy (Sigma-Aldrich, 06522).

### *In Vivo* Leukaemia Analysis: Subcutaneous Model

Female NOD SCID mice aged 5-7 weeks were used for the HL-60 subcutaneous models. Cells (5 x 10^6^ cells) were implanted subcutaneously in a Matrigel matrix (1:1) onto the flank of each mouse and allowed to grow to the pre-specified size of 150 mm^3^. Mice were grouped randomly into treatment groups based on their bodyweight to ensure even distribution. Mice were treated as indicated in Table 2. Tumours were measured 3 times per week using digital callipers. The length and width of the tumour were measured, and volume calculated using the following formula: volume = (length x width^2^)/2. The tumour control ratio (T/C) was calculated in the following way: ((Mean tumour volume on day 28 – mean starting volume)/ (Mean vehicle tumour volume on day 28 – mean vehicle starting volume))*100. The study was terminated the end of the 28-day treatment period.

### *In Vivo* Leukaemia Analysis: Subcutaneous Model Ethics

All protocols used in this study were approved by the Axis Bioservices Animal Welfare and Ethical Review Committee, and all procedures were carried out under the guidelines of the Animal (Scientific Procedures) Act 1986.

### *In Vivo* Leukaemia Analysis: Orthotopic Model

Female NCG recipient mice 6–8 weeks of age were used for the HL-60 orthotopic model. Cells (1 x 10^7^) were introduced intravenously by tail vein injection. Cells were given 7 days to engraft before commencement of treatment. Animals were then dosed with OXS007417 (PO, 10 mg/kg) for 3 weeks (Fig. 4B).

### Isolation of RNA

Total RNA for the RNA-seq was isolated using QIAGEN RNeasy-Plus Mini columns as per the manufacturer’s instructions. RNA purity was analysed using RNA Screen Tape with a TapeStation system (Agilent).

### RNA-Seq

Poly-A containing mRNA molecules were purified from total RNA using oligo-dT attached magnetic beads. Following purification, the mRNA was fragmented using divalent cations under elevated temperature. First strand cDNA was synthesised using random primers (NEB). Following second strand cDNA synthesis, cDNA libraries underwent end repair, a single adenylation of the 3’ ends and TRUE-seq adapter ligation. Libraries were enriched by PCR (15 cycles). Library quality was assessed by DNA Screen Tape with Tape Station system (Agilent), quantified by Qubit assay (Thermo Fisher Scientific) and pooled. Next-generation sequencing of pooled libraries was performed (Illumina NextSeq), resulting in approximately 10 million pairs of 75-bp reads per sample.

### Gene Expression Analysis

Following sequencing, QC analysis was conducted using the fastQC package (http://www.bioinformatics.babraham.ac.uk/projects/fastqc). Reads were mapped to the human genome assembly hg19 using STAR ^54^. The featureCounts function from the Subread package was used to quantify gene expression levels using standard parameters ^55^. This was used to identify differential gene expression globally, using the DESeq2 package ^56^.

### Analysis of Cell Cycle by Flow Cytometry

Cells were harvested at indicated time points, washed in PBS and suspended in hypotonic fluorochrome solution [50 μg/ml propidium iodide (PI), 0.1% (w/v) sodium citrate, 0.1% (v/v) Triton X-100] and stored for at least 1 h in the dark at 4 °C. Cells were washed in PBS and samples then incubated with anti-PH3 (1:40) in FACS staining buffer (IMDM, no phenol red + 10% FBS) for 20 min at 4 °C in the dark. Cells were washed in FACS buffer (IMDM, no phenol red + 1% FBS) and flow cytometry was performed using an Attune NxT. Results were analysed using FlowJo_V10 software.

### Immunohistochemistry

Cells were washed with PBS and resuspended in 100% FCS and cytospun onto coated slides using a Shandon Cytospin 4 (Thermo Scientific) at 30g for 5 min. Cytospins were fixed in methanol for 7 minutes at −20°C and dipped 10 times in ice-cold acetone. Slides were then washed three times in TBS 0.01% Tween 20 (TBST) for 5 minutes on a mechanical rocker. Cells were subsequently blocked for 15 minutes at room temperature in TBS, 0.05% Tween 20, 1% bovine serum albumin (BSA). Samples were covered with 50 μl of TBS, 0.025% Tween 20, 1% BSA containing anti-tubulin, overlain with a cover slip and incubated overnight at 4°C in a humid chamber. Cover slips were then removed and slides were washed three times for five minutes with TBST, covered with 50 μl containing the appropriate secondary antibody and incubated in the dark for 40 minutes. Slides were then washed three times for five minutes with TBST, and once for 2 minutes with PBS. Samples were then counter stained with 0.25 μg/ml DAPI for 1 min, mounted using ProLong Gold antifade reagent (Invitrogen) and imaged using a widefield fluorescence microscope (DeltaVision Elite, imsol).

### General protocol for treatment and lysis

HL-60 cells (2 x 10^6^ cells/mL in serum free RPMI media) were treated for 1 h with the probe **4** or DMSO vehicle at 37 °C. In the case of competition experiments, cells were pre-treated with competitor or DMSO vehicle for 30 min followed by 1 h treatment with probe **4**. Treated cells were pelleted and washed with PBS. The resulting pellets were resuspended in PBS and irradiated at 365 nm for 5 min (100 W lamp, VWR 36595-021) on ice. Cells were lysed in buffer containing 0.1% SDS, 1% Triton −X-100 and 1× EDTA-free protease inhibitor cocktail (Calbiochem set III, 539134) in PBS. Protein concentration of each lysate was determined using a BCA assay (Merck, 71285).

### In-gel fluorescence

40 μL of each lysate (concentrations adjusted to 1 μg/μL) was treated with 2.4 μL of premixed click chemistry mixture (final concentrations of 100 μM TAMRA-N_3_ (Sigma-Aldrich, 760757), 1 mM CuSO_4_, 1 mM TCEP and 100 μM TBTA) for 1 h. Proteins were precipitated using MeOH/CHCl_3_ and the resulting pellets washed twice with MeOH. The air-dried pellets were dissolved in 20 μL of 1× NuPAGE LDS buffer with 0.1% mercaptoethanol and heated at 95 °C for 5 min. The proteins were separated by NuPAGE 4-12% Bis-Tris gel in MES SDS running buffer. The gel was imaged using a Typhoon FLA 9500 scanner and then stained with Coomassie (InstantBlue™, Expedeon) and imaged using a BioRad ChemiDoc scanner.

### Proteomics

HL-60 cells were treated in triplicate and lysed as described above. 400 μL of each lysate (concentrations adjusted to 2.5 μg/μL) was treated with 24 μL of a click chemistry master mix (final concentrations of 100 μM AzRB, 1 mM CuSO_4_, 1 mM TCEP and 100 μM TBTA) for 1 h. The click reaction was quenched by adding 8 μL of 500 mM EDTA (10 mM final concentration). Proteins were precipitated using MeOH/CHCl_3_/H_2_O and the resulting pellets washed twice with MeOH. The air-dried pellets were dissolved in 80 μL of 1% SDS in 50 mM HEPES pH 8.0 by vortexing and sonicating and then diluted to 400 μL with 50 mM HEPES pH 8.0 (0.2% SDS final concentration).

Samples were incubated with 100 μL (1:10 ratio of bead suspension:protein) of NeutrAvidin agarose resin (Thermo Scientific 29201, pre-washed three times with 1 mL of 0.2% SDS in 50 mM HEPES pH 8.0) for 2 h at room temperature. The supernatants were removed and the beads washed three times with 1mL of 0.2% SDS in 50 mM HEPES pH 8.0 and then twice with 50 mM HEPES pH 8.0. The beads were then resuspended in 150 μL of 50 mM HEPES pH 8.0 and on-bead proteins were reduced with TCEP (5 mM final concentration) and alkylated with CAA (15 mM final concentration) for 10 min with gentle shaking. Proteins were digested overnight at 37 °C with 5 μL of trypsin (1 μg dissolved in 50 mM HEPES pH 8.0, Promega V5111). The trypsin digestion was quenched by adding 4 μL of 1× EDTA-free protease inhibitor cocktail (Roche 11873580001). The supernatants were collected and the beads washed (50μL) with 50 mM HEPES pH 8.0. The second wash was combined with the corresponding supernatant and vacuum-dried. The peptide solutions were desalted on stage-tips according to a published protocol ^57^. The peptides were eluted from the sorbent (Empore™ SDB-XC solid phase extraction discs, 3M, 2240) with 60% acetonitrile in water (60 μL), dried in a Savant SPD1010 SpeedVac^®^ Concentrator (Thermo Scientific) and stored at −80 °C until LC-MS/MS analysis. Peptides were reconstituted in 2% acetonitrile in water with 0.5% trifluoroacetic acid for LC-MS/MS analysis.

### NanoLC-MS/MS analysis

Peptides were separated on an EASY-Spray™ Acclaim PepMap C18 column (50 cm × 75 μm inner diameter, Thermo Fisher Scientific) using a binary solvent system of 2% acetonitrile with 0.1% formic acid (Solvent A) and 80% acetonitrile with 0.1% formic acid (Solvent B) in an Easy nLC-1000 system (Thermo Fisher Scientific). 2 μL of peptide solution was loaded using Solvent A onto an Acclaim PepMap100 C18 trap column (2 cm x 75 μm inner diameter), followed by a linear gradient separation of 0-100% Solvent B over 70 mins at a flow rate of 250 nL/min. Liquid chromatography was coupled to a QExactive mass spectrometer via an easy-spray source (Thermo Fisher Scientific). The QExactive was operated in data-dependent mode with survey scans acquired at a resolution of 70,000 at *m/z* 200 (transient time 256 ms). Up to 10 of the most abundant isotope patterns with charge +2 to +7 from the survey scan were selected with an isolation window of 2.0 m/z and fragmented by HCD with normalized collision energies of 25. The maximum ion injection times for the survey scan and the MS/MS scans (acquired with a resolution of 17 500 at m/z 200) were 20 and 120 ms, respectively. The ion target value for MS was set to 10^6^ and for MS/MS to 10^5^, and the intensity threshold was set to 8.3 × 10^2^.

### Proteomics database search and data analysis

Processing of LC-MS/MS data was performed in MaxQuant version 1.6.6.0 using the built-in Andromeda search engine. Peptides were identified from the MS/MS spectra searched against the human reference proteome (Uniprot, Taxon ID: 9606, accessed 4th September 2019). Cysteine carbamidomethylation was set as a fixed modification, and methionine oxidation and N-terminal acetylation were set as variable modifications. ‘Trypsin/P’ was chosen as digestion mode enzyme. Minimum peptide length was set to 7 residues and maximum 2 missed cleavages were allowed. ‘Unique and razor peptides’ were chosen for protein quantification. Quantification parameters were set to ‘standard’ and ‘LFQ’. Other parameters were used as pre-set in the software.

Data analysis was performed using Perseus (version 1.6.6.0). MaxQuant proteinGroups.txt output files were filtered against contaminants and reverse dataset. Base 2 logarithm was applied to all measurements and the median values within each sample were subtracted to normalise for sample variation associated with overall protein abundance. The replicates for each condition were grouped and the proteins with at least two valid values within a group were kept. A student’s t-test (FDR = 0.05; S0 = 0.1) was performed between the active probe sample and each DMSO control, and between active probe sample and probe/parent competition samples. For mathematical reasons ^58^, S_0_ was kept low. The results were plotted using GraphPad Prism.

### Data availability

Processed proteomics data are available in Supplementary tables 2-6. The raw mass spectrometry proteomics files and database search results have been deposited at the ProteomeXchange Consortium (http://proteomecentral.proteomexchange.org) via the PRIDE partner repository ^59^ with data set identifier PXD0022038.

### Western blotting

HL-60 cells were treated and lysed as described above. 100 μL of each lysate (concentrations adjusted to 2.5 μg/μL) was treated with 6 μL of premixed click chemistry mixture (final concentrations of 100 μM biotin-N_3_ (Sigma-Aldrich, 762024), 1 mM CuSO_4_, 1 mM TCEP and 100 μM TBTA) for 1 h. The click reactions were quenched by adding 2 μL of 500 mM EDTA (10 mM final concentration). Proteins were precipitated using MeOH/CHCl_3_/H_2_O and the resulting pellets were washed twice with MeOH. The air-dried pellets were dissolved in 80 μL of 1% SDS in PBS by vortexing and sonicating and then diluted to 400 μL with PBS.

Samples were incubated with 15 μL of MyOne™ Streptavidin T1 Dynabeads™ (Thermo Scientific, 29201, pre-washed three times with 0.2% SDS in PBS) for 1 h in the shaker. The supernatants were removed and the beads washed three times with 0.1% SDS, 1% TritonX-100 in PBS and then three times with 0.2% SDS in PBS. The beads were resuspended in 50 μL of 1× NuPAGE LDS buffer with 0.1% mercaptoethanol and heated at 95 °C for 5 min. The eluted proteins were separated by NuPAGE 4-12% Bis-Tris gel in MES SDS running buffer and transferred to a PVDF membrane (Bio-Rad, 162-0263). Tubulin beta chain protein was detected with an anti-TUBB antibody (1:500 in 5% fat-free milk solution in TBST, Invitrogen MA5-16308) followed by an anti-mouse secondary antibody (Alexa Fluor™ Plus 800, 1:10000, ThermoFisher A32730). The blots were imaged with a Licor Odyssey system.

### Microtubule polymerisation assay

The microtubule polymerisation assay was performed using porcine neuronal tubulin (Cytoskeleton, Inc, BK006P) as an adaptation of the original method of Shelanski et al. and Lee et al.^60, 61^ at Cytoskeleton, Inc.

## Acknowledgments

T.A.M. is supported by Medical Research Council (MRC, UK) Molecular Haematology Unit grant MC_UU_00016/6. T.R.J., A.V, L.J-C., T.J.C., and D.Z. and the experimental data were all supported with a grant from OxStem Oncology. We’d like to acknowledge Ashley David and Robert Hom from Cytoskeleton Inc (1830 S. Acoma St. Denver, CO 80223, USA) for suggestions and productive discussions.

## Author contributions

T.R.J., A.V, L.J-C., K.S.M., D.C., T.J.C., I.V.L.W., L.K., L.M., R.W., S.G.D., E.W.T., G.M.W., P.V., A.J.R. and T.A.M. conceived the experimental design; T.R.J., A.V, L.J-C., K.S.M., D.C., T.J.C., I.V.L.W., L.K., D.Z., and D.G. carried out experiments; T.R.J., A.V, L.J-C. and L.K. analysed and curated the data; T.R.J., A.V, L.J-C., R.W., S.G.D., E.W.T., G.M.W., P.V., A.J.R. and T.A.M. interpreted the data; R.M., A.D., A.O’M., R.W., G.C.T. and S.G.D. provided expertise; T.R.J., A.V, L.J-C., A.J.R. and T.A.M. wrote the manuscript; all authors contributed to reviewing and editing the manuscript; P.V., A.J.R. and T.A.M. provided supervision and funding.

## References

1. Cancer Research UK (http://www.cancerresearchuk.org/cancer-info/cancerstats/types/leukaemia-aml/.).

2. Siegel, R.L., Miller, K.D. & Jemal, A. Cancer statistics, 2019. CA Cancer J Clin 69, 7–34 (2019).

3. Daver, N. et al. New directions for emerging therapies in acute myeloid leukemia: the next chapter. Blood Cancer J 10, 107 (2020).

4. Yates, J.W., Wallace, H.J., Jr., Ellison, R.R. & Holland, J.F. Cytosine arabinoside (NSC-63878) and daunorubicin (NSC-83142) therapy in acute nonlymphocytic leukemia. Cancer Chemother Rep 57, 485–488 (1973).

5. Dombret, H. & Gardin, C. An update of current treatments for adult acute myeloid leukemia. Blood 127, 53–61 (2016).

6. Daver, N., Schlenk, R.F., Russell, N.H. & Levis, M.J. Targeting FLT3 mutations in AML: review of current knowledge and evidence. Leukemia 33, 299–312 (2019).

7. Liu, X. & Gong, Y. Isocitrate dehydrogenase inhibitors in acute myeloid leukemia. Biomark Res 7, 22 (2019).

8. Ivey, A. et al. Assessment of Minimal Residual Disease in Standard-Risk AML. N Engl J Med 374, 422–433 (2016).

9. Quek, L. et al. Clonal heterogeneity of acute myeloid leukemia treated with the IDH2 inhibitor enasidenib. Nat Med 24, 1167–1177 (2018).

10. Stein, E.M. et al. Molecular remission and response patterns in patients with mutant-IDH2 acute myeloid leukemia treated with enasidenib. Blood 133, 676–687 (2019).

11. Pollyea, D.A. et al. Enasidenib, an inhibitor of mutant IDH2 proteins, induces durable remissions in older patients with newly diagnosed acute myeloid leukemia. Leukemia 33, 2575–2584 (2019).

12. Bristol Myers Squibb (https://news.bms.com/press-release/corporatefinancial-news/bristol-myers-squibb-provides-update-phase-3-idhentify-trial-p 2020).

13. Cucchi, D.G.J. et al. Two decades of targeted therapies in acute myeloid leukemia. Leukemia (2021).

14. Jakobsen, N.A. & Vyas, P. From genomics to targeted treatment in haematological malignancies: a focus on acute myeloid leukaemia. Clin Med (Lond) 18, s47–s53 (2018).

15. DiNardo, C.D. et al. Azacitidine and Venetoclax in Previously Untreated Acute Myeloid Leukemia. N Engl J Med 383, 617–629 (2020).

16. DiNardo, C.D. et al. Safety and preliminary efficacy of venetoclax with decitabine or azacitidine in elderly patients with previously untreated acute myeloid leukaemia: a non-randomised, open-label, phase 1b study. Lancet Oncol 19, 216–228 (2018).

17. Killock, D. Venetoclax in AML: efficacy confirmed. Nat Rev Clin Oncol 17, 592 (2020).

18. Watts, J. & Nimer, S. Recent advances in the understanding and treatment of acute myeloid leukemia. F1000Res 7 (2018).

19. Wang, Z.Y. & Chen, Z. Acute promyelocytic leukemia: from highly fatal to highly curable. Blood 111, 2505–2515 (2008).

20. Tallman, M.S. & Altman, J.K. Curative strategies in acute promyelocytic leukemia. Hematology Am Soc Hematol Educ Program, 391–399 (2008).

21. Borrow, J., Goddard, A.D., Sheer, D. & Solomon, E. Molecular Analysis of Acute Promyelocytic Leukemia Breakpoint Cluster Region on Chromosome-17. Science 249, 1577–1580 (1990).

22. Dethe, H., Chomienne, C., Lanotte, M., Degos, L. & Dejean, A. The T(15-17) Translocation of Acute Promyelocytic Leukemia Fuses the Retinoic Acid Receptor-Alpha Gene to a Novel Transcribed Locus. Nature 347, 558–561 (1990).

23. Coombs, C.C., Tavakkoli, M. & Tallman, M.S. Acute promyelocytic leukemia: where did we start, where are we now, and the future. Blood Cancer Journal 5 (2015).

24. Miller, W.H., Jr., Schipper, H.M., Lee, J.S., Singer, J. & Waxman, S. Mechanisms of action of arsenic trioxide. Cancer Res 62, 3893–3903 (2002).

25. Dhillon, S. Ivosidenib: First Global Approval. Drugs 78, 1509–1516 (2018).

26. Kim, E.S. Enasidenib: First Global Approval. Drugs 77, 1705–1711 (2017).

27. Amatangelo, M.D. et al. Enasidenib induces acute myeloid leukemia cell differentiation to promote clinical response. Blood 130, 732–741 (2017).

28. DiNardo, C.D. et al. Durable Remissions with Ivosidenib in IDH1-Mutated Relapsed or Refractory AML. N Engl J Med 378, 2386–2398 (2018).

29. ClinicalTrials.gov (https://ClinicalTrials.gov/show/NCT03404726, 2021).

30. Sykes, D.B. et al. Inhibition of Dihydroorotate Dehydrogenase Overcomes Differentiation Blockade in Acute Myeloid Leukemia. Cell 167, 171–186 e115 (2016).

31. Christian, S. et al. The novel dihydroorotate dehydrogenase (DHODH) inhibitor BAY 2402234 triggers differentiation and is effective in the treatment of myeloid malignancies. Leukemia 33, 2403–2415 (2019).

32. Papaemmanuil, E. et al. Genomic Classification and Prognosis in Acute Myeloid Leukemia. New England Journal of Medicine 374, 2209–2221 (2016).

33. DiScipio, R.G., Daffern, P.J., Schraufstatter, I.U. & Sriramarao, P. Human polymorphonuclear leukocytes adhere to complement factor H through an interaction that involves alphaMbeta2 (CD11b/CD18). J Immunol 160, 4057–4066 (1998).

34. Losse, J., Zipfel, P.F. & Jozsi, M. Factor H and factor H-related protein 1 bind to human neutrophils via complement receptor 3, mediate attachment to Candida albicans, and enhance neutrophil antimicrobial activity. J Immunol 184, 912–921 (2010).

35. Lee, C.W., Sokoloski, J.A., Sartorelli, A.C. & Handschumacher, R.E. Induction of the differentiation of HL-60 cells by phorbol 12-myristate 13-acetate activates a Na(+)-dependent uridine-transport system. Involvement of protein kinase C. Biochem J 274 (Pt 1), 85–90 (1991).

36. Schenk, T. et al. Inhibition of the LSD1 (KDM1A) demethylase reactivates the all-trans-retinoic acid differentiation pathway in acute myeloid leukemia. Nat Med 18, 605–611 (2012).

37. Corces, M.R. et al. Lineage-specific and single-cell chromatin accessibility charts human hematopoiesis and leukemia evolution. Nat Genet 48, 1193–1203 (2016).

38. Josa-Culleré, L. et al. A phenotypic screen identifies a compound series that induces differentiation of acute myeloid leukemia cells in vitro and shows anti-tumour effects in vivo. bioRxiv (2020).

39. Altman, J.K. et al. Inhibition of Mnk kinase activity by cercosporamide and suppressive effects on acute myeloid leukemia precursors. Blood 121, 3675–3681 (2013).

40. Duan, Q. et al. L1000CDS(2): LINCS L1000 characteristic direction signatures search engine. NPJ Syst Biol Appl 2 (2016).

41. Eirich, J. et al. Pretubulysin derived probes as novel tools for monitoring the microtubule network via activity-based protein profiling and fluorescence microscopy. Mol Biosyst 8, 2067–2075 (2012).

42. Li, W. et al. Chemoproteomics Reveals the Antiproliferative Potential of Parkinson’s Disease Kinase Inhibitor LRRK2-IN-1 by Targeting PCNA Protein. Mol Pharm 15, 3252–3259 (2018).

43. Parker, C.G. et al. Ligand and Target Discovery by Fragment-Based Screening in Human Cells. Cell 168, 527–541 e529 (2017).

44. Shi, H., Zhang, C.J., Chen, G.Y. & Yao, S.Q. Cell-based proteome profiling of potential dasatinib targets by use of affinity-based probes. J Am Chem Soc 134, 3001–3014 (2012).

45. van Delft, M.F. et al. A small molecule interacts with VDAC2 to block mouse BAK-driven apoptosis. Nat Chem Biol 15, 1057–1066 (2019).

46. Wilkinson, I.V.L., Terstappen, G.C. & Russell, A.J. Combining experimental strategies for successful target deconvolution. Drug Discov Today (2020).

47. Nogales, E., Wolf, S.G., Khan, I.A., Luduena, R.F. & Downing, K.H. Structure of tubulin at 6.5 A and location of the taxol-binding site. Nature 375, 424–427 (1995).

48. Broncel, M. et al. Multifunctional reagents for quantitative proteome-wide analysis of protein modification in human cells and dynamic profiling of protein lipidation during vertebrate development. Angew Chem Int Ed Engl 54, 5948–5951 (2015).

49. Patel, S.A. & Gerber, J.M. A User’s Guide to Novel Therapies for Acute Myeloid Leukemia. Clin Lymphoma Myeloma Leuk 20, 277–288 (2020).

50. Sykes, D.B. The emergence of dihydroorotate dehydrogenase (DHODH) as a therapeutic target in acute myeloid leukemia. Expert Opin Ther Targets 22, 893–898 (2018).

51. Swinney, D.C. The contribution of mechanistic understanding to phenotypic screening for first-in-class medicines. J Biomol Screen 18, 1186–1192 (2013).

52. Moffat, J.G., Vincent, F., Lee, J.A., Eder, J. & Prunotto, M. Opportunities and challenges in phenotypic drug discovery: an industry perspective. Nat Rev Drug Discov 16, 531–543 (2017).

53. Arnst, K.E. et al. Current advances of tubulin inhibitors as dual acting small molecules for cancer therapy. Med Res Rev 39, 1398–1426 (2019).

54. Dobin, A. & Gingeras, T.R. Mapping RNA-seq Reads with STAR. Curr Protoc Bioinformatics 51, 11 14 11–11 14 19 (2015).

55. Liao, Y., Smyth, G.K. & Shi, W. The R package Rsubread is easier, faster, cheaper and better for alignment and quantification of RNA sequencing reads. Nucleic Acids Res 47, e47 (2019).

56. Love, M.I., Huber, W. & Anders, S. Moderated estimation of fold change and dispersion for RNA-seq data with DESeq2. Genome Biol 15, 550 (2014).

57. Rappsilber, J., Ishihama, Y. & Mann, M. Stop and go extraction tips for matrix-assisted laser desorption/ionization, nanoelectrospray, and LC/MS sample pretreatment in proteomics. Anal Chem 75, 663–670 (2003).

58. Giai Gianetto, Q., Coute, Y., Bruley, C. & Burger, T. Uses and misuses of the fudge factor in quantitative discovery proteomics. Proteomics 16, 1955–1960 (2016).

59. Perez-Riverol, Y. et al. The PRIDE database and related tools and resources in 2019: improving support for quantification data. Nucleic Acids Res 47, D442–D450 (2019).

60. Shelanski, M.L., Gaskin, F. & Cantor, C.R. Microtubule assembly in the absence of added nucleotides. Proc Natl Acad Sci U S A 70, 765–768 (1973).

61. Lee, J.C. & Timasheff, S.N. In vitro reconstitution of calf brain microtubules: effects of solution variables. Biochemistry 16, 1754–1764 (1977).

