## Supplemental Figures for "A novel tubulin binding molecule drives differentiation of acute myeloid leukaemia cells"

Supplementary Figure 1: High-Throughput screening in multiple AML cell lines. A) Upon treatment with positive control, PMA, KG1 cells differentiate and up-regulate sell surface marker CD11b as detected by flow cytometry (left). Scatter plot distribution showing the results of HL-60 high-throughput screening of 1000 compound library (right). B) Upon treatment with positive control, GS87 (30µM), OCI-AML3 cells differentiate and up-regulate sell surface marker CD11b as detected by flow cytometry (left). Scatter plot distribution showing the results of HL-60 high-throughput screening of 1000 compound library (right). C) Upon treatment with positive control, TCP, THP-1 cells differentiate and up-regulate sell surface marker CD11b as detected by flow cytometry (left). Scatter plot distribution showing the results of HL-60 high-throughput screening of 1000 compound library (right).
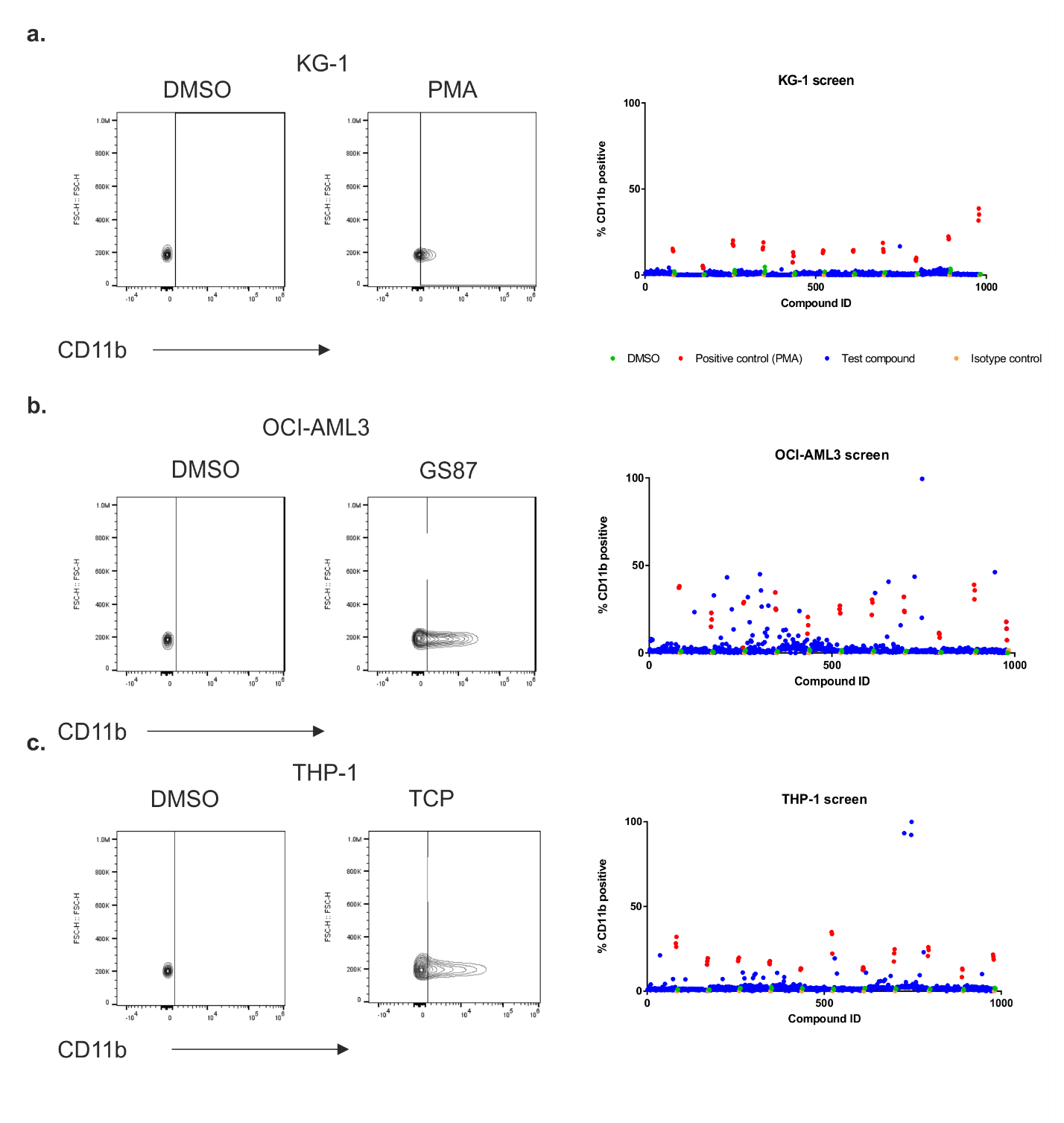

Supplementary Figure 2: Macrophage-like morphology observed in AML cell lines treated with OXS000275. Cytospin preparations of OXS000275-treated cells lines stained with Wright-Giemsa showed signs of myeloid maturation.

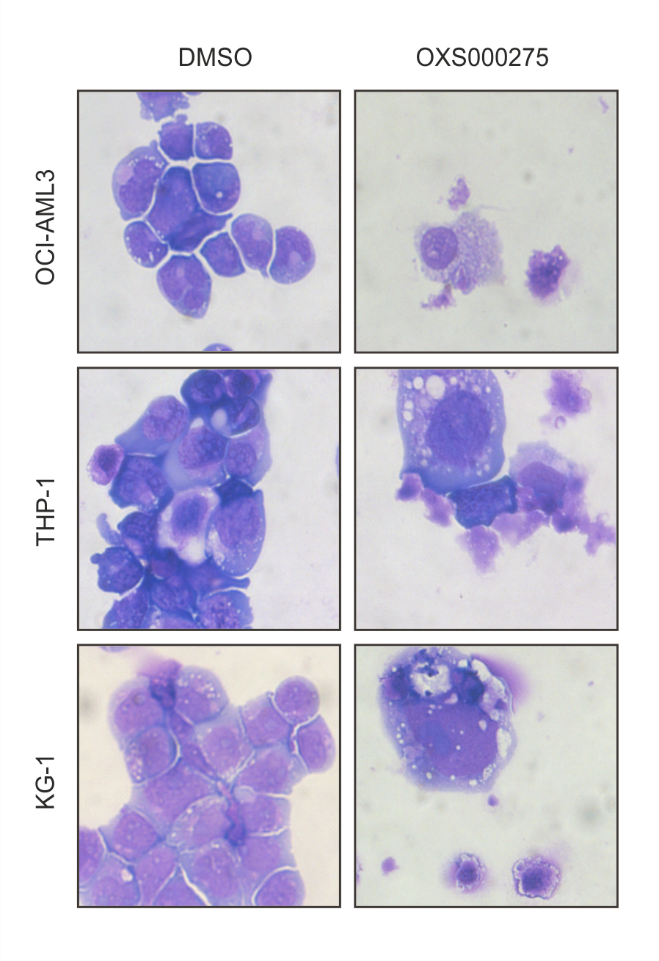

Supplementary Figure 3: Normalized body weight of mice in the subcutaneous xenograft study assessed over time with indicated treatments. Red arrow indicated start of treatment.

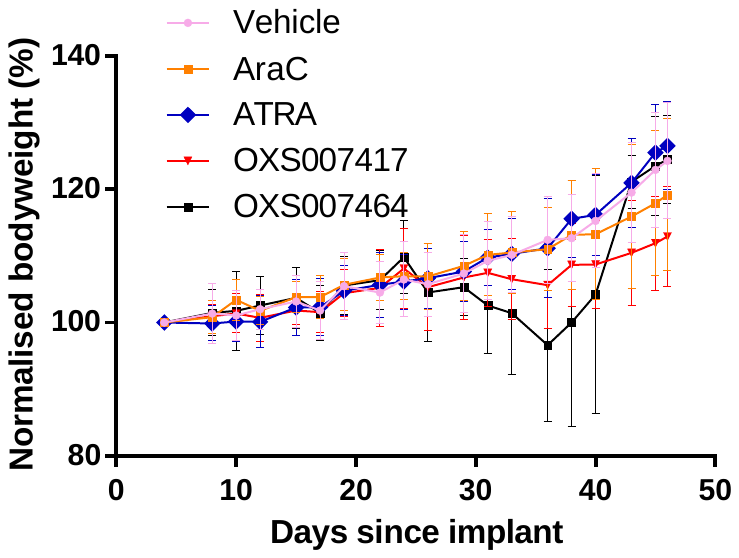

Supplementary Figure 4. In-gel fluorescence showing non-irradiated control and competition of OXS007464 **3** and OXS007564 **6** with probe **4.**

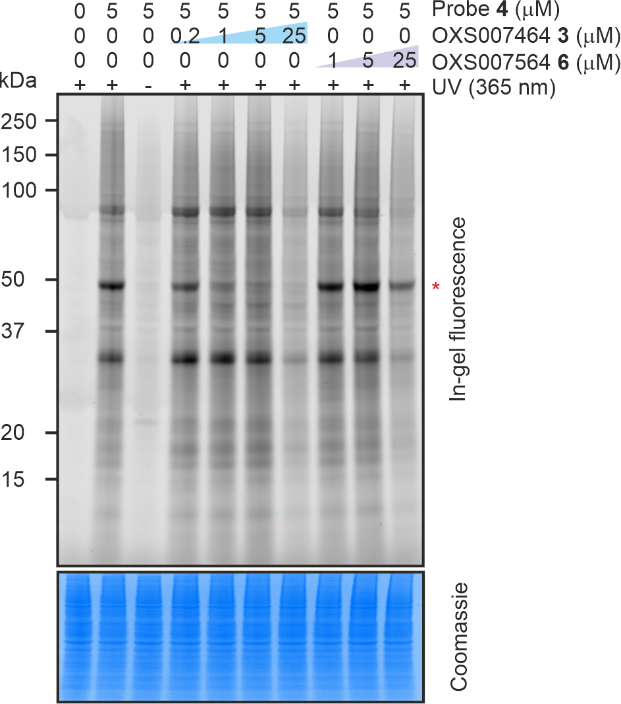

Supplementary Figure 5. Chemical structure of OXS007564 **6**. EC_50_ > 10 µM.

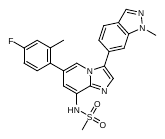

Supplementary Figure 6. Chemical structure of AzRB capture reagent. (PMID: 25807930)

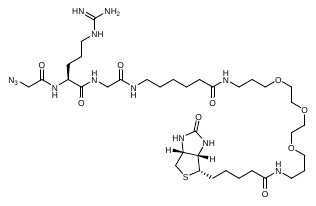

Supplementary Figure 7. Volcano plot showing significantly enriched proteins in the pull-down experiment by probe **4** compared to DMSO vehicle.

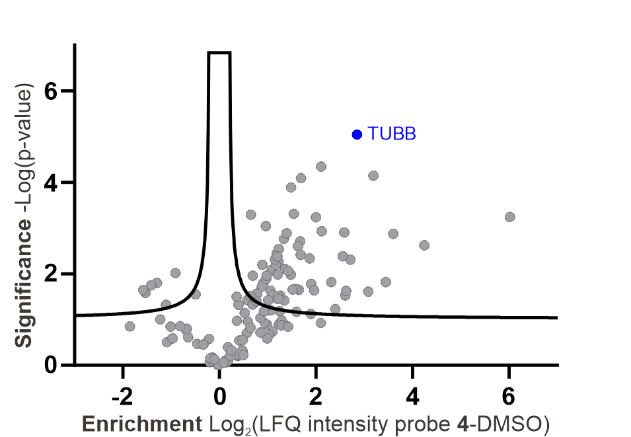

Supplementary Figure 8. Uncropped gels.

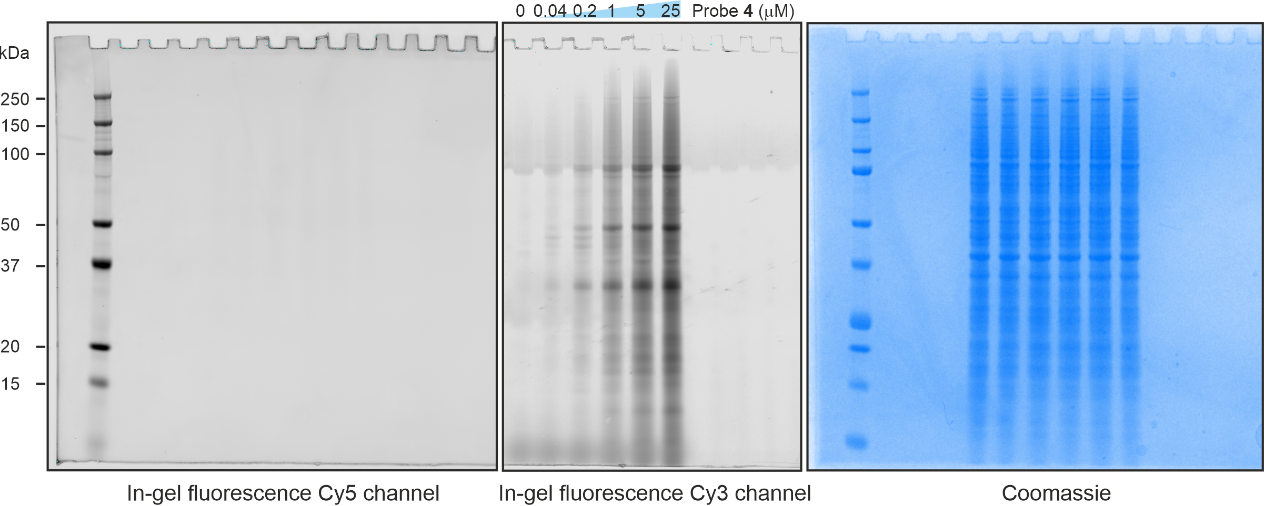

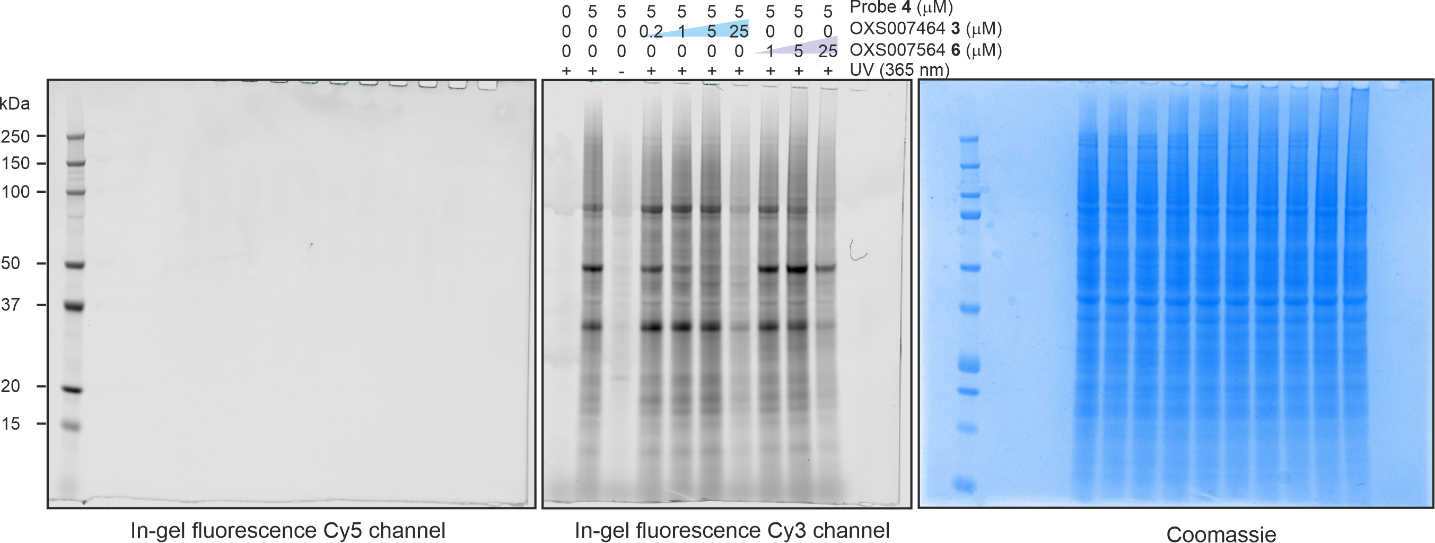

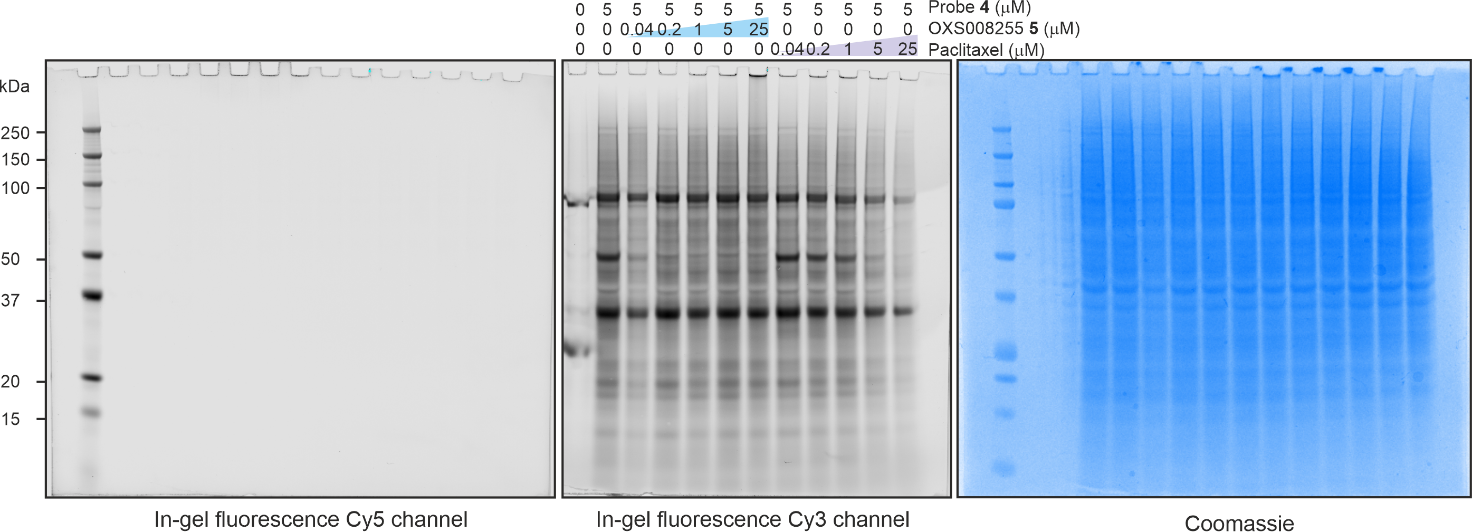

Supplementary Figure 8. Uncropped western blot.

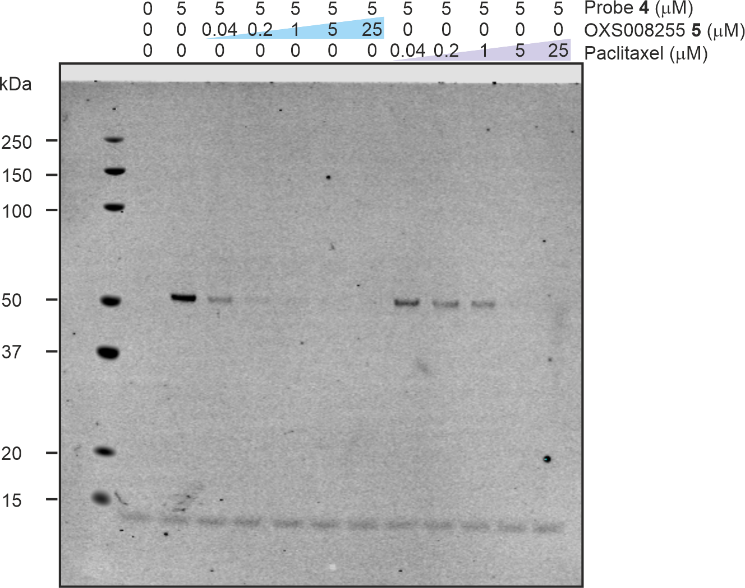

Supplementary Table 1. Summary of EnrichR analysis of RNA-seq signatures for a range in chemically distinct screening hits

| Compound | FACS  %CD11b | log2foldchange  CD11b | Total up | Total  down | Macrophage  No. up | Macrophage  % of up | Fisher exact  p-value | Neutrophil  No. up | Neutrophil  % of up | Fisher  exact  p-value |
| --- | --- | --- | --- | --- | --- | --- | --- | --- | --- | --- |
| OXS000651 | 15 | 1.5 | 1241 | 973 | 294 | 23.69057 | 1.92E-42 | 299 | 24.09347 | 1.01E-44 |
| OXS006988 | 16 | 1.8 | 2704 | 2382 | 301 | 11.13166 | 1.02E-08 | 282 | 10.42899 | 0.001 |
| OXS006996 | 42 | 2.8 | 2544 | 2312 | 420 | 16.50943 | 3.45E-22 | 517 | 20.32233 | 3.91E-54 |
| OXS003976 | 72.5 | 3.4 | 4183 | 3717 | 1429 | 34.16208 | NA | 724 | 17.30815 | 2.7E-49 |
| OXS006976 | 20 | 2.3 | 2827 | 2434 | 343 | 12.133 | 0.005964 | 333 | 11.77927 | 0.033432 |
| OXS004030 | 58 | 4.4 | 3384 | 3144 | 700 | 20.68558 | 7.8E-251 | 504 | 14.89362 | 3.12E-15 |
| OXS000493 | 25 | 2 | 2869 | 2714 | 260 | 9.062391 | 0.86 | 248 | 8.644127 | 0.8 |
| OXS000366 | 20 | 1.3 | 1395 | 1380 | 161 | 11.54122 | 0.243117 | 150 | 10.75269 | 2E-12 |
| PMA | 80 | 5.4 | 4255 | 3738 | 748 | 17.57932 | 2.64E-54 | 504 | 11.84489 | 0.004274 |
| ATRA | 24 | 3.4 | 3133 | 2753 | 808 | 25.78998 | 9.9E-156 | 571 | 18.22534 | 4.2E-44 |
| OXS006974 (inactive control) |  | 0.355725 | 45 | 0 | 3 | 6.666667 | 0.624736 | 0 | 0 | NA |

Supplementary Table 2. Significantly enriched proteins by probe **4** compared to DMSO. Full dataset is available in supplementary Excel file.

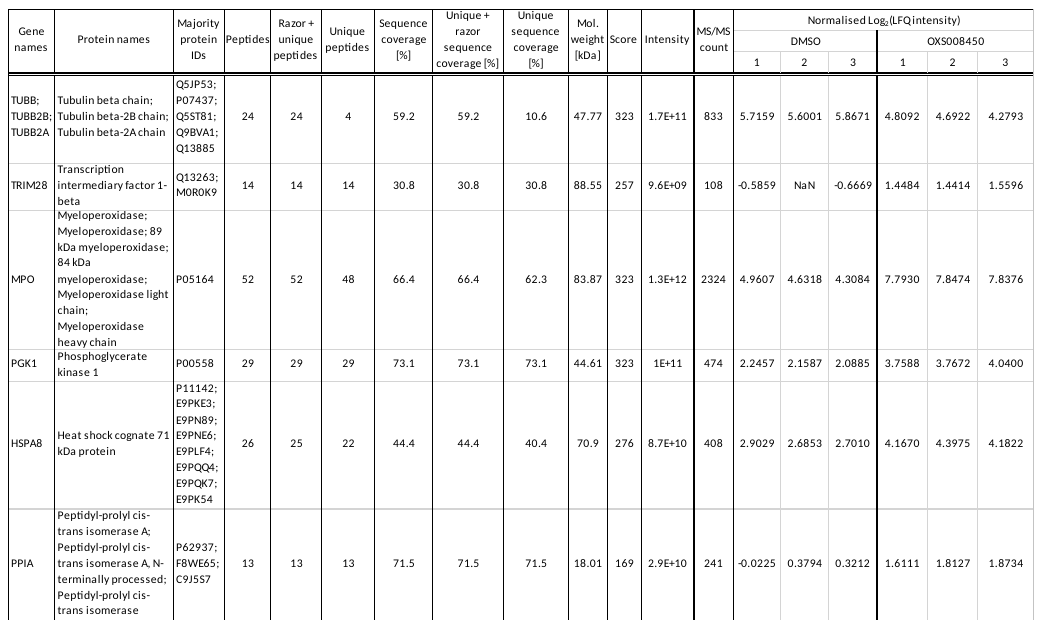

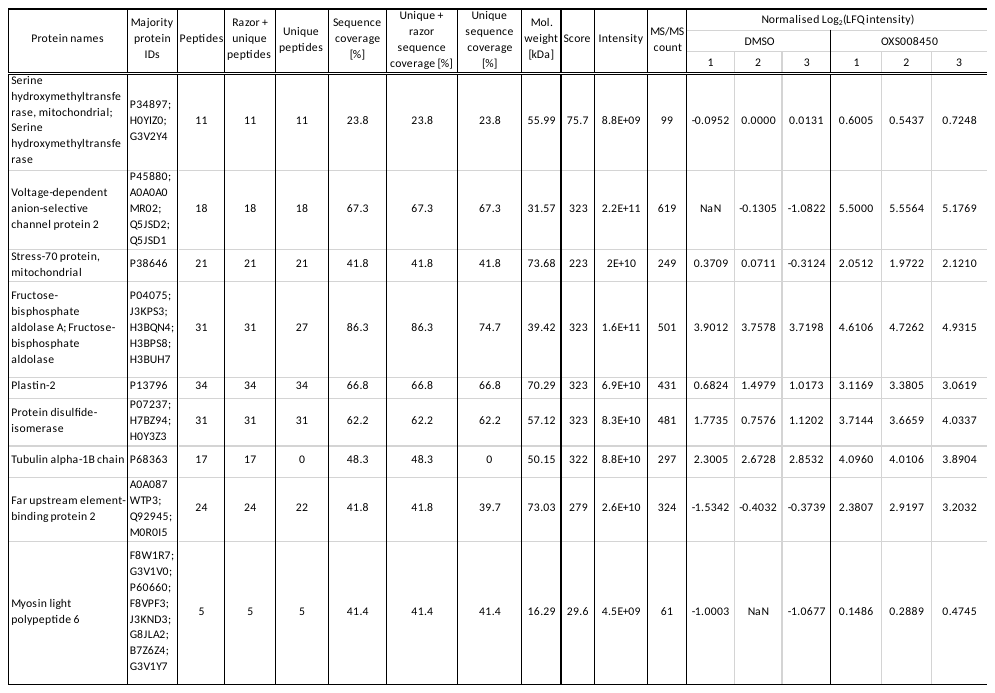

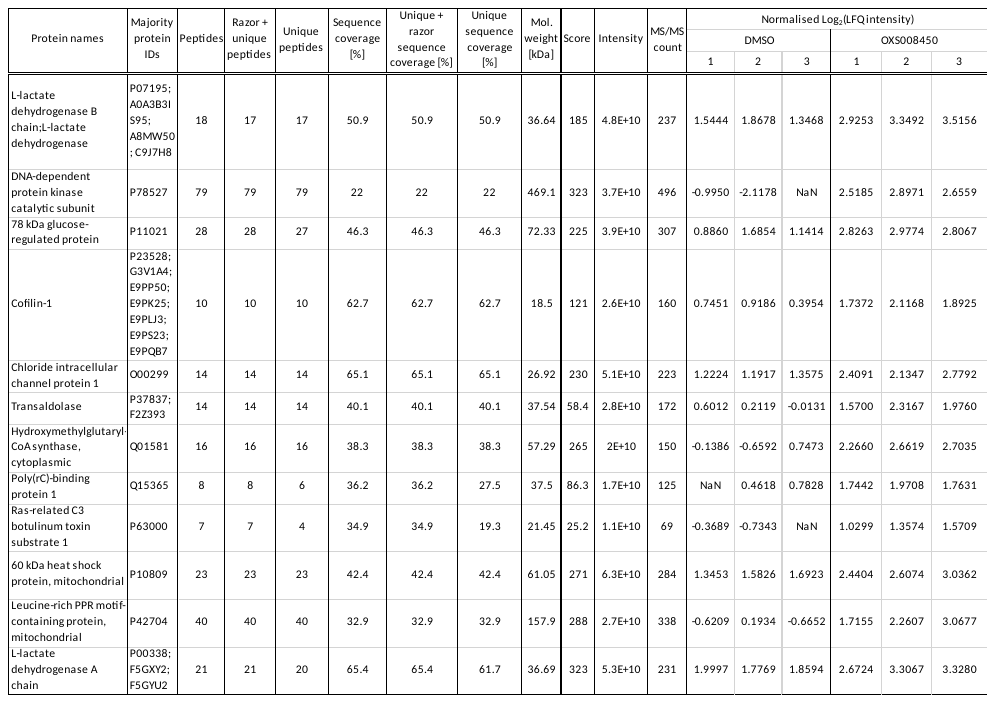

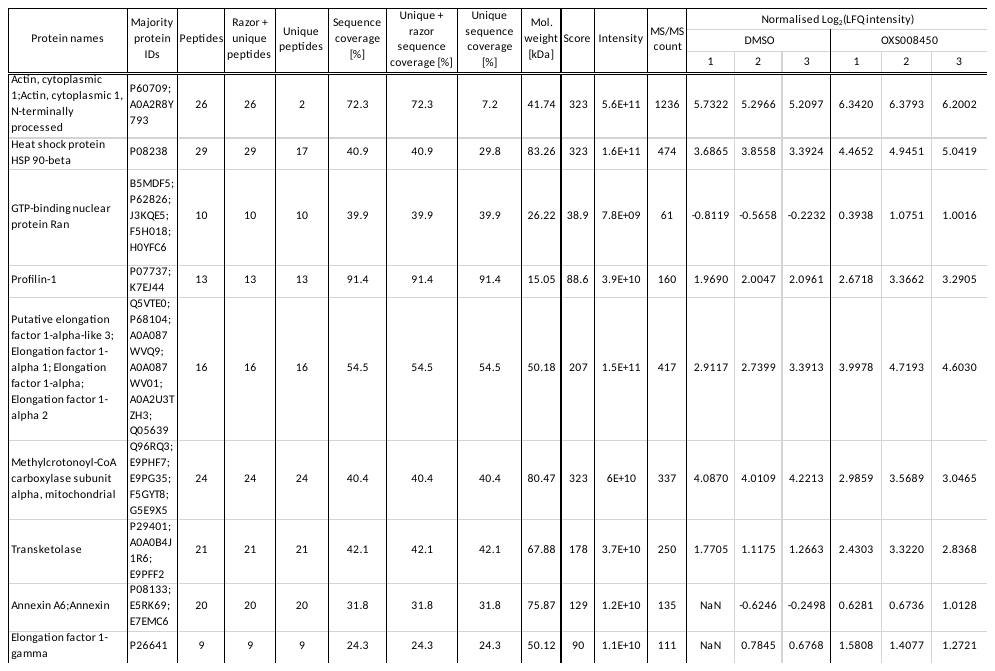

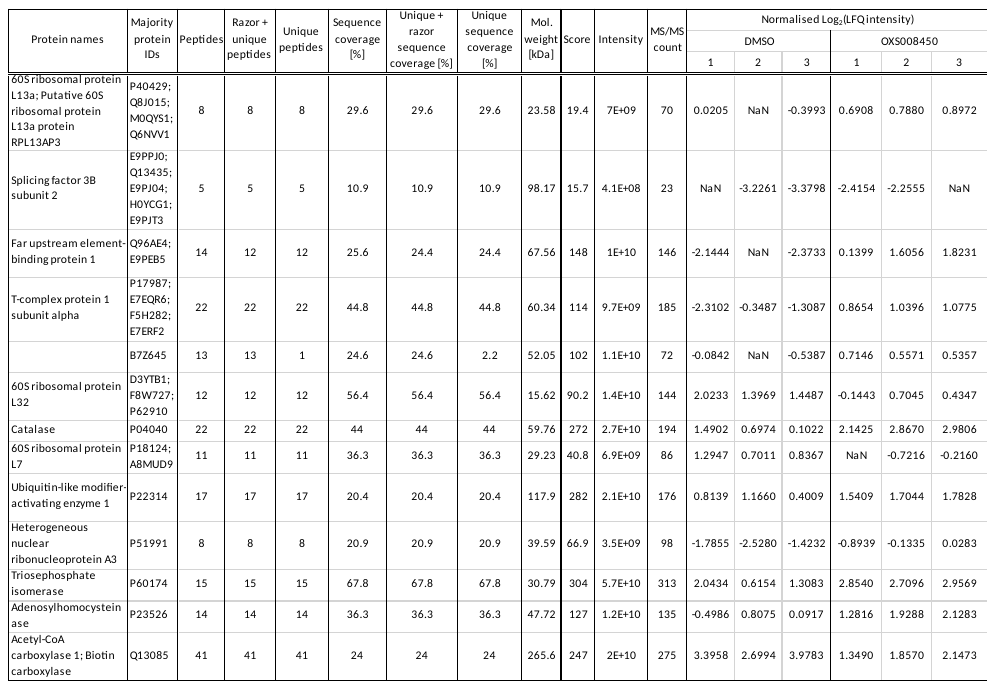

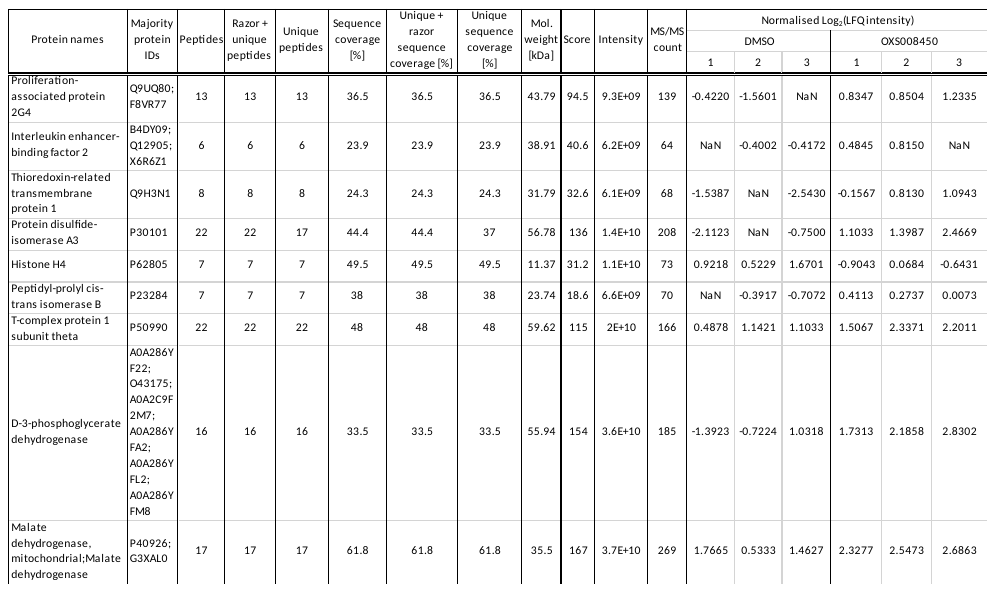

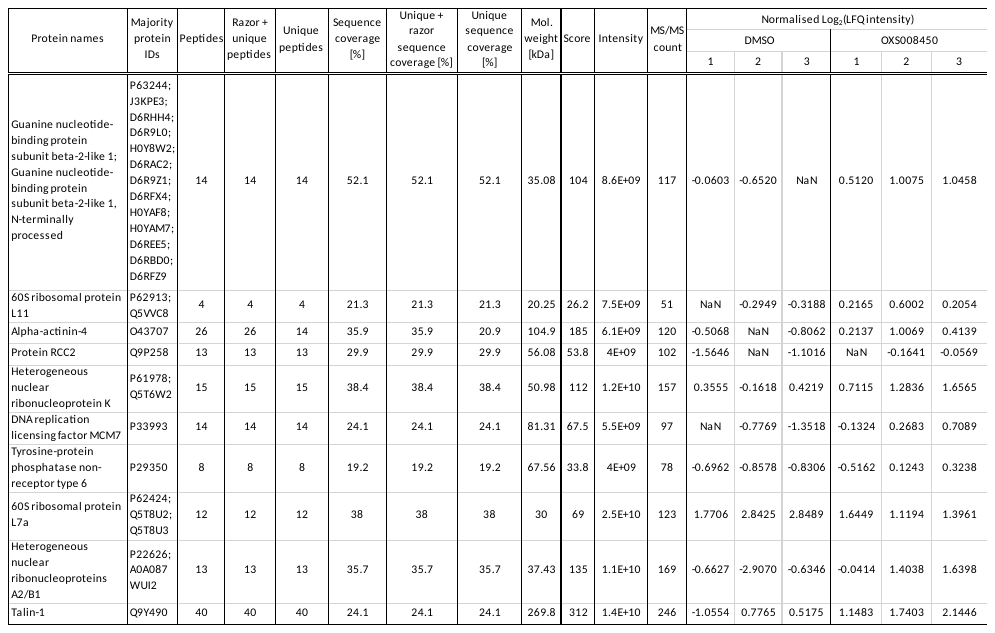

Supplementary table 3. Significantly enriched proteins by probe **4** compared to competition with OXS008255 **5** 1 µM. Full dataset is available in supplementary excel file.

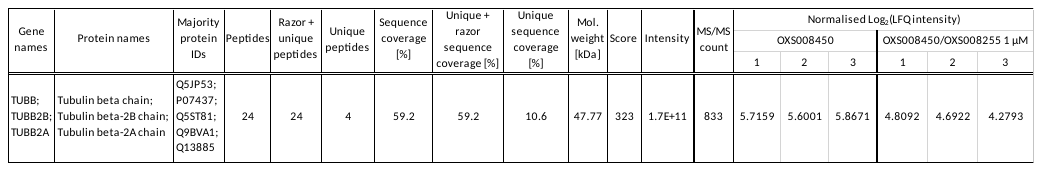

Supplementary table 4. Significantly enriched proteins by probe **4** compared to competition with OXS008255 **5** 5 µM. Full dataset is available in supplementary excel file.

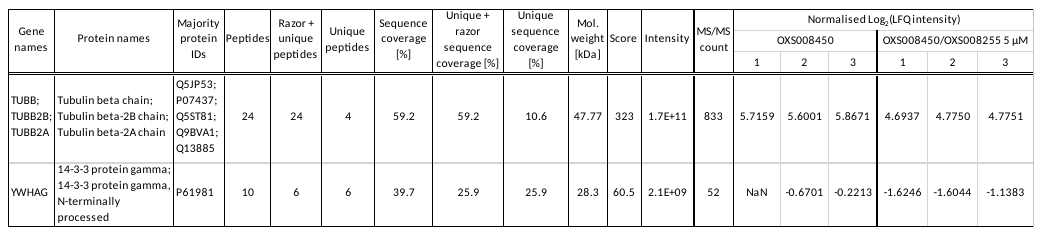

Supplementary Table 5. Significantly enriched proteins by probe **4** compared to competition with OXS008255 **5 (**25 µM). Full data is available in supplementary Excel file.

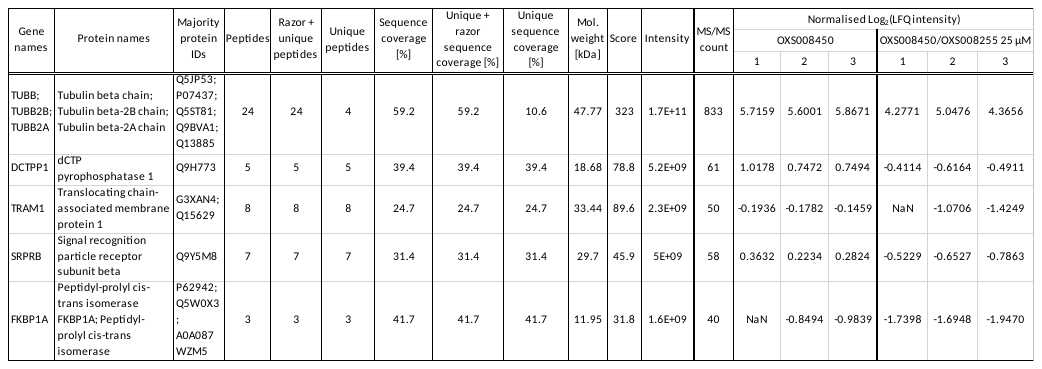

Supplementary Table 6. Significantly enriched proteins by probe **4** compared to competition with paclitaxel **5 (**25 µM). Full dataset is available in supplementary Excel file.

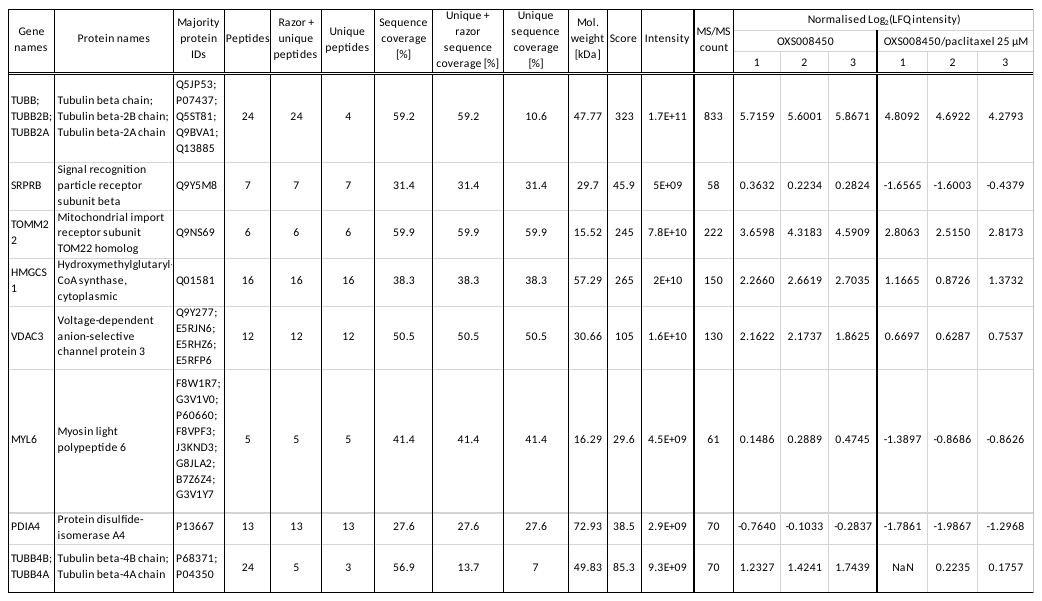

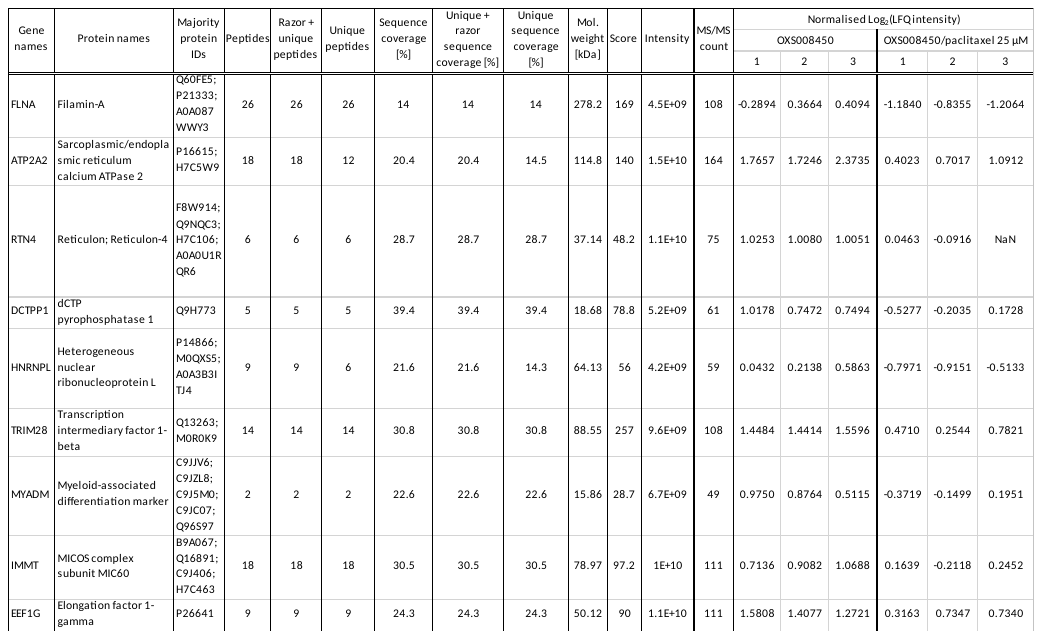

Supplementary Table 7: Structurally distinct tubulin disruptors up regulate CD11b as measured by FACS.

| **Compound** | **CAS** | **Compound** | **%CD11b EC_50_ (nM)** |
| --- | --- | --- | --- |
| nocodazol | 31430-18-9 | Nacodazol | 97.1 ± 1.4 |
| ABT-751 | 141430-65-1 | ABT-751 | 855.0 ± 38.2 |
| paclitaxel | 33069-62-4 | Paclitaxel | 4.3 ± 7.2 |
| CYT997 | 917111-44-5 | CYT997 | 15.6 ± 0.42 |
| colchicine | 64-86-8 | Colchicine | 11.8 ± 1.2 |
| vinblastine | 865-21-4 | Vinblastine | 1.89 ± 0.4 |
