## Supplementary Note for "A novel tubulin binding molecule drives differentiation of acute myeloid leukaemia cells"

**General chemistry experimental**

All reactions involving moisture sensitive reagents were carried out under a nitrogen or argon atmosphere. Solvents were dried following the procedure outlined by Grubbs et al.^54^ Water was purified by an Elix® UV-10 system. All other solvents and reagents were used as supplied (analytical or HPLC grade) without prior purification. Organic layers were dried over anhydrous Na_2_SO_4_ or MgSO_4_. Brine refers to a saturated aqueous solution of NaCl. *In vacuo* refers to the use of a rotary evaporator attached to a diaphragm pump. Analytical thin layer chromatography (TLC) was performed on Merck aluminium plates coated with 60 F_254_ silica. Plates were visualised using UV irradiation (λ 254 nm) and staining with a KMnO_4_ solution. Flash column chromatography was performed on Kieselgel 60 silica gel (230-400 mesh particle size) in a glass column, or using a Biotage Isolera One 3.0 or SP4 automated purification system with the default settings for a KP-Sil cartridge, monitoring at 254 and 280 nm. NMR spectra were recorded on Bruker Advance spectrometers in the deuterated solvent stated. The field was locked by external referencing to the relevant deuteron resonance. Chemical shifts (δ) are reported in parts per million (ppm) and coupling constants (*J*), determined by analysis using MestreNova software, are quoted in Hz. Data are reported as follows: chemical shift, multiplicity (s = singlet, bs = broad singlet, d = doublet, t = triplet, q = quartet and m = multiplet), coupling constant and integration. Low-resolution mass spectra (*m/z*) were recorded on an Agilent 1260 Infinity II with Diode Array and Single Quadrupole Detectors in solutions of MeOH. A selected peak is reported in Daltons and its intensity given as percentage of the base peak. High resolution mass spectra (HRMS) were run on a Bruker microTOF (ESI and APCI) or on a Waters GCT (EI), by the mass spectrometry department of the Chemistry Research Laboratory, University of Oxford, UK. Temperatures below 25 °C were obtained using the following cooling baths: 0 °C ice/water, -15 °C dry ice/ethylene glycol and -78 °C dry ice/acetone.

**Synthesis of OXS000275 (1)**

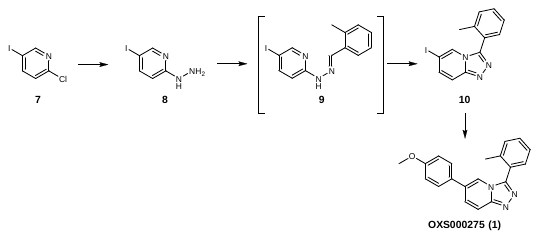

**(5-Iodo-2-pyridyl)hydrazine (8)**

To a solution of 2-chloro-5-iodopyridine **7** (2.48 g, 10.4 mmol) in pyridine (30 mL) was added hydrazine hydrate (5.04 mL, 104 mmol) dropwise. The mixture was refluxed overnight. The solution was concentrated to give a solid, which was triturated in pentane (5 mL) to give hydrazine **8** (2.4 g, 10 mmol, quant.) as a grey solid. ^1^H NMR (400 MHz, CD_3_OD) δ 8.02 (s, 1H), 7.56 (dd, *J* = 8.9, 2.2 Hz, 1H), 6.52 (d, *J* = 8.9 Hz, 1H); *m/z* (ESI^+^) 235.9 ([M+H]^+^, 100%); HRMS (ESI^+^) C_5_H_7_N_3_I^+^ ([M+H]^+^) requires 235.9679; found 235.9679. The data was in agreement with the reported values.^1^

**6-Iodo-3-(*o*-tolyl)-[1,2,4]triazolo[4,3-*a*]pyridine (10)**

To a suspension of hydrazine **8** (875 mg, 3.72 mmol) in EtOH (12 mL) was added 2-methylbenzaldehyde (447 mg, 3.72 mmol). The mixture was heated to reflux for 2 h, then cooled to room temperature. The resulting solid was filtered and washed with EtOH to give the hydrazone **9**. To a suspension of the crude hydrazone **9** in anhydrous CH_2_Cl_2_/MeOH (10:2, 19 mL) was added (diacetoxyiodo)benzene (1.68 g, 5.21 mmol) at 0 °C. The mixture was stirred at room temperature overnight, then diluted with water, extracted three times with CH_2_Cl_2_ and dried over anhydrous Na_2_SO_4_. The crude product was purified using flash column chromatography to give the triazolopyridine **10** (748 mg, 2.23 mmol, 60%) as an off-white solid. ^1^H NMR (CDCl_3_, 500 MHz) δ 8.03 (t, *J* = 1.3 Hz, 1H), 7.61 (dd, *J* = 9.6, 1.0 Hz, 1H), 7.47 (td, *J* = 7.5, 1.5 Hz, 1H), 7.44–7.39 (m, 3H), 7.37 (t, *J* = 7.4 Hz, 1H), 2.25 (s, 3H); ^13^C NMR (CDCl_3_, 125 MHz) δ 148.5, 145.8, 138.7, 135.1, 131.3, 130.8, 130.2, 127.6, 126.4, 125.0, 117.5, 77.7, 19.9; *m/z* (ESI^+^) 336.0 ([M+H]^+^, 100%); HRMS (ESI^+^) C_13_H_11_N_3_I^+^ ([M+H]^+^) requires 335.9992; found 335.9988.

**6-(4-Methoxyphenyl)-3-(*o*-tolyl)-[1,2,4]triazolo[4,3-*a*]pyridine (OXS000275, 1)**

To a solution of iodide **10** (126 mg, 0.378 mmol) in degassed 1,4-dioxane/water (5:2, 3.8 mL) were added (4-methoxyphenyl)boronic acid (69 mg, 0.45 mmol), potassium carbonate (78 mg, 0.57 mmol) and Pd(dppf)Cl_2_ (14 mg, 0.19 mmol), and the mixture was heated to 100 °C. After 16 h, the mixture was diluted with EtOAc, filtered through Celite® and concentrated. The crude product was purified using flash column chromatography to give **OXS000275 (1)** (112 mg, 0.355 mmol, 94%) as a brown solid. ^1^H NMR (400 MHz, CDCl_3_) δ 7.85 (dd, *J* = 9.5, 0.9 Hz, 1H), 7.82 (s, 1H), 7.52 (dd, *J* = 9.5, 1.6 Hz, 1H), 7.50–7.33 (m, 6H), 6.96 (d, *J* = 8.8 Hz, 2H), 3.82 (s, 3H), 2.29 (s, 3H); ^13^C NMR (100 MHz, CDCl_3_) δ 160.2, 149.4, 146.7, 138.8, 131.3, 130.7, 130.5, 128.7, 128.5, 128.4, 128.3, 126.5, 125.9, 118.7, 116.5, 114.8, 55.6, 20.0; *m/z* (ESI^+^) 316.1 ([M+H]^+^, 100%); HRMS (ESI^+^) C_20_H_18_N_3_O^+^ ([M+H]^+^) requires 316.1444; found 316.1442.

**Synthesis of OXS007417 (2)**

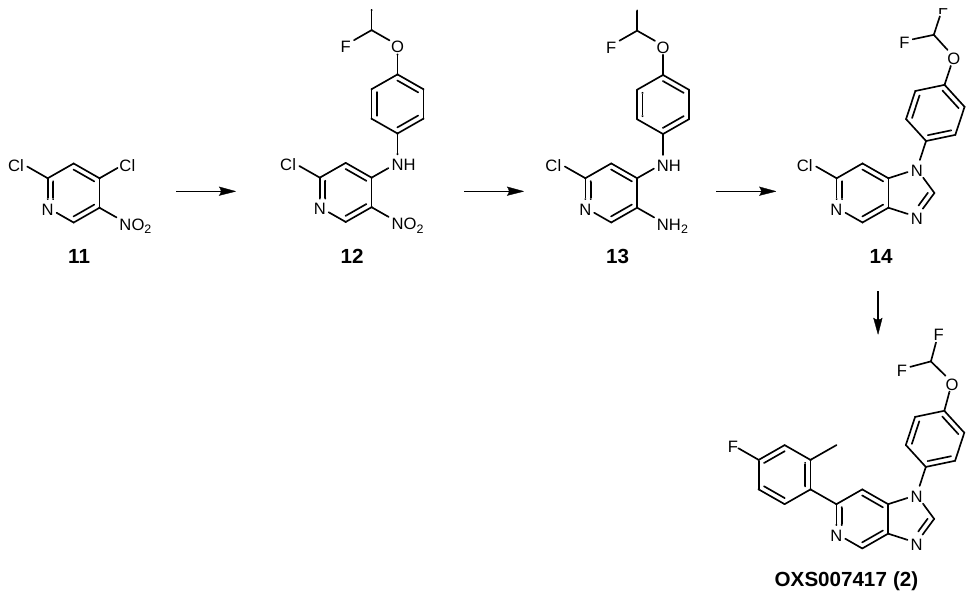

**2-Chloro-*N*-(4-(difluoromethoxy)phenyl)-5-nitropyridin-4-amine (12)**

To a solution of 2,4-dichloro-5-nitro-pyridine **11** (10.0 g, 51.8 mmol) in MeCN (150 mL) was added 4-(difluoromethoxy)aniline (6.29 mL, 51.8 mmol), followed by triethylamine (14.4 mL, 104 mmol). The mixture was stirred at room temperature for 2 days, then the mixture was evaporated to dryness. The residue was dissolved in EtOAc and washed with water, followed by brine, then dried over MgSO_4_ and concentrated *in vacuo* to give **12** (14.9 g, 47.1 mmol, 91%) as a gold solid. ^1^H NMR (CDCl_3_, 400 MHz) δ 9.58 (bs, 1H), 9.10 (s, 1H), 7.32–7.26 (m, 4H), 6.85 (s, 1H), 6.57 (t, *J* = 73.2 Hz, 1H); ^13^C NMR (CDCl_3_, 100 MHz) δ 156.9, 150.3 (t, *J* = 3 Hz), 149.5, 149.0, 133.2, 129.8, 127.4, 121.7, 115.6 (t, *J* = 262 Hz), 108.1; *m/z* (ESI^+^) 316.0 ([M+H]^+^, 100 %).

**6-Chloro-*N*^4^-(4-(difluoromethoxy)phenyl)pyridine-3,4-diamine (13)**

Nitro **12** (14.9 g, 47.1 mmol) was dissolved in IMS (125 mL) and water (250 mL), and ammonium chloride (10.1 g, 189 mmol) and iron (13.2 g, 236 mmol) were added and the mixture heated to 80 °C overnight. The mixture was cooled, diluted with CH_2_Cl_2_ and filtered through Celite®. The organic phase was washed with water then brine, dried over anhydrous MgSO_4_ and concentrated *in vacuo* to give amine **13** (14.7 g, 46.4 mmol, 98%) as a red solid, which was used in the following step without further purification. ^1^H NMR (CDCl_3_, 400 MHz) δ 7.79 (s, 1H), 7.17–7.11 (m, 4H), 6.86 (s, 1H), 6.51 (t, *J* = 74 Hz, 1H), 6.06 (bs, 1H), 3.29 (bs, 2H); ^13^C NMR (CDCl_3_, 100 MHz) δ 147.3 (t, *J* = 3 Hz), 144.2, 144.0, 138.0, 137.0, 129.6, 123.0, 121.3, 115.9 (t, *J* = 261 Hz), 107.0; *m/z* (ESI^+^) 286.0 ([M+H]^+^, 100%).

**6-Chloro-1-(4-(difluoromethoxy)phenyl)-1*H*-imidazo[4,5-*c*]pyridine (14)**

To a suspension of diamine **13** (10.6 g, 37.1 mmol) in diethoxymethoxyethane (110 g, 742 mmol), was added formic acid (1.71 g, 37.1 mmol) and the mixture heated to 100 °C overnight. The mixture was evaporated to dryness, saturated aqueous NaHCO_3_ was added, and the resulting precipitate was filtered, washed with water, followed by 1:1 Et_2_O/hexane, and concentrated *in vacuo* to give imidazopyridine **14** (8.90 g, 30.1 mmol, 81%) as a light brown solid. ^1^H NMR (CDCl_3_, 400 MHz) δ 8.94 (s, 1H), 8.13 (s, 1H), 7.51–7.39 (m, 5H), 6.63 (t, *J* = 72.8 Hz, 1H); ^13^C NMR (CDCl_3_, 100 MHz) δ 151.2 (t, *J* = 3 Hz), 145.1, 144.7, 142.8, 140.8, 140.7, 132.1, 125.8, 121.9, 115.5 (t, *J* = 263 Hz), 105.6; *m/z* (ESI^+^) 296.0 ([M+H]^+^, 100%).

**1-(4-(Difluoromethoxy)phenyl)-6-(4-fluoro-2-methylphenyl)-1*H*-imidazo[4,5-*c*]pyridine (OXS007417, 2)**

Chloride **14** (6.50 g, 22.0 mmol), (4-fluoro-2-methyl-phenyl)boronic acid (4.06 g, 26.4 mmol) and K_2_CO_3_ (6.08 g, 44.0 mmol) were dissolved in DME (50 mL) and water (25 mL). Nitrogen was bubbled though the solution for 10 min, then Pd(dppf)Cl_2_ (0.804 g, 1.10 mmol) was added and nitrogen bubbled for a further 5 min. The mixture was heated to 80 °C overnight. The mixture was cooled, diluted with EtOAc and filtered through Celite®. The aqueous phase was extracted with EtOAc and the combined organics were dried over anhydrous MgSO_4_ and concentrated *in vacuo*. The crude residue was purified using flash column chromatography (0% to 60% EtOAc in hexane), and the resulting solid triturated with hexane and dried to give OXS007417 **2** (3.10 g, 8.39 mmol, 38%) as an off-white solid. ^1^H NMR (500 MHz, CDCl_3_) δ 9.25 (d, *J* = 1.1 Hz, 1H), 8.17 (s, 1H), 7.55–7.50 (m, 2H), 7.46 (d, *J* = 1.1 Hz, 1H), 7.37 (dd, *J* = 8.5, 5.5 Hz, 3H), 7.04–6.90 (m, 2H), 6.60 (t, *J* = 72.8 Hz, 1H), 2.35 (s, 3H); ^13^C NMR (126 MHz, CDCl_3_) δ 162.6 (d, *J* = 247 Hz), 153.6, 150.9 (t, *J* = 3 Hz), 143.9, 142.9, 139.9, 139.3, 138.8 (d, *J* = 8 Hz), 137.0 (d, *J* = 3 Hz), 132.6, 131.6 (d, *J* = 8 Hz), 125.7, 121.9, 117.4 (d, *J* = 21 Hz), 115.5 (t, *J* = 263 Hz), 112.8 (d, *J* = 21 Hz), 105.7, 20.7; *m/z* (ESI^+^) 370.1 ([M+H]^+^, 100%); HRMS (APCI) C_20_H_15_ON_3_F_3_ ([M+H]^+^) requires 370.1158; found 370.1162.

**Synthesis of OXS007464 (3) and OXS007564 (6)**

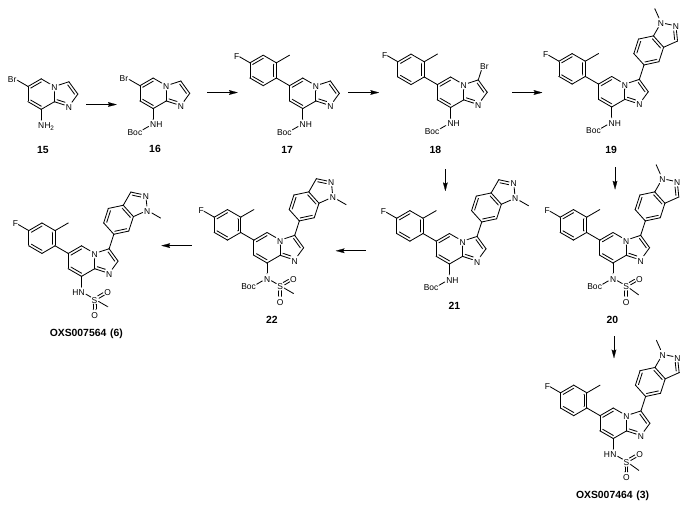

***tert*-Butyl (6-bromoimidazo[1,2-*a*]pyridin-8-yl)carbamate (16)**

To a solution of 6-bromoimidazo[1,2-*a*]pyridin-8-amine **15** (2.05 g, 9.67 mmol) in anhydrous THF (20 mL), sodium bis(trimethylsilyl)amide solution (1 M in THF, 29 mL, 29 mmol) was added dropwise. The solution was stirred at room temperature for 30 min, then Boc anhydride (2.11 g, 9.67 mmol) was added portionwise. The reaction mixture was stirred at room temperature for another 2 h, then quenched with saturated aqueous NH_4_Cl, extracted with three times with CH_2_Cl_2_, dried over anhydrous Na_2_SO_4_ and concentrated. The crude product was purified using flash column chromatography (2% to 25% EtOAc in pentane) to give Boc-protected **16** (2.03 g, 6.50 mmol, 67%) as a green solid. ^1^H NMR (400 MHz, CDCl_3_) δ 7.93–7.89 (m, 2H), 7.90 (s, 1H), 7.49 (d, *J* = 1.2 Hz, 1H), 7.48 (d, *J* = 1.2 Hz, 1H), 1.52 (s, 9H); ^13^C NMR (100 MHz, CDCl_3_) δ 152.3, 137.8, 132.5, 128.3, 119.2, 113.8, 111.4, 108.2, 81.7, 28.3; *m/z* (ESI^+^) 312.0 ([M+H]^+^, 100%); HRMS (ESI^+^) C_12_H_15_N_3_O_2_Br^+^ ([M+H]^+^) requires 312.0342; found 312.0343.

***tert*-Butyl (6-(4-fluoro-2-methylphenyl)imidazo[1,2-*a*]pyridin-8-yl)carbamate (17)**

Pd(PPh_3_)_4_ (259 mg, 0.22 mmol) was added to a stirred solution of **16** (700 mg, 2.24 mmol), (4-fluoro-2-methyl-phenyl)boronic acid (345 mg, 2.24 mmol) and K_3_PO_4_ (1.43 g, 6.73 mmol) in degassed DME/water (3:1, 14 mL) at room temperature. The resultant mixture was heated to 90 °C for 16 h, then cooled to room temperature, diluted with EtOAc, filtered through Celite® and concentrated. Purification using flash column chromatography (20% EtOAc in pentane) gave the aryl-substituted product **17** (741 mg, 2.17 mmol, 97%) as a pale yellow foam. ^1^H NMR (400 MHz, CDCl_3_) δ 7.91 (s, 1H), 7.76 (s, 1H), 7.69 (d, *J* = 1.5 Hz, 1H), 7.58 (d, *J* = 1.2 Hz, 1H), 7.56 (d, *J* = 1.3 Hz, 1H), 7.21 (dd, *J* = 8.4, 5.9 Hz, 1H), 6.98 (dd, *J* = 9.7, 2.7 Hz, 1H), 6.92 (td, *J* = 8.4, 2.7 Hz, 1H), 2.30 (s, 3H), 1.53 (s, 9H); ^13^C NMR (101 MHz, CDCl_3_) δ 162.3 (d, *J* = 246.4 Hz), 152.5, 138.6 (d, *J* = 8.0 Hz), 138.3, 133.8 (d, *J* = 3.2 Hz), 132.3, 131.4 (d, *J* = 8.4 Hz), 127.2, 127.1, 118.0, 116.9 (d, *J* = 21.2 Hz), 113.8, 112.6 (d, *J* = 21.2 Hz), 110.6, 81.1, 28.2, 20.6; HRMS (ESI^+^) C_19_H_21_FN_3_O_2_^+^ ([M+H]^+^) requires 342.1612; found 342.1612.

***tert*-Butyl (3-bromo-6-(4-fluoro-2-methylphenyl)imidazo[1,2-*a*]pyridin-8-yl)carbamate (18)**

N-bromosuccinimide (334 mg, 1.87 mmol) was added to a stirred solution of **17** (640 mg, 1.87 mmol) in THF (4 mL) at room temperature. The resultant mixture was stirred for 2 h, then concentrated. Purification using flash column chromatography (3% EtOAc in pentane) gave the brominated product **18**; (619 mg, 1.47 mmol, 79%) as a pale yellow foam. ^1^H NMR (400 MHz, CDCl_3_) δ 7.90–7.82 (m, 2H), 7.69 (d, *J* = 1.5 Hz, 1H), 7.55 (s, 1H), 7.24 (dd, *J* = 8.4, 5.9 Hz, 1H), 7.00 (dd, *J* = 9.7, 2.7 Hz, 1H), 6.98–6.91 (m, 1H), 2.31 (s, 3H), 1.53 (s, 9H); ^13^C NMR (101 MHz, CDCl_3_) δ 162.5 (d, *J* = 246.7 Hz), 152.4, 138.7 (d, *J* = 8.1 Hz), 138.5, 133.7 (d, *J* = 3.2 Hz), 132.4, 131.5 (d, *J* = 8.4 Hz), 128.0, 127.3, 117.0 (d, *J* = 21.2 Hz), 115.9, 112.7 (d, *J* = 21.2 Hz), 110.9, 96.3, 81.4, 28.2, 20.6 (d, *J* = 1.6 Hz); HRMS (ESI^+^) C_19_H_20_BrFN_3_O_2_^+^ ([M+H]^+^) requires 420.0717; found 420.0720.

***tert*-Butyl (6-(4-fluoro-2-methylphenyl)-3-(1-methyl-1*H*-indazol-5-yl)imidazo[1,2-*a*]pyridin-8-yl)carbamate (19)**

Pd(dppf)Cl_2_ (71 mg, 0.10 mmol) was added to a stirred solution of **18** (580 mg, 1.38 mmol), (1-methylindazol-5-yl)boronic acid (291 mg, 1.66 mmol) and K_2_CO_3_ (572 mg, 4.14 mmol) in degassed DME/water (3:1, 9 mL) at room temperature. The resultant mixture was heated to 70 °C for 16 h, then cooled to room temperature, diluted with EtOAc, filtered through Celite® and concentrated. Purification using flash column chromatography (30% EtOAc in pentane) gave the aryl-substituted product **19** (641 mg, 1.36 mmol, 99%) as a pale yellow foam. ^1^H NMR (400 MHz, CDCl_3_) δ 8.05 (d, *J* = 0.8 Hz, 1H), 7.99 (s, 1H), 7.91 (t, *J* = 1.2 Hz, 1H), 7.85–7.80 (m, 2H), 7.63 (s, 1H), 7.57 (dd, *J* = 8.7, 1.5 Hz, 1H), 7.53 (dt, *J* = 8.7, 0.9 Hz, 1H), 7.20 (dd, *J* = 8.4, 5.9 Hz, 1H), 6.96 (dd, *J* = 9.7, 2.7 Hz, 1H), 6.89 (td, *J* = 8.4, 2.7 Hz, 1H), 4.13 (s, 3H), 2.31 (s, 3H), 1.55 (s, 9H); ^13^C NMR (101 MHz, CDCl_3_) δ 162.3 (d, *J* = 246.5 Hz), 152.6, 139.5, 138.6 (d, *J* = 5.3 Hz), 138.5, 134.1 (d, *J* = 3.2 Hz), 133.1, 131.5, 131.4, 131.0, 127.5, 127.4 (d, *J* = 13.6 Hz), 126.9, 124.4, 121.3, 121.1, 116.9 (d, *J* = 21.2 Hz), 115.6, 112.6 (d, *J* = 21.1 Hz), 110.5, 109.9, 81.1, 35.7, 28.2, 20.7 (d, *J* = 1.5 Hz); HRMS (ESI^+^) C_27_H_27_FN_5_O_2_^+^ ([M+H]^+^) requires 472.2143; found 472.2135.

***tert*-Butyl (6-(4-fluoro-2-methylphenyl)-3-(1-methyl-1*H*-indazol-5-yl)imidazo[1,2-*a*]pyridin-8-yl)(methylsulfonyl)carbamate (20)**

Sodium hydride (60% w/w dispersion in mineral oil, 407 mg, 10.2 mmol) was added to a stirred solution of **19** (1.60 g, 3.39 mmol) in THF (150 mL) at room temperature. The resultant mixture was stirred for 30 min before the addition of methanesulfonyl chloride (466 mg, 4.07 mmol). The resultant mixture was stirred at room temperature for 16 h, before the addition of water. The aqueous layer was extracted with EtOAc, dried over anhydrous MgSO_4_ and concentrated. Purification using flash column chromatography (0% to 60% EtOAc in pentane) gave mesyl **20** (1.70 g, 3.09 mmol, 91%) as an orange oil. ^1^H NMR (CDCl_3_, 400 MHz) δ 8.15 (d, *J* = 1.2 Hz, 1H), 8.06 (s, 1H), 7.90 (m, 1H), 7.70 (s, 1H), 7.55 (d, *J* = 0.8 H z, 2H), 7.30 (d, *J* = 1.6 Hz, 1H), 7.22 (dd, *J* = 5.6, 8.4 H , 1H), 6.99 (dd, *J* = 2.4, 9.6 Hz, 1H), 6.92 (td, *J* = 2.8, 8.4 Hz, 1H), 4.13 (s, 3H), 3.79 (s, 3H), 2.30 (s, 3H), 1.51 (s, 9H); *m/z* (ESI^+^) 550.1 ([M+H]^+^, 100%).

***N*-(6-(4-Fluoro-2-methylphenyl)-3-(1-methyl-1*H*-indazol-5-yl)imidazo[1,2-a]pyridin-8-yl)methanesulfonamide (OXS007464, 3)**

A solution of **20** (130 mg, 0.25 mmol) in 2 M HCl in Et_2_O (4.0 mL) was stirred at room temperature for 16 h, before concentration *in vacuo*. The residue was dissolved in EtOAc, and basified with saturated aqueous NaHCO_3_. The aqueous layer was extracted with EtOAc, dried over anhydrous MgSO_4_ and concentrated. Purification using flash column chromatography (0% to 100% EtOAc in pentane) gave the product OXS007464 **3** (54 mg, 0.12 mmol, 50%) as an off-white solid. ^1^H NMR (400 MHz, DMSO-*d*_6_) δ 8.12 (s, 1H), 8.09 (s, 1H), 8.07 (s, 1H), 7.81 (s, 1H), 7.80 (d, *J* = 8.8 Hz, 1H), 7.69 (dd, *J* = 8.7, 1.3 Hz, 1H), 7.38 (dd, *J* = 8.4, 6.1 Hz, 1H), 7.18 (dd, *J* = 10.1, 2.6 Hz, 1H), 7.14 (d, *J* = 1.4 Hz, 1H), 7.08 (td, *J* = 8.5, 2.7 Hz, 1H), 4.09 (s, 3H), 3.28 (s, 3H), 2.29 (s, 3H); ^13^C NMR (101 MHz, DMSO-*d*_6_) δ 161.7 (d, *J* = 244 Hz), 139.3, 139.1, 138.6 (d, *J* = 8 Hz), 133.6 (d, *J* = 3 Hz), 132.8, 131.6 (d, *J* = 8 Hz), 131.5, 127.3, 126.6, 126.5, 125.4, 123.8, 120.5, 120.4, 118.1, 116.8 (d, *J* = 21 Hz), 115.1, 112.6 (d, *J* = 21 Hz), 110.6, 40.9, 35.4, 20.0; *m/z* (ESI^+^) 450.1 ([M+H]^+^, 100%); HRMS (ESI^+^) C_23_H_21_O_2_N_5_FS^+^ ([M+H]^+^) requires 450.1395, found 450.1390.

***tert*-Butyl (6-(4-fluoro-2-methylphenyl)-3-(1-methyl-1*H*-indazol-6-yl)imidazo[1,2-*a*]pyridin-8-yl)carbamate (21)**

Bromide **18** (250 mg, 0.59 mmol), (1-methylindazol-6-yl)boronic acid (126 mg, 0.716 mmol) and potassium phosphate tribasic (379 mg, 1.79 mmol) were dissolved in degassed Dimethoxyethane /water (4:1, 5 mL). Pd(dppf)Cl_2_ (22 mg, 0.030 mmol) was added and the reaction heated at 80 °C overnight. The mixture was cooled to room temperature and concentrated *in vacuo*. The residue was purified using flash column chromatography (0% to 40% EtOAc in hexane) to give aryl-substituted **21** (190 mg, 0.403 mmol, 68%) as an orange solid. ^1^H NMR (CDCl_3_, 400 MHz) δ 8.03 (s, 1H), 7.96 (bs, 1H), 7.91 (d, *J* = 1.6 Hz, 1H), 7.85 (m, 2H), 7.69 (s, 1H), 7.56 (s, 1H), 7.34 (dd, *J* = 1.2, 8.4 Hz, 1H), 7.21 (dd, *J* = 5.6, 8.4 Hz, 1H), 6.95–6.98 (m, 1H), 6.92-6.87 (m, 1H), 4.12 (s, 3H), 2.32 (s, 3H), 1.55 (s, 9H); *m/z* (ESI^+^) 472.2 ([M+H]^+^, 100%).

***tert*-Butyl (6-(4-fluoro-2-methylphenyl)-3-(1-methyl-1*H*-indazol-6-yl)imidazo[1,2-*a*]pyridin-8-yl)(methylsulfonyl)carbamate (22)**

To **21** (182 mg, 0.382 mmol) in THF (4 mL) was added sodium hydride (60% w/w dispersion in mineral oil, 44 mg, 1.2 mmol) and stirred for 15 min. Methanesulfonyl chloride (53 mg, 0.46 mmol) was added and the reaction stirred overnight. Two further equivalents of sodium hydride and methanesulfonyl chloride were added to complete the reaction. The reaction mixture was quenched with ice water, extracted with CH_2_Cl_2_, dried over anhydrous MgSO_4_ and concentrated *in vacuo*. The crude product was purified using flash column chromatography (30% to 80% EtOAc in hexane) to give the mesyl product **22** (130 mg, 0.237 mmol, 61%) as an orange solid. ^1^H NMR (CDCl_3_, 400 MHz) δ 8.23 (d, *J* = 1.6 Hz, 1H), 8.04 (d, *J* = 1.2 Hz,1H), 7.86 (dd, *J* = 0.4, 8.4 Hz, 1H), 7.75 (s, 1H), 7.55 (d, *J* = 1.2 Hz, 1H), 7.33–7.31 (m, 2H), 7.23 (dd, *J* = 2.0, 8.4 Hz, 1H), 7.00 (dd, *J* = 2.4, 9.6 Hz, 1H), 6.94 (dt, *J* = 2.8, 8.4 Hz, 1H), 4.12 (s, 3H), 3.79 (s, 3H), 2.31 (s, 3H), 1.52 (s, 9H); *m/z* (ESI^+^) 550.1 ([M+H]^+^, 100%).

***N*-(6-(4-Fluoro-2-methylphenyl)-3-(1-methyl-1*H*-indazol-6-yl)imidazo[1,2-*a*]pyridin-8-yl)methanesulfonamide (OXS007564, 6)**

To a solution of **22** (120 mg, 0.218 mmol) in 4 M HCl in dioxane (4.0 mL) was added 3 drops of MeOH and the reaction mixture stirred at room temperature for 3 days. The reaction mixture was then concentrated *in vacuo*, basified with saturated aqueous NaHCO_3_ and extracted with EtOAc. The combined organic layers were concentrated and purified using flash column chromatography (10% to 100% EtOAc in hexane), then triturated with Et_2_O and dried to give **OXS007564 (6)** (65 mg, 0.14 mmol, 66%) as an off-white solid. ^1^H NMR (CDCl_3_, 400 MHz) δ 8.04 (d, *J* = 1.2 Hz, 1H), 8.01 (d, *J* = 1.2 Hz, 1H), 7.88–7.85 (m, 1H), 7.74 (s, 1H), 7.56 (s, 1H), 7.35–7.31 (m, 2H), 7.21 (dd, *J* = 5.6, 8.4 Hz, 1H), 7.02–6.99 (m, 1H), 6.94 (td, *J* = 2.8, 8.4 Hz, 1H), 4.12 (s, 3H), 3.16 (s, 3H), 2.32 (s, 3H); ^13^C NMR (CDCl_3_, 100 MHz) δ 162.7 (d, *J* = 248 Hz), 140.3, 139.3, 138.6 (d, *J* = 8 Hz), ), 133.3 (d, *J* = 3 Hz), 133.1, 132.3, 131.6 (d, *J* = 8 Hz), 128.3, 127.3, 126.6, 126.5, 124.1, 122.4, 120.7, 118.1, 117.5 (d, *J* = 21 Hz), 113.3, 113.2, 113.0, 109.1, 40.2, 35.9, 20.8; *m/z* (ESI^+^) 450.1 ([M+H]^+^, 100%). HRMS (ESI^+^) C_23_H_21_O_2_N_5_FS^+^ ([M+H]^+^) requires 450.1395, found 450.1394.

**Synthesis of diazirine 29**

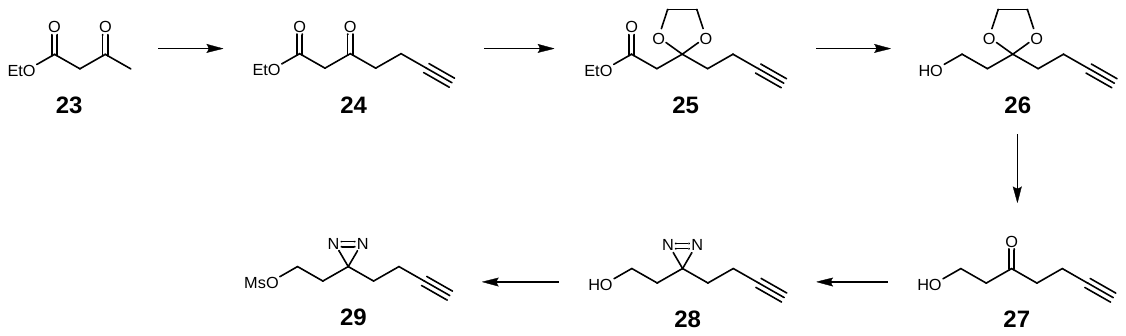

**Ethyl 3-oxohept-6-ynoate (24)**

Lithium diisopropylamide (2 M in THF/heptane/ethylbenzene, 16.9 mL, 33.8 mmol) was added dropwise to a solution of ethyl acetoacetate **23** (1.9 mL, 15 mmol) in THF (15.4 mL) at 0 °C under argon. Propargyl bromide (80% w/w, 1.7 mL, 15 mmol) was added dropwise and the reaction was stirred at room temperature for 4 h. The mixture was neutralised with saturated aqueous NH_4_Cl, extracted three times with EtOAc, dried over anhydrous Na_2_SO_4_ and concentrated *in vacuo*. The crude product was purified using flash column chromatography (5% EtOAc in pentane) to give the product **24** (1.25 g, 7.39 mmol, 48%) as a yellow oil. ^1^H NMR (CDCl_3_, 400 MHz) δ 4.13 (q, *J* = 7.2 Hz, 2H), 3.41 (s, 2H), 2.77 (t, *J* = 7.1 Hz, 2H), 2.40 (td, *J* = 7.0, 2.7 Hz, 2H), 1.91 (t, *J* = 2.7 Hz, 1H), 1.21 (t, *J* = 7.1 Hz, 3H); ^13^C NMR (CDCl_3_, 100 MHz) δ 200.6, 166.9, 82.5, 69.0, 61.4, 49.1, 41.5, 14.0, 12.7; *m/z* (ESI^+^) 191.0 ([M+Na]^+^, 100%); HRMS (ESI^+^) C_9_H_13_O_3_^+^ ([M+H]^+^) requires 169.0859, found 169.0861. NMR was in agreement with the reported values.^2^

**Ethyl 2-(2-(but-3-yn-1-yl)-1,3-dioxolan-2-yl)acetate (25)**

BF_3_·OEt_2_ (800 μL, 6.27 mmol) was added dropwise to a solution of **24** (703 mg, 4.18 mmol) and ethylene glycol (900 μL, 16.7 mmol) in CH_2_Cl_2_ (8.4 mL) at 0 °C under nitrogen. The reaction was stirred at 0 °C for 1 h, before warming to room temperature and stirring overnight. The mixture was cooled to 0 °C and water was added dropwise. The crude product was extracted three times with CH_2_Cl_2_, washed with brine, dried over anhydrous Na_2_SO_4_ and concentrated *in vacuo*. The crude product was purified using flash column chromatography (5% EtOAc in pentane) to give dioxolane **25** (560 mg, 2.63 mmol, 63%) as a yellow oil. ^1^H NMR (CDCl_3_, 400 MHz) δ 4.15 (q, *J* = 7.1 Hz, 2H), 4.04–3.92 (m, 4H), 2.65 (s, 2H), 2.33–2.26 (m, 2H), 2.15–2.09 (m, 2H), 1.93 (t, *J* = 2.7 Hz, 1H), 1.26 (t, *J* = 7.1 Hz, 3H); ^13^C NMR (CDCl_3_, 100 MHz) δ 169.3, 108.5, 84.1, 68.2, 65.4, 60.8, 42.8, 36.6, 14.3, 13.0; *m/z* (ESI^+^) 235.0 ([M+Na]^+^, 100%); HRMS (ESI^+^) C_11_H_16_O_4_Na^+^ ([M+Na]^+^) requires 235.0941, found 235.0941. NMR was in agreement with the reported values.^3^

**2-(2-(But-3-yn-1-yl)-1,3-dioxolan-2-yl)ethan-1-ol (26)**

A solution of **25** (535 mg, 2.52 mmol) in Et_2_O (12.6 mL) was added dropwise to a solution of LiAlH_4_ (1 M in THF, 2.5 mL, 2.5 mmol) in Et_2_O (15 mL) at 0 °C under nitrogen. The reaction was stirred at room temperature for 30 min. The mixture was quenched with a few drops of aqueous 1 M NaOH and water, filtered and concentrated *in vacuo* to give alcohol **26** (314 mg, 1.84 mmol, 73%) as a yellow oil, which was used in the following step without further purification. ^1^H NMR (CDCl_3_, 400 MHz) δ 4.07–3.93 (m, 4H), 3.76 (q, *J* = 5.7 Hz, 2H), 2.65 (t, *J* = 5.7 Hz, 1H), 2.31–2.23 (m, 2H), 1.96–1.91 (m, 5H); ^13^C NMR (CDCl_3_, 100 MHz) δ 111.2, 84.1, 68.4, 65.1, 58.9, 38.4, 36.1, 13.3; *m/z* (ESI^+^) 235.0 ([M+Na]^+^, 100%); HRMS (ESI^+^) C_9_H_14_O_3_Na^+^ ([M+Na]^+^) requires 193.0835, found 193.0837. NMR was in agreement with the reported values.^3^

**1-Hydroxyhept-6-yn-3-one (27)**

Aqueous HCl (5 M, 7.2 mL, 36 mmol) was added to a solution of **26** (1.44 g, 8.46 mmol) in THF (21.7 mL), and the mixture was stirred at room temperature for 3 h. The mixture was diluted with water, neutralised with aqueous saturated NaHCO_3_ and extracted three times with EtOAc. The organic layer was dried over anhydrous Na_2_SO_4_ and concentrated *in vacuo* to give deprotected ketone **27** (990 mg, 7.85 mmol, 93%) as a yellow oil. ^1^H NMR (CDCl_3_, 400 MHz) δ 3.81 (t, *J* = 5.4 Hz, 2H), 2.70–2.56 (m, 4H), 2.41 (td, *J* = 7.2, 2.7 Hz, 2H), 1.89 (t, *J* = 2.7 Hz, 1H); ^13^C NMR (CDCl_3_, 100 MHz) δ 209.2, 82.9, 69.1, 57.9, 44.7, 41.9, 13.0; HRMS (ESI^+^) C_7_H_10_O_2_Na^+^ ([M+Na]^+^) requires 149.0573, found 149.0573. NMR was in agreement with the reported values.^4^

**2-(3-(But-3-yn-1-yl)-3*H*-diazirin-3-yl)ethan-1-ol (28)**

Ammonia solution (7 M in MeOH, 3.4 mL, 24 mmol) was added to ketone **27** (200 mg, 1.59 mmol) and stirred for 4.5 h at -10 °C. A solution of hydroxylamine-*O*-sulfonic acid (233 mg, 2.06 mmol) in dry MeOH (1.2 mL) was then added dropwise at 0 °C and stirred for 1 h. The reaction was warmed to room temperature and stirred overnight. Excess ammonia was removed using a stream of nitrogen, and then the mixture was filtered and the solid washed with dry MeOH. Triethylamine (1.0 mL, 12 mmol) was added to the organic phase and cooled to 0 °C. Iodine crystals (523 mg, 2.06 mmol) were added portion-wise and the mixture was stirred at 0 °C for 1 h. The mixture was extracted with three times with Et_2_O, washed with brine and concentrated *in vacuo*. The crude product was purified using flash column chromatography (20% EtOAc in pentane) to give diazirine **28** (35 mg, 0.25 mmol, 16%) as a pale yellow oil. ^1^H NMR (CDCl_3_, 400 MHz) δ 3.49 (t, *J* = 6.2 Hz, 2H), 2.04 (td, *J* = 7.3, 2.6 Hz, 2H), 2.00 (t, *J* = 2.6 Hz, 1H), 1.74–1.65 (m, 4H); ^13^C NMR (CDCl_3_, 100 MHz) δ 83.0, 69.4, 57.5, 35.7, 32.8, 26.7, 13.4; *m/z* not found. NMR was in agreement with the reported values.^4^

**2-(3-(But-3-yn-1-yl)-3*H*-diazirin-3-yl)ethyl methanesulfonate (29)**

Mesyl chloride (23 μL, 0.29 mmol) was added dropwise to a solution of **28** (40 mg, 0.29 mmol) and triethylamine (49 μL, 0.35 mmol) in THF (1.5 mL) at 0°C under argon, and stirred at room temperature for 2 h. The mixture was diluted with water and extracted with three times with CH_2_Cl_2_. The organic layer was washed with brine, dried with anhydrous Na_2_SO_4_ and concentrated *in vacuo* to give mesyl **29** (53 mg, 0.25 mmol, 85%) as a yellow oil. ^1^H NMR (CDCl_3_, 400 MHz) δ 4.08 (t, *J* = 6.2Hz, 2H), 3.06 (s, 3H), 2.04 (td, *J* = 7.2, 2.4 Hz, 1H), 2.01 (t, *J* = 2.6 Hz, 1H), 1.90 (t, *J* = 6.2 Hz, 2H), 1.70 (t, *J* = 7.3, 2H); ^13^C NMR (CDCl_3_, 100 MHz) δ 82.5, 69.7, 64.1, 37.8, 33.0, 32.4, 26.0, 13.5; HRMS (ESI^+^) C_8_H_12_N_2_O_3_SNa^+^ ([M+Na]^+^) requires 239.0461, found 239.0463.

**Synthesis of probe OXS008450 (4)**

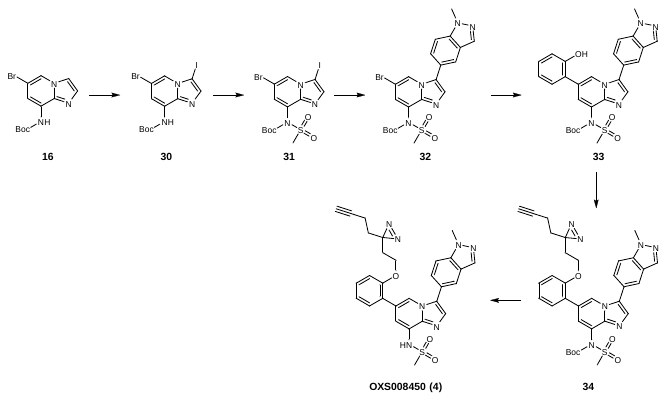

***tert*-Butyl (6-bromo-3-iodoimidazo[1,2-*a*]pyridin-8-yl)carbamate (30)**

To a stirred solution of **16** (12.7 g, 40.7 mmol) in THF (150 ml) was added *N*-iodosuccinimide (9.15 g, 40.7 mmol), and the mixture was stirred at room temperature for 23 h. The mixture was diluted with water and extracted into CH_2_Cl_2_. The combined extracts were washed with aqueous 1 M Na_2_S_2_O_3_ and brine, dried over MgSO_4_ and dried to give a green solid, which was stirred in hexane for 2 h, filtered and dried to give iodide **30** (13.8 g, 31.5 mmol, 78%) as a light green solid. ^1^H NMR (CDCl_3_, 400 MHz) δ 8.02 (bs, 1H), 7.91 (d, *J* = 1.6 Hz, 1H), 7.83 (bs, 1H), 7.55 (s, 1H), 1.54 (s, 9H); ^13^C NMR (CDCl_3_, 100 MHz) δ 152.1, 139.8, 138.9, 128.1, 119.4, 112.1, 109.3, 81.8, 62.1, 28.2; *m/z* (ESI^+^) 437.9 ([M+H]^+^, 100%).

***tert*-Butyl (6-bromo-3-iodoimidazo[1,2-*a*]pyridin-8-yl)(methylsulfonyl)carbamate (31)**

Sodium hydride (60% w/w dispersion in mineral oil, 302 mg, 7.56 mmol) was added portionwise to a solution of iodide **30** (1.10 g, 2.52 mmol) in dry THF (37.6 mL) at 0 °C under nitrogen. Mesyl chloride (214 μL, 2.77 mmol) was added dropwise at 0 °C. The reaction was stirred at room temperature for 1 h, then diluted with water and the crude product extracted three times with EtOAc. The organic layer was dried with anhydrous Na_2_SO_4_ and concentrated *in vacuo*. The crude product was purified by trituration with isopropyl alcohol to give mesyl **31** (955 mg, 1.84 mmol, 73%) as a white solid. ^1^H NMR (CDCl_3_, 400 MHz) δ 8.29 (d, *J* = 1.7 Hz, 1H), 7.66 (s, 1H), 7.46 (d, *J* = 1.7 Hz, 1H), 3.67 (s, 3H), 1.44 (s, 9H); ^13^C NMR (CDCl_3_, 100 MHz) δ 150.6, 144.2, 141.1, 131.2, 127.2, 125.4, 106.9, 85.9, 62.3, 42.1, 27.9; *m/z* (ESI^+^) 515.5 ([M+H]^+^, 100%); HRMS (ESI^+^) C_13_H_16_^79^BrIN_3_O_4_S^+^ ([M^79^Br+H]^+^) requires 515.9084, found 515.9084.

**t*ert*-Butyl (6-bromo-3-(1-methyl*-1H-*indazol-5-yl)imidazo[1,2-*a*]pyridin-8-yl)-(methylsulfonyl)carbamate (32)**

Pd(dppf)Cl_2_ (7 mg, 0.01 mmol) was added to a stirred solution of 31 (98 mg, 0.19 mmol), (1-methylindazol-5-yl)boronic acid (37 mg, 0.21 mmol) and K_2_CO_3_ (80 mg, 0.58 mmol) in degassed dioxane/water (4:1, 2.0 mL) at room temperature under argon. The resultant mixture was heated to 50 °C for 3 h, then cooled to room temperature, diluted with EtOAc, filtered through Celite® and concentrated. Purification using flash column chromatography (30% to 80% EtOAc in pentane) gave the product 32 (76 mg, 0.15 mmol, 77%) as a yellow solid. ^1^H NMR (CDCl_3_, 400 MHz) δ 8.36 (d, *J* = 1.7 Hz, 1H), 8.09 (d, *J* = 1.0 Hz, 1H), 7.87 (dd, *J* = 1.6, 0.9 Hz, 1H), 7.65 (s, 1H), 7.58 (dt, *J* = 8.7, 1.0 Hz, 1H), 7.50 (dd, *J* = 8.7, 1.6 Hz, 1H), 7.41 (d, *J* = 1.7 Hz, 1H), 4.15 (s, 3H), 3.75 (s, 3H), 1.50 (s, 9H); ^13^C NMR (CDCl_3_, 100 MHz) δ 150.8, 142.1, 139.8, 133.4, 133.2, 130.5, 127.5, 126.9, 125.7, 124.6, 124.4, 121.7, 120.4, 110.4, 106.1, 85.7, 42.2, 35.9, 28.0; *m/z* (ESI^+^) 520.6 ([M+H]^+^, 100%); HRMS (ESI^+^) C_21_H_23_^79^BrN_5_O_4_S^+^ ([M^79^Br+H]^+^) requires 520.0649, found 520.0646.

***tert*-Butyl (6-(2-hydroxyphenyl)-3-(1-methyl-1H-indazol-5-yl)imidazo[1,2-*a*]-pyridin-8-yl)(methylsulfonyl)carbamate (33)**

Pd(dppf)Cl_2_ (34 mg, 0.046 mmol) was added to a stirred solution of **32** (398 mg, 0.764 mmol), 2-hydroxyphenylboronic acid (105 mg, 0.764 mmol) and K_3_PO_4_ (568 mg, 2.68 mmol) in degassed DME/water (2.3:1, 5 mL) at room temperature under argon. The resultant mixture was heated to 60 °C for 1 h, then cooled to room temperature, diluted with EtOAc, filtered through Celite® and concentrated. Purification using flash column chromatography (80% EtOAc in pentane) gave the product **33** (389 mg, 0.729 mmol, 95%) as an orange solid. ^1^H NMR (CDCl_3_, 400 MHz) δ 8.65 (d, *J* = 1.5 Hz, 1H), 7.92 (d, *J* = 0.6 Hz, 1H), 7.84 (dd, *J* = 1.2, 0.3 Hz, 1H), 7.64 (s, 1H), 7.60 (d, *J* = 1.4 Hz, 1H), 7.52–7.49 (m, 2H), 7.33 (dd, *J* = 7.6, 1.7 Hz, 1H), 7.20 (ddd, *J* = 8.0, 7.4, 1.7 Hz, 1H), 6.99–6.90 (m, 2H), 4.08 (s, 3H), 3.71 (s, 3H), 1.49 (s, 9H); ^13^C NMR (CDCl_3_, 100 MHz) δ 153.8, 151.1, 142.5, 139.7, 133.2, 132.5, 130.5, 130.1, 129.8, 127.4, 127.4, 124.6, 124.5, 124.2, 123.4, 123.2, 121.4, 121.1, 121.0, 116.8, 110.2, 85.7, 42.4, 35.8, 28.0; *m/z* (ESI^+^) 534.2 ([M+H]^+^, 100%); HRMS (ESI^+^) C_27_H_28_N_5_O_5_S^+^ ([M+H]^+^) requires 534.1806, found 534.1799.

***tert*-Butyl (3-(1-methyl-1*H*-indazol-5-yl)-6-(2-(2-(3-(prop-2-yn-1-yl)-3*H*-diazirin-3-yl)ethoxy)phenyl)imidazo[1,2-*a*]pyridin-8-yl)(methylsulfonyl)carbamate (34)**

A solution of mesyl **29** (24 mg, 0.11 mmol) in dry DMF (0.5 mL) was added dropwise to a solution of phenol **33** (50 mg, 0.094 mmol) and K_2_CO_3_ (16 mg, 0.11 mmol) in dry Dimethylformamide (1.0 mL) at 0 °C under nitrogen, and the mixture was stirred at 60 °C for 30 h. The mixture was diluted with water (40 mL), extracted with three times with EtOAc, dried over anhydrous Na_2_SO_4_ and concentrated *in vacuo* to give the product **34** (36 mg, 0.056 mmol, 60%) as a white solid, which was used in the following step without further purification. ^1^H NMR (CDCl_3_, 400 MHz) δ 8.47 (d, *J* = 1.5 Hz, 1H), 8.06 (s, 1H), 7.92 (s, 1H), 7.66 (s, 1H), 7.63 (d, *J* = 1.5 Hz, 1H), 7.61–7.50 (m, 2H), 7.36–7.29 (m, 2H), 7.02 (td, *J* = 7.5, 0.7 Hz, 1H), 6.90 (dd, *J* = 8.4, 0.7 Hz, 1H), 4.14 (s, 3H), 3.78 (s, 3H), 3.74 (t, *J* = 6.3 Hz, 1H), 1.87–1.75 (m, 5H), 1.52 (s, 9H), 1.46 (t, *J* = 7.4 Hz, 1H); ^13^C NMR (CDCl_3_, 100 MHz) δ 155.5, 151.1, 142.4, 139.6, 133.2, 132.6, 130.7, 130.4, 129.7, 127.22, 127.16, 125.8, 124.5, 124.2, 123.47, 123.45, 121.44, 121.36, 121.3, 112.2, 110.0, 85.1, 82.7, 69.0, 62.9, 42.1, 35.8, 32.4, 32.3, 27.9, 13.0; HRMS (ESI^+^) C_34_H_36_N_7_O_5_S^+^ ([M+H]^+^) requires 654.2493, found 654.2481.

***N*-(3-(1-Methyl-1*H*-indazol-5-yl)-6-(2-(2-(3-(prop-2-yn-1-yl)-3*H*-diazirin-3-yl)-ethoxy)phenyl)imidazo[1,2-*a*]pyridin-8-yl)methanesulfonamide (OXS008450, 4)**

A solution of Boc-protected **34** (32 mg, 0.048 mmol) and trifluoroacetic acid (185 μL, 2.42 mmol) in CH_2_Cl_2_ (1.0 mL) was stirred at room temperature under nitrogen for 3 h. The reaction mixture was neutralised with saturated NaHCO_3_, extracted with three times with CH_2_Cl_2_ and washed with saturated aqueous NH_4_Cl. The organic layer was dried with anhydrous Na_2_SO_4_, filtered and concentrated *in vacuo* to obtain the deprotected product OXS008450 **4** (24 mg, 0.044 mmol, 91%) as a beige solid. ^1^H NMR (CDCl_3_, 400 MHz) δ 8.26 (d, *J* = 1.0 Hz, 1H), 8.06 (s, 1H), 7.93 (s, 1H), 7.67 (s, 1H), 7.64 (d, *J* = 1.0 Hz, 1H), 7.61–7.51 (m, 2H), 7.38–7.29 (m, 2H), 7.03 (td, *J* = 7.5, 0.6 Hz, 1H), 6.93 (dd, *J* = 8.2, 0.7 Hz, 1H), 4.14 (s, 3H), 3.83 (t, *J* = 6.2 Hz, 2H), 3.17 (s, 3H), 1.82–1.70 (m, 5H), 1.45 (t, *J* = 7.5 Hz, 2H); ^13^C NMR (CDCl_3_, 126 MHz) δ 155.7, 139.7, 139.3, 133.3, 131.6, 131.0, 129.8, 128.2, 127.2, 126.5, 125.9, 124.7, 124.6, 121.5, 121.4, 121.2, 118.9, 114.5, 112.2, 110.1, 82.5, 69.1, 63.1, 40.1, 35.9, 32.8, 32.5, 26.6, 13.1; *m/z* (ESI^+^) 554.2 ([M+H]^+^, 100%); HRMS (ESI^+^) C_29_H_28_N_7_O_3_S^+^ ([M+H]^+^) requires 554.1969, found 554.1969.

**Synthesis of OXS008255**

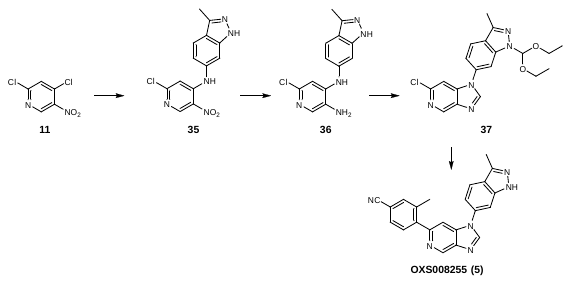

***N*-(2-Chloro-5-nitropyridin-4-yl)-3-methyl-1*H*-indazol-6-amine (35)**

2,4-Dichloro-5-nitro-pyridine **11** (500 mg, 2.59 mmol) and 3-methyl-1*H*-indazol-6-amine (381 mg, 2.59 mmol) were dissolved in MeCN (10 mL), triethylamine (722 µL, 5.18 mmol) was added and the mixture stirred at room temperature for 3 days. The mixture was concentrated *in vacuo*, water was added and the resulting precipitate filtered off, and washed with water and Et_2_O to give the product **35** (918 mg, 3.02 mmol, quant.) as an orange solid. ^1^H NMR (DMSO-*d*_6_, 400 MHz) δ 12.74 (bs, 1H), 10.02 (s, 1H), 8.97 (s, 1H), 7.81 (d, *J* = 8.8 Hz, 1H), 7.46 (s, 1H), 7.06 (dd, *J* = 8.4, 1.6 Hz, 1H), 6.78 (s, 1H), 2.51 (s, 3H); ^13^C NMR (DMSO-*d*_6_, 100 MHz) δ 155.0, 149.3, 149.2, 141.9, 141.7, 135.4, 130.8, 121.8, 121.6, 118.5, 108.9, 107.6, 12.1; *m/z* (ESI^+^) 304.0 ([M+H]^+^, 100%).

**6-Chloro-*N*^4^-(3-methyl-1*H*-indazol-6-yl)pyridine-3,4-diamine (36)**

Nitro compound **35** (303 mg, 2.47 mmol) in IMS (40 mL) was placed in an autoclave. Pd/C (10%, 131 mg, 0.123 mmol) was added and the reaction was heated to 25 °C at 10 bar H_2_ for 3.5 h. The mixture was filtered through Celite® and concentrated *in vacuo*. The residue was purified using flash column chromatography (0% to 100% EtOAc in hexane) to give amine **36** (450 mg, 1.64 mmol, 66%) as an orange solid. ^1^H NMR (DMSO-*d*_6_, 400 MHz) δ 12.35 (s, 1H), 7.82 (s, 1H), 7.50 (dd, *J* = 8.4 Hz, 1H), 7.65 (s, 1H), 7.14 (d, *J* = 1.6 Hz, 1H), 6.94–6.91 (m, 1H), 6.83 (s, 1H), 5.03 (s, 2H), 2.45 (s, 3H); ^13^C NMR (DMSO-*d*_6_, 100 MHz) δ 142.2, 141.6, 140.6, 139.5, 139.1, 135.0, 133.5, 121.3, 119.2, 115.5, 107.1, 100.2, 12.1; *m/z* (ESI^+^) 274.1 ([M+H]^+^, 100%).

**6-Chloro-1-(1-(diethoxymethyl)-3-methyl-1*H*-indazol-6-yl)-1*H*-imidazo[4,5-c]pyridine (37)**

Compound **36** (450 mg, 1.64 mmol) was dissolved in diethoxymethoxyethane (4.87 g, 32.9 mol), formic acid (62 µL, 1.6 mmol) was added and the mixture heated to 100 °C overnight. The mixture was cooled and concentrated *in vacuo*. Saturated aqueous NaHCO_3_ (5 mL) was added, and the resulting precipitate was filtered off, and washed with water and hexane to give the cyclised product **37** (420 mg, 1.09 mmol, 66%) as a light brown solid.^1^H NMR (DMSO-*d*_6_, 400 MHz) δ 8.92 (d, *J* = 0.8 Hz, 1H), 8.83 (s, 1H), 8.00 (d, *J* = 8.4 Hz, 1H), 7.97 (d, *J* = 1.2 Hz, 1H), 7.61 (d, *J* = 0.8 Hz, 1H), 7.50 (dd, *J* = 2.0, 8.4 Hz, 1H), 6.52 (s, 1H), 3.75‒3.68 (m, 2H), 3.60‒3.49 (m, 2H), 2.57 (s, 3H), 1.18‒1.14 (m, 6H); ^13^C NMR (DMSO-*d*_6_, 100 MHz) δ 147.2, 143.8, 142.8, 142.3, 141.2, 141.0, 139.1, 133.8, 124.2, 122.9, 117.8, 107.0, 106.3, 105.9, 62.4, 15.2, 12.0; *m/z* (ESI^+^) 284.1 ([M+H]^+^, 100%).

**3-Methyl-4-[1-(3-methyl-1*H*-indazol-6-yl)imidazo[4,5-*c*]pyridin-6-yl]benzonitrile (OXS008255, 5)**

To a solution of **37** (75 mg, 0.19 mmol) in degassed 1,4-dioxane/water (4:1, 1.0 mL) were added potassium phosphate (121 mg, 0.57 mmol), (4-cyano-2-methylphenyl)boronic acid (37 mg, 0.23 mmol) and Pd(dppf)Cl_2_ (15 mg, 0.020 mmol). The mixture was heated at 100 °C overnight, then diluted with EtOAc, filtered through Celite® and concentrated. To a solution of the resulting material (30 mg) in ethanol (1.2 mL) was added aqueous HCl (2 M, 110 µL, 0.21 mmol) at 0 °C. After stirring for 30 min at room temperature, the mixture was basified with aqueous 2 M NaOH. After a further 30 min, water was added and the mixture was extracted with EtOAc. The organic layer was dried over anhydrous Na_2_SO_4_, evaporated and purified using flash column chromatography (EtOAc) to give OXS008255 **5** (5 mg, 0.01 mmol, 7%). ^1^H NMR (500 MHz, CDCl_3_) δ 9.32 (d, *J* = 1.1 Hz, 1H), 8.33 (s, 1H), 7.93 (d, *J* = 8.4 Hz, 1H), 7.64–7.57 (m, 3H), 7.57–7.50 (m, 2H), 7.32 (dd, *J* = 8.4, 1.8 Hz, 1H), 2.69 (s, 3H), 2.43 (s, 3H); ^13^C NMR (126 MHz, CDCl_3_) δ 152.2, 145.3, 144.5, 144.1, 143.1, 141.1, 140.3, 139.3, 137.8, 134.3, 133.8, 130.7, 129.6, 122.9, 122.6, 118.9, 116.8, 111.8, 106.0, 105.3, 20.4, 12.0; *m/z* (ESI^+^) 365.1 ([M+H]^+^, 100%); HRMS (ESI^+^) C_22_H_17_N_6_^+^ ([M+H]^+^) requires 365.1509; found 365.1512.

**References**

1.	Hapuarachchige, S., Montaño, G., Ramesh, C., Rodriguez, D., Henson, L. H., Williams, C. C., Kadavakkollu, S., Johnson, D. L., Shuster, C. B. & Arterburn, J. B. Design and synthesis of a new class of membrane-permeable triazaborolopyridinium fluorescent probes. *J. Am. Chem. Soc.* **133**, 6780–6790 (2011).

2.	Morcillo, S. P., Leboeuf, D., Bour, C. & Gandon, V. Calcium-Catalyzed Synthesis of Polysubstituted 2-Alkenylfurans from β-Keto Esters Tethered to Propargyl Alcohols. *Chem. - A Eur. J.* **22**, 16974–16978 (2016).

3.	Hayakawa, K., Yodo, M., Ohsuki, S. & Kanematsu, K. Novel Bicycloannulation via Tandem Vinylation and Intramolecular Diels-Alder Reaction of Five-Membered Heterocycles: A New Approach to Construction of Psoralen and Azapsoralen. *J. Am. Chem. Soc.* **106**, 6735–6740 (1984).

4.	Li, Z., Hao, P., Li, L., Tan, C. Y. J., Cheng, X., Chen, G. Y. J., Sze, S. K., Shen, H. M. & Yao, S. Q. Design and synthesis of minimalist terminal alkyne-containing diazirine photo-crosslinkers and their incorporation into kinase inhibitors for cell- and tissue-based proteome profiling. *Angew. Chemie - Int. Ed.* **52**, 8551–8556 (2013).

**6-(4-Methoxyphenyl)-3-(*o*-tolyl)-[1,2,4]triazolo[4,3-*a*]pyridine (OXS000275, 1)**

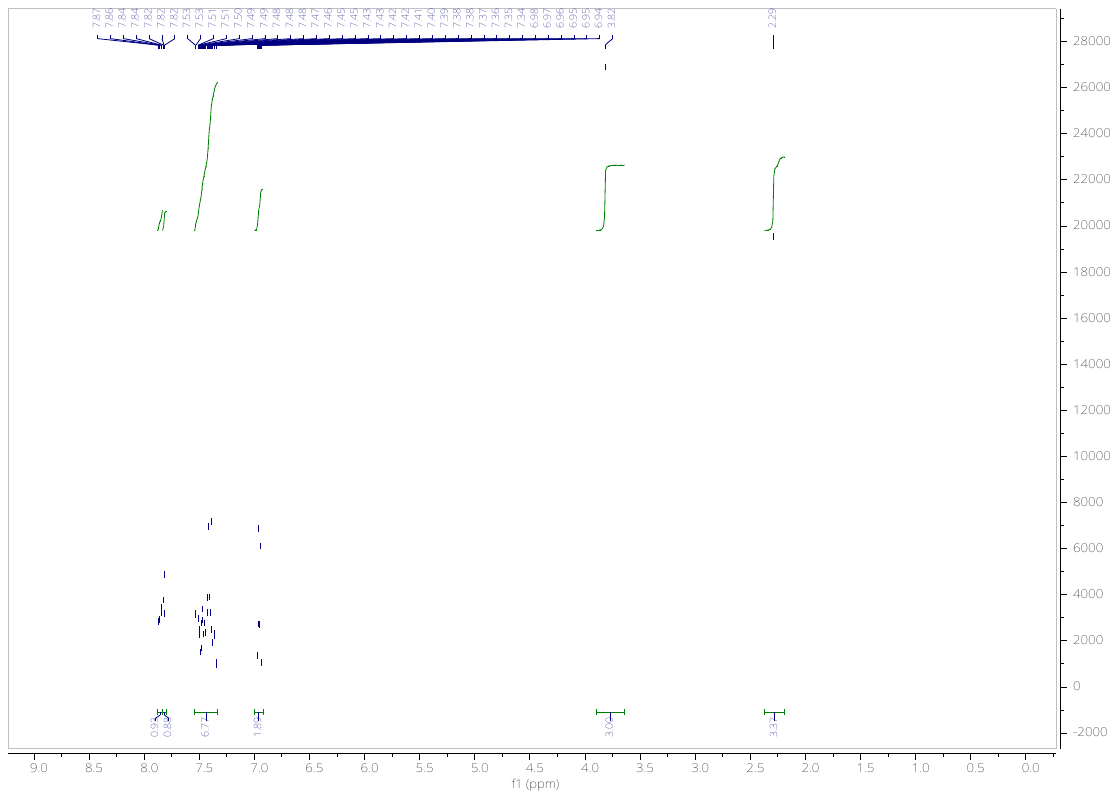

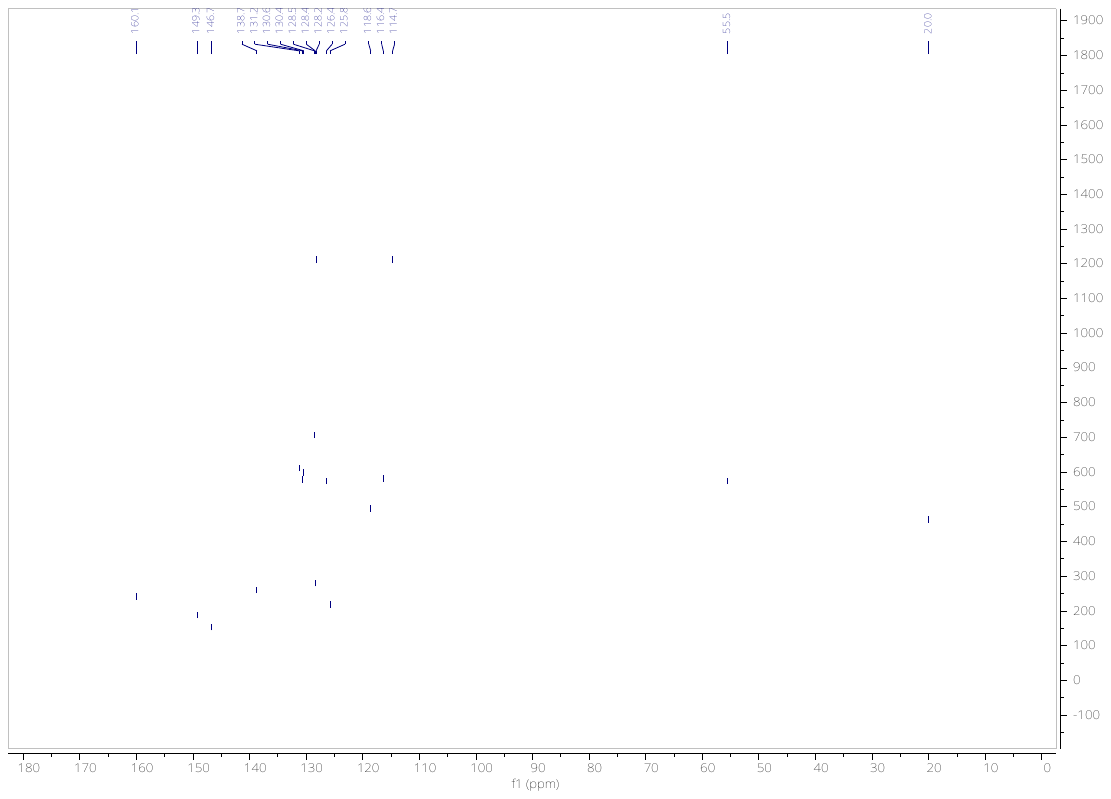

**1-(4-(Difluoromethoxy)phenyl)-6-(4-fluoro-2-methylphenyl)-1*H*-imidazo[4,5-*c*]pyridine (OXS007417, 2)**

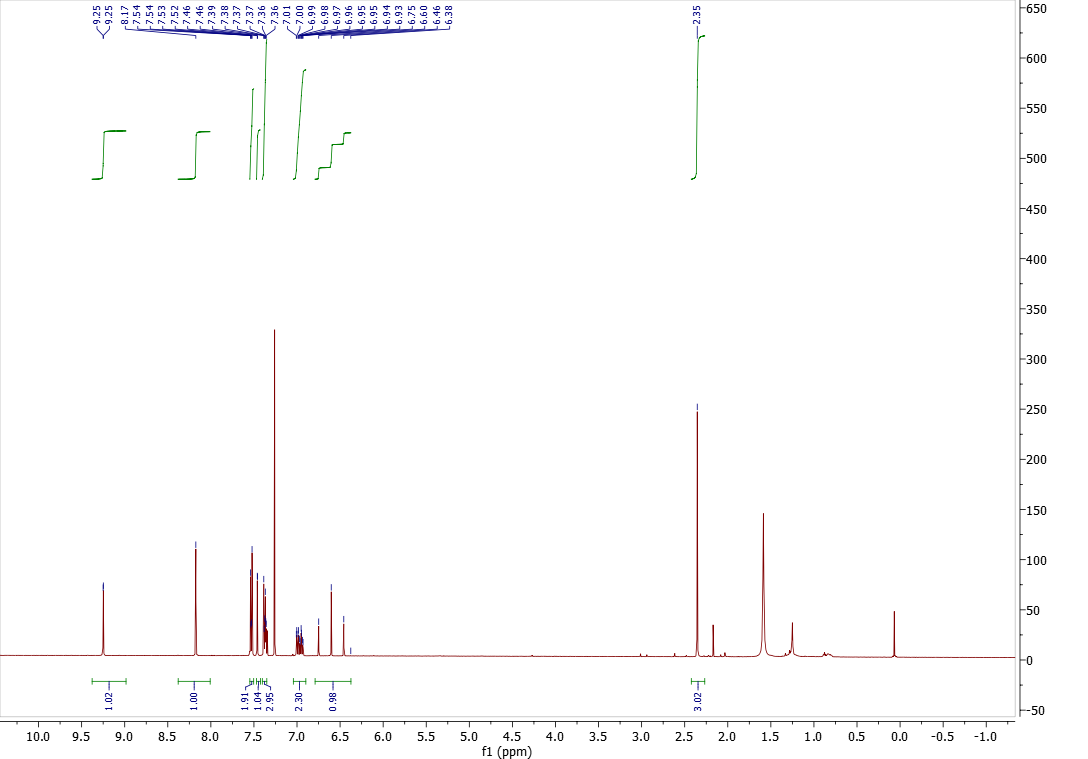

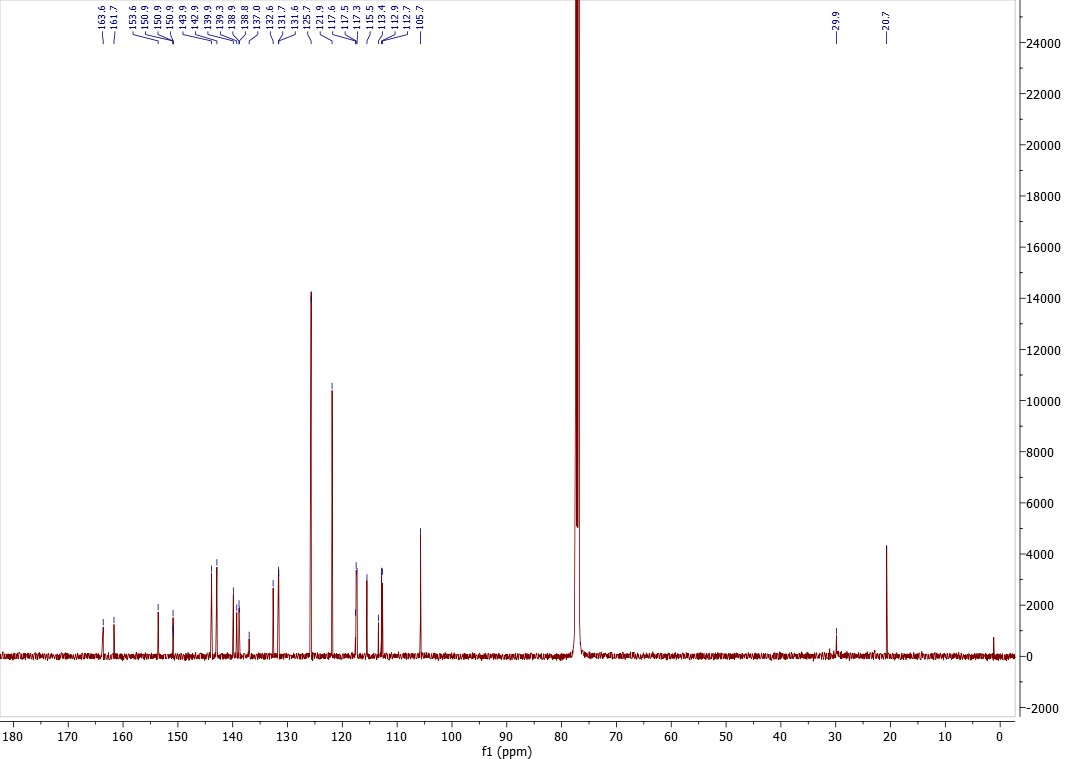

***N*-(6-(4-fluoro-2-methylphenyl)-3-(1-methyl-1*H*-indazol-5-yl)imidazo[1,2-*a*]pyridin-8-yl)methanesulfonamide (OXS007464, 3)**

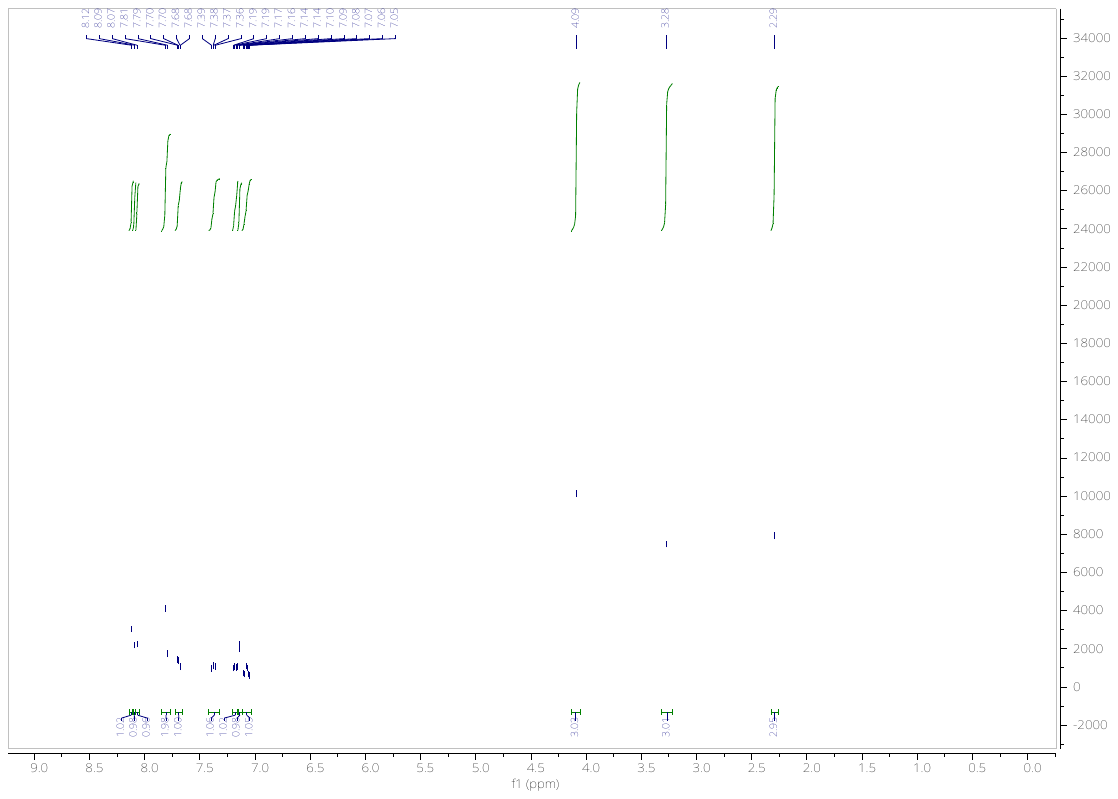

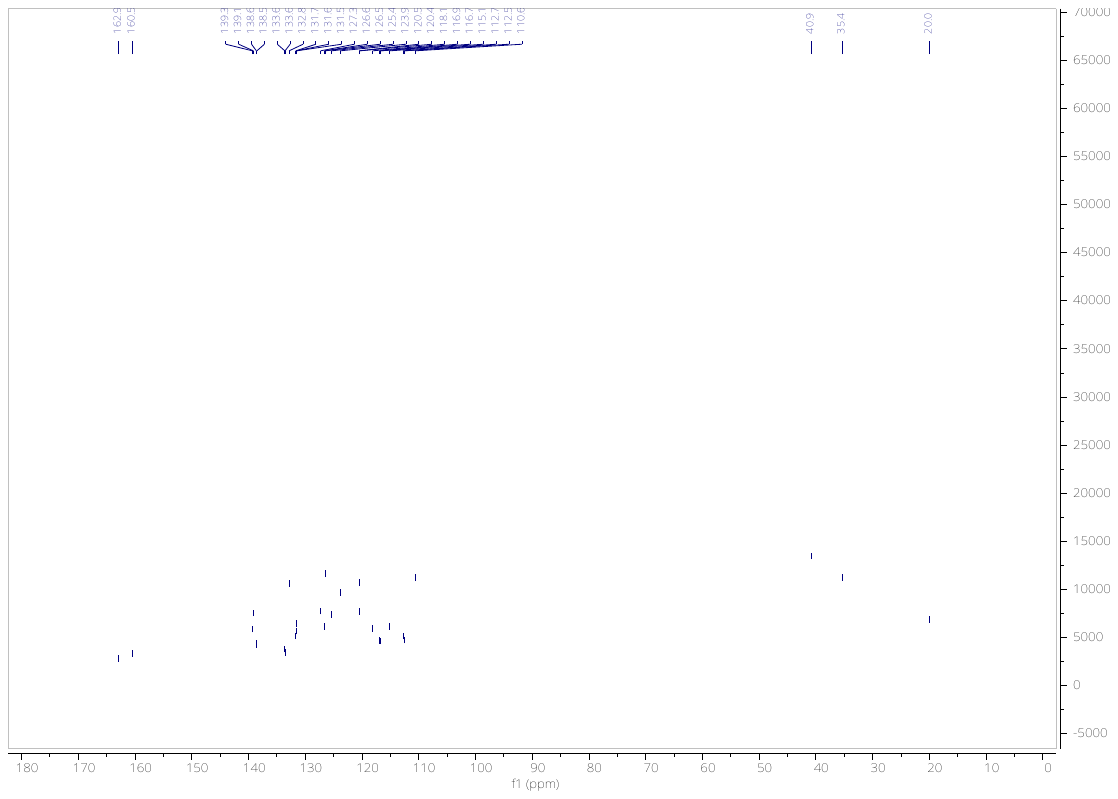

***N*-(3-(1-methyl-1*H*-indazol-5-yl)-6-(2-(2-(3-(prop-2-yn-1-yl)-3*H*-diazirin-3-yl)-ethoxy)phenyl)imidazo[1,2-*a*]pyridin-8-yl)methanesulfonamide (OXS008540, 4)**

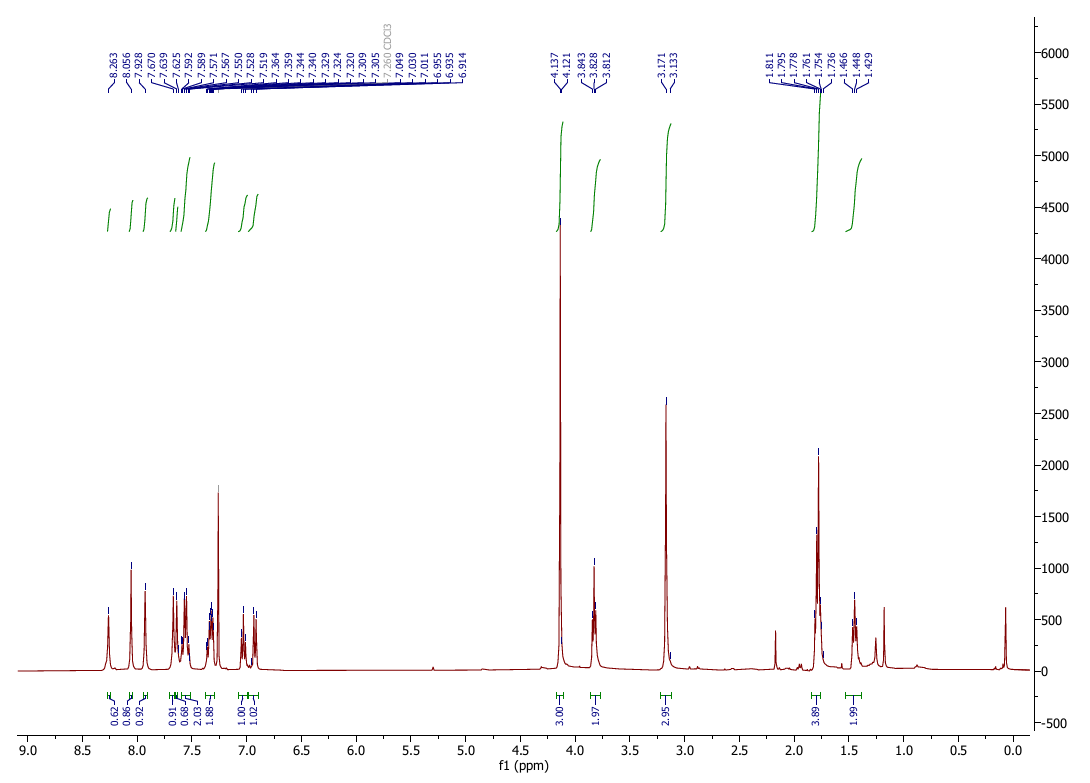

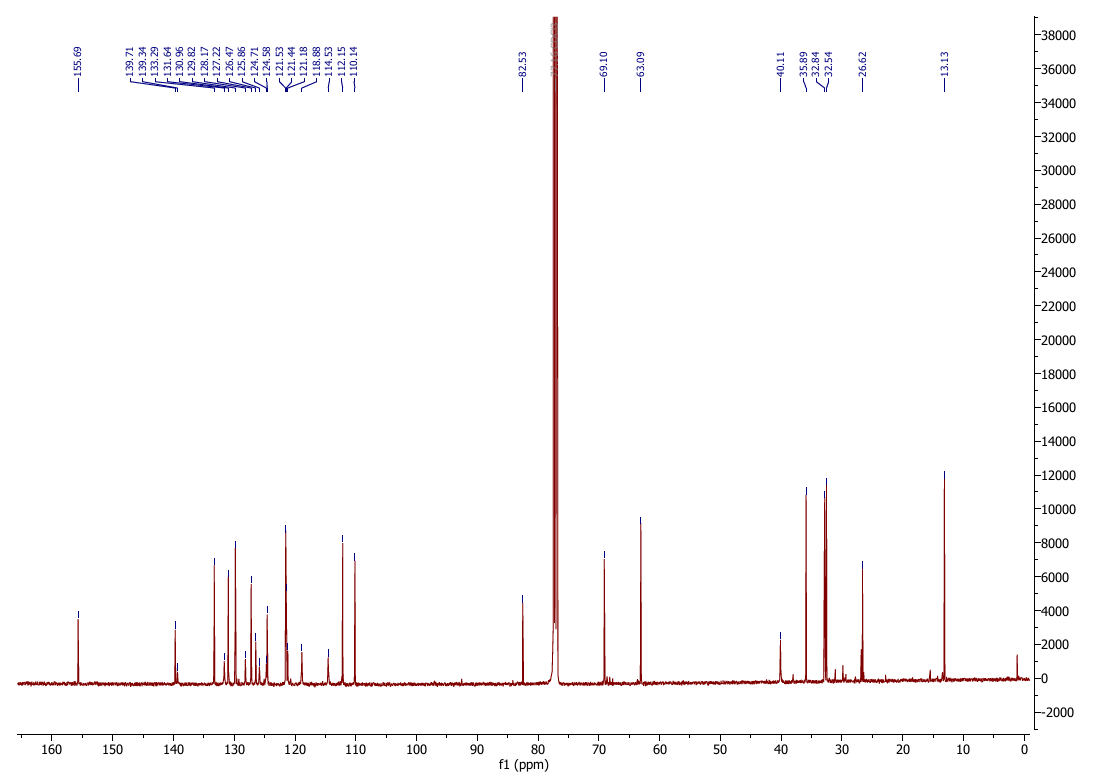

**3-Methyl-4-[1-(3-methyl-1*H*-indazol-6-yl)imidazo[4,5-*c*]pyridin-6-yl]benzonitrile (OXS008255, 5)**

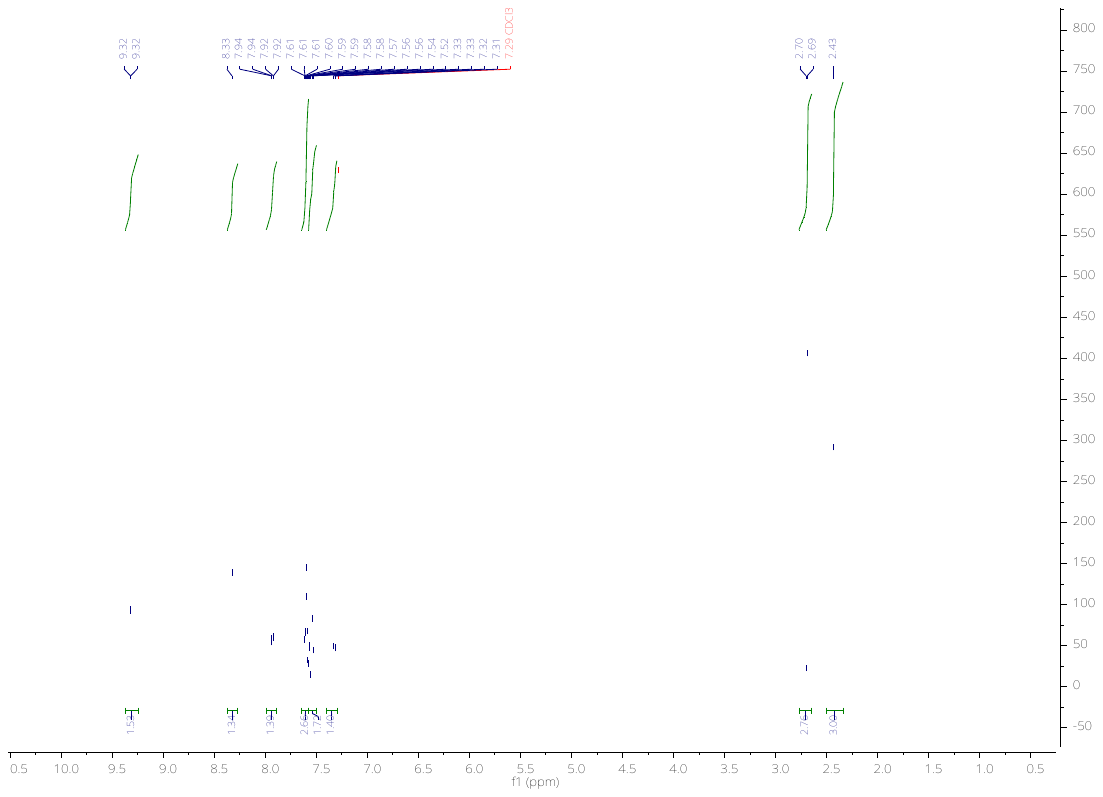

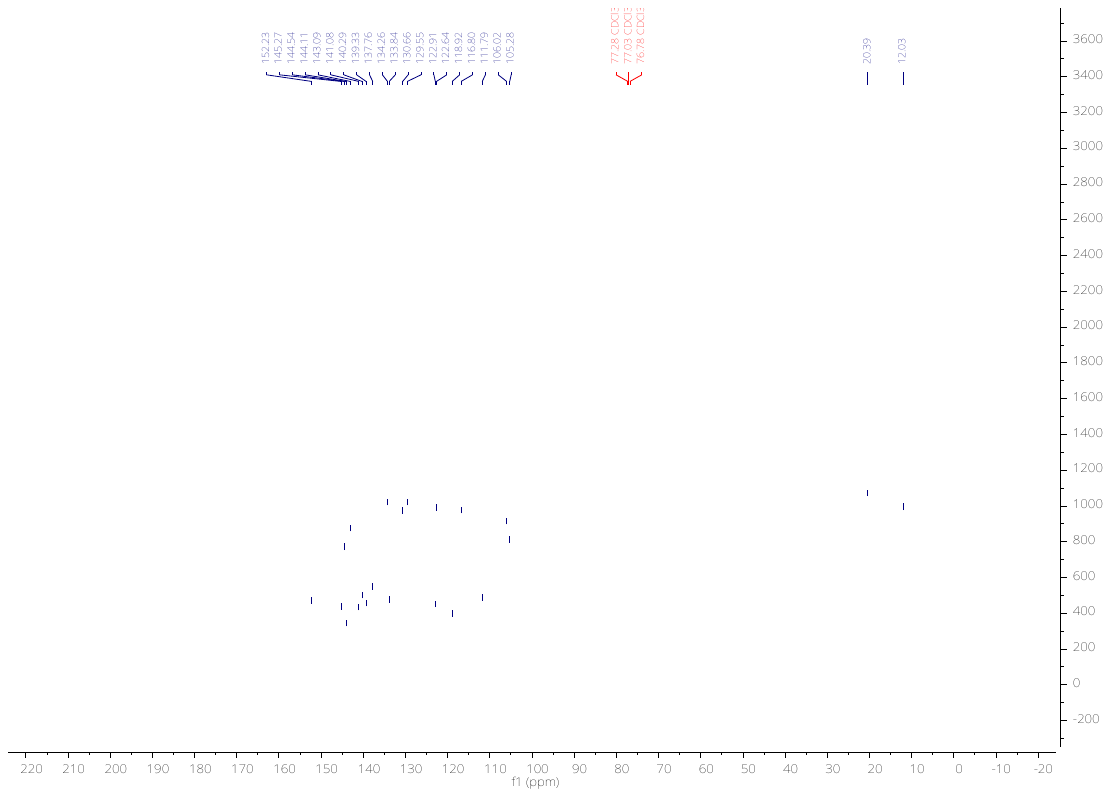

***N*-(6-(4-Fluoro-2-methylphenyl)-3-(1-methyl-1*H*-indazol-6-yl)imidazo[1,2-*a*]pyridin-8-yl)methanesulfonamide (OXS007564, 6)**

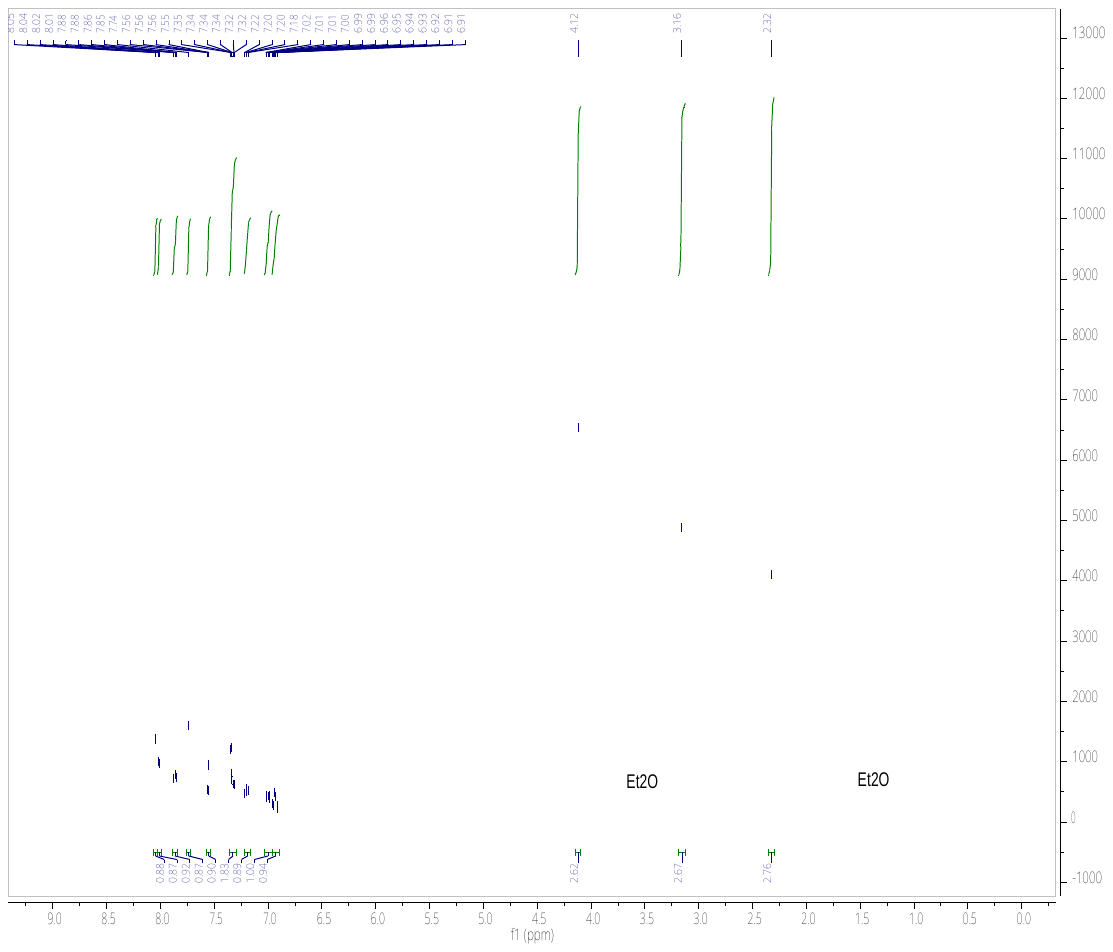

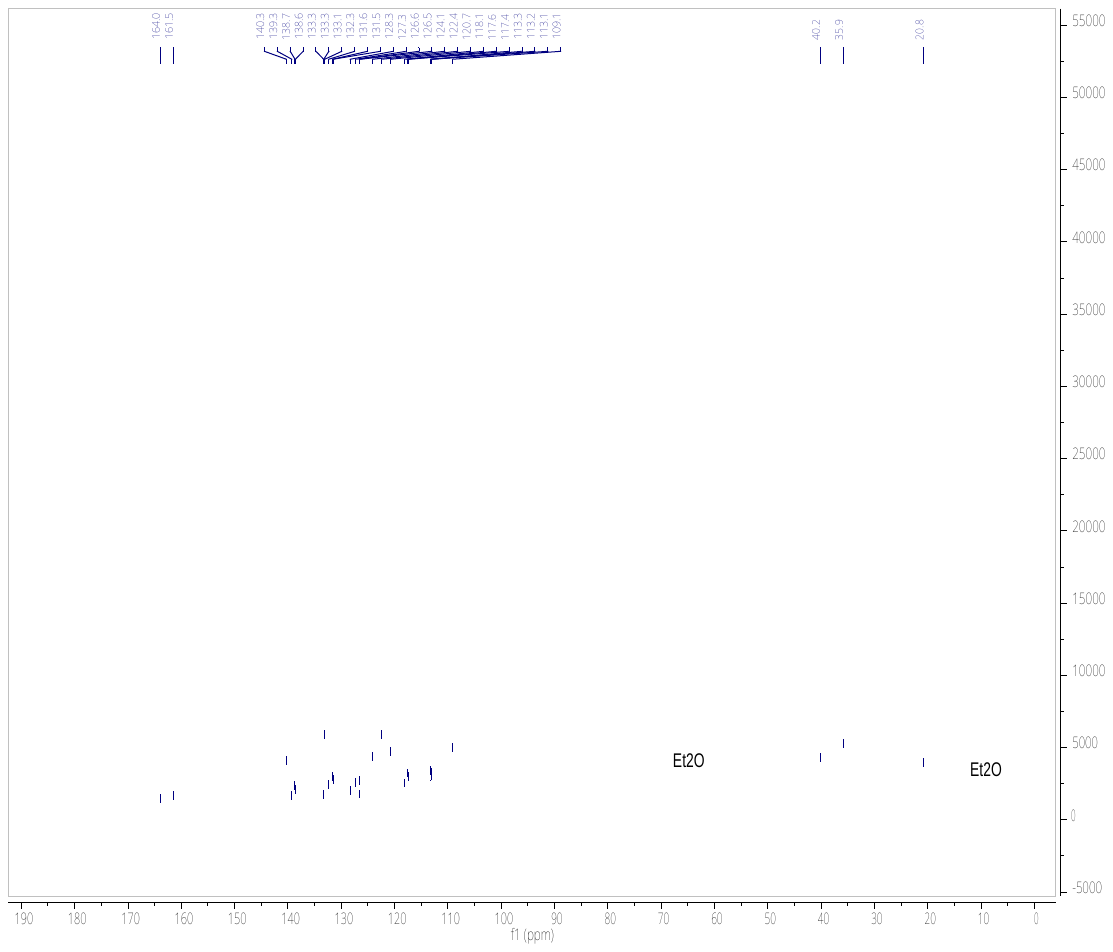
